# Distinct mutational landscapes when comparing germline and somatic cancer variants in forty tumor suppressor genes

**DOI:** 10.64898/2026.02.26.707048

**Authors:** Suhasini D. Lulla, Deborah I Ritter, Chimene Kesserwan, Sharon E. Plon

## Abstract

Germline and somatic cancer variants in tumor suppressor genes (TSGs) share loss of function mechanisms with studies of a few genes (*DICER1*, *CEBPA*) highlighting differences in variant consequence and location. To systematically assess whether TSGs display distinct mutational patterns we leveraged large public genetic databases and compared high-quality pathogenic/likely pathogenic (P/LP) germline variants in ClinVar, with oncogenic/likely oncogenic (O/LO) somatic tumor variants from cBioPortal across 40 TSGs. We obtained 32,941 P/LP germline and 12,907 O/LO somatic variants. Only 3,863 (9.2%) variants were shared. Eighteen TSGs showed significant differences in distributions of variant occurrences by consequence replicated with non-overlapping somatic data from the COSMIC database (chi-squared tests, false discovery rate=5%). *DICER1*, *TP53*, and *SMAD4* displayed excess somatic missense events, while 9 TSGs (e.g., *RB1*, *APC*) contained excess somatic stop-gains throughout the coding sequence. Analysis of tumor type revealed excess stop-gains in tissues exposed to environmental mutagens with corresponding mutation signatures. Germline and somatic events also distributed unevenly across cDNA location with 103 regions of preferential clustering in 39 TSGs (78 somatic, 25 germline). Twenty somatic clusters overlapped recurring frameshifts in homopolymer runs, many in tumors with microsatellite instability. Germline clusters contain more germline-exclusive variants, some driving non-cancer phenotypes reflecting genetic pleiotropy. In some TSGs (*WT1*), germline variants predispose to tumors representing a minority of somatic data. Altogether, germline and somatic variants of TSGs represent unique sets with substantially different patterns shaped by selection pressures including environmental, phenotypic, and functional mechanisms. Characterizing these distinctions enables more accurate clinical interpretation of cancer variants.

## Introduction

The molecular interpretation of the ‘two-hit’ hypothesis proposes that biallelic inactivation of a tumor suppressor gene (TSG) is required for tumor initiation^1–3^. Inactivating events can be coding variants causing loss of gene function (LoF), epigenetic alterations resulting in silencing, or larger genomic events (chromosome loss, large deletions, or recombination events), leading to loss of heterozygosity (LOH)^4,5^. Sporadic tumors involving loss of TSG function result from somatic biallelic inactivating events. Alternatively, having a germline (constitutional) pathogenic LoF variant in a TSG reduces the somatic mutational burden and latency required for oncogenic transformation, thereby conferring inherited cancer susceptibility^6^. Accurate classification of germline and somatic variants in TSGs as being cancer-associated relies on understanding the mutational mechanisms of tumorigenesis.

Cancer-associated variants in TSGs are expected to distribute similarly by variant type and location within the gene, be it in the germline or somatic contexts. In 2018, the Clinical Genome Resource (ClinGen) Germline/Somatic Variant Subcommittee (GSVS) published recommendations that leverage large tumor sequencing datasets, allowing somatic variant hotspots to be used as supporting evidence when classifying the same variant in the germline context as cancer predisposing^7^. However, germline and somatic cancer TSG variants do not always have similar distributions by variant type and location. In *DICER1* associated cancers, a LoF variant can be inherited in the germline or occur sporadically in somatic tissue. But the other variant required to drive tumorigenesis is almost exclusively a somatic missense variant occurring in one of few hotspots in the RNase IIIB domain that impairs RNase IIIB activity without fully inactivating DICER1^8,9^. This expected pattern is incorporated into the ClinGen *DICER1* variant curation expert panel (VCEP) specifications^10^. Similarly, the *CEBPA* TSG has distinct germline and somatic variant locations and mutation types^11,12^. In *CEBPA*-associated acute myeloid leukemias, germline variants are typically small insertions or deletions (indels) causing frameshifts clustered in the N-terminal TAD1 domain of the CEBPA protein. Somatic variants are typically in-frame indels clustering in the C-terminal bZIP domain and impair DNA binding and dimerization without completely eliminating CEBPA function. Knowledge of these differences provide insight into the underlying mechanism of tumorigenesis and have practical applications in informing recommendations for variant classification.

To date, detailed assessment of germline versus somatic cancer variant patterns has not been robustly conducted to identify if other TSGs, in addition to *DICER1* and *CEBPA*, display distinct germline and somatic patterns. Therefore, in this study, we harmonized genetic data from large public germline and somatic variant databases and built a custom pipeline to systematically characterize variant distributions for 40 TSGs having substantial data. We compared germline and somatic cancer variant distributions by 1) identity, 2) variant molecular consequence, 3) underlying cancer tissue type, and 4) location within the genes. Our findings provide a comprehensive landscape of cancer genetic variants in 40 TSGs and demonstrate substantial differences in germline and somatic mutational mechanisms.

## Methods

### Gene list curation

We selected TSGs with well-characterized autosomal dominant germline cancer predisposition for our analyses (Figure 1A). We downloaded 102 cancer predisposition gene annotations with germline mutations from the Catalogue of Somatic Mutations in Cancer (COSMIC) Cancer Gene Census (August 2022)^13^. We selected 56 tumor suppressor genes from this list whose HUGO gene symbol^14^ overlapped with TSGene2.0, a database of 1217 TSGs (August 2022)^15^. Genes were annotated with associated phenotypes and inheritance patterns using Online Mendelian Inheritance in Man (OMIM)^16^. We excluded genes lacking autosomal dominant inheritance of cancer predisposition (n=13), as well as genes lacking somatic variant classification in OncoKB^17^ (November 2022, n=3) and those with insufficient data (<25 germline or somatic events, n=3). *CEBPA*, *MSH6*, and *SMARCA4* were added to the list, being well-documented cancer predisposing TSGs that otherwise met our criteria but were present in only one dataset. Our resulting list consisted of 40 TSGs (Table 1).

**Figure 1:**
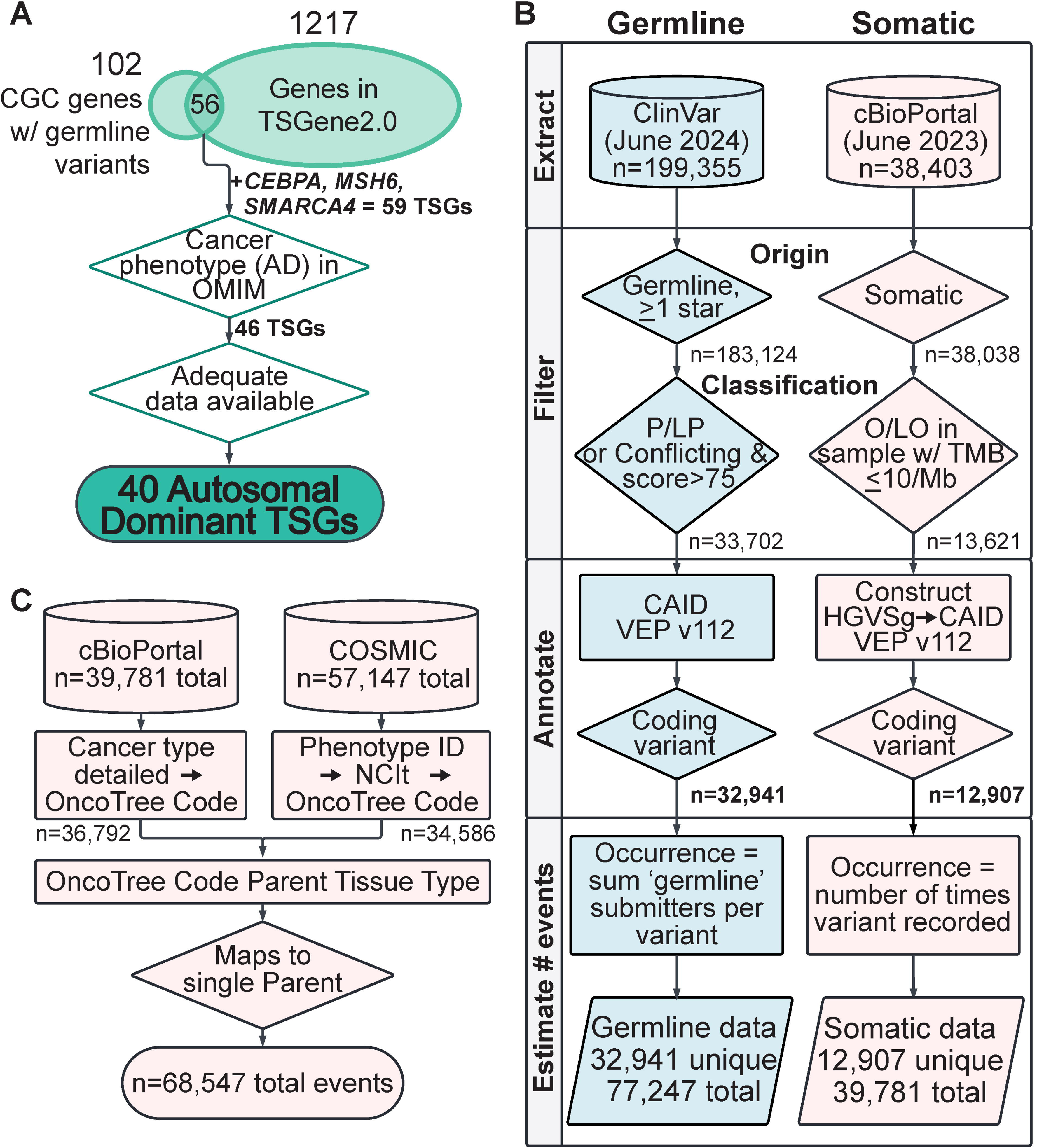
Tumor Suppressor Gene (TSG) list compilation and pipeline to extract germline and somatic cancer variants. (A) Selected 56 TSGs from the Cancer Gene Census genes with germline variants and added *CEBPA*, *MSH6*, and *SMARCA4*. Forty-six TSGs associated with cancer phenotypes in OMIM were retained, but 6 TSGs with inadequate germline or somatic data were removed, resulting in a list of 40 TSGs to analyze. (B) Extracting and processing germline variants from ClinVar and somatic variants from cBioPortal. ClinVar data from 40 TSGs was filtered to 32,941 unique P/LP or high scoring conflicting variants with at least 1 germline submission and 1 or more stars for the assertion provided. estimated. Somatic data from cBioPortal was filtered to 12,907 unique O/LO variants in samples with TMB≤10/Mb. Germline and somatic variants were annotated with their CAID and molecular consequence using VEP (v112). Total occurrence per variant was estimated resulting in 77,247 germline events and 39,781 somatic events. (C) Parent tissue type assignment of somatic variants using the OncoTree ontology. COSMIC Phenotype IDs were mapped to OncoTree codes through the NCIt ontology. Combined somatic data from cBioPortal and COSMIC totals 68,547 variant occurrences that mapped to a single parent tissue type. Charts created in Lucid (lucid.co).  TSG, tumor suppressor gene; AD, autosomal dominant; OMIM, Online Mendelian Inheritance in Man^16^; P/LP, pathogenic/likely pathogenic; O/LO, oncogenic/likely oncogenic; TMB, tumor mutation burden; CAID, ClinGen Allele Registry ID^30^; VEP, Variant Effect Predictor^32^; COSMIC, Catalogue of Somatic Mutations in Cancer; NCIt, National Cancer Institute Thesaurus.

**Table 1.**
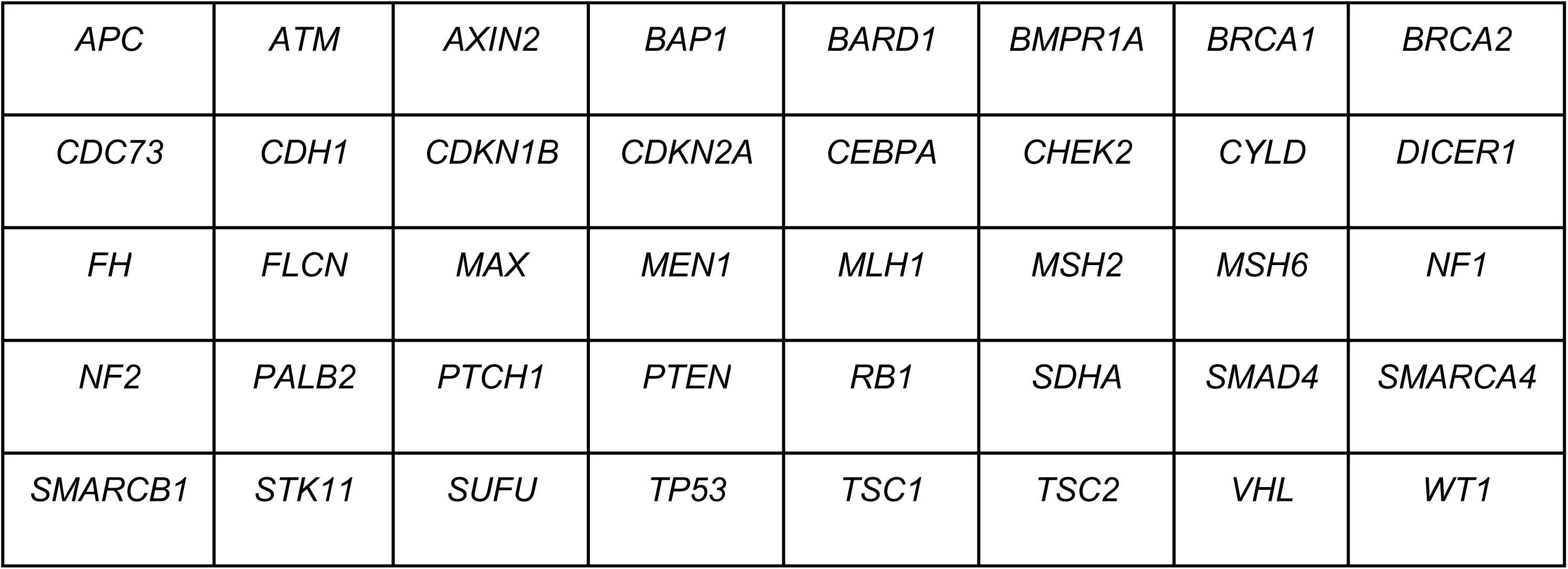
List of 40 tumor suppressor genes in this study.

|  |  |  |  |  |  |  |  |
| --- | --- | --- | --- | --- | --- | --- | --- |
| *APC* | *ATM* | *AXIN2* | *BAP1* | *BARD1* | *BMPR1A* | *BRCA1* | *BRCA2* |
| *CDC73* | *CDH1* | *CDKN1B* | *CDKN2A* | *CEBPA* | *CHEK2* | *CYLD* | *DICER1* |
| *FH* | *FLCN* | *MAX* | *MEN1* | *MLH1* | *MSH2* | *MSH6* | *NF1* |
| *NF2* | *PALB2* | *PTCH1* | *PTEN* | *RB1* | *SDHA* | *SMAD4* | *SMARCA4* |
| *SMARCB1* | *STK11* | *SUFU* | *TP53* | *TSC1* | *TSC2* | *VHL* | *WT1* |

### Germline variant data extraction and filtering

Figure 1B provides the bioinformatic pipeline used to extract and annotate the TSG data. We extracted germline variant data for the 40 TSGs from ClinVar (https://www.ncbi.nlm.nih.gov/clinvar/)^18^, a publicly available variant repository with >3 million unique variation records that include germline pathogenicity and somatic oncogenicity classifications when provided by the submitter. Germline classifications are based on the American College of Medical Genetics and Genomics and the Association for Molecular Pathology (ACMG/AMP) guidelines (benign (B), likely benign (LB), uncertain significance (VUS), likely pathogenic (LP), pathogenic (P))^19^. We used Entrez Direct^20^ to extract 201,061 unique variants in 40 TSGs by Gene ID (June 2024, see Data and code availability). We limited data to single-gene variants by removing ‘Variation type’ of ‘copy number loss’, ‘copy number gain’, ‘Complex’, ‘fusion’, ‘Haplotype’, ‘Translocation’, and ‘Variation’, leaving 199,355 unique variants. We excluded the following variant entries: lacking a standard HGVS name, unknown breakpoints (for example: NC_000017.11:g.(?_31095304)_(31374161_?)del), and ‘Review Status’ of ‘no assertion criteria provided’, ‘no assertion provided’, ‘no assertion for the individual variant’, ‘no classification provided’, or ‘no classification for the individual variant.’ To include only germline variant classifications, we filtered by ‘Origin’ set to one of the following terms: ‘germline’, ‘de novo’, ‘maternal’, ‘paternal’, ‘inherited’, ‘uniparental’, or ‘biparental’.

Our focus is on clinically relevant germline variants that result in cancer predisposition; hence, we initially included only those variants that have clinical assertions of ‘Pathogenic’, ‘Likely pathogenic’, or ‘Pathogenic/likely pathogenic’ (P/LP).

To include variants with conflicting classifications but having a majority of pathogenic submissions, we developed a scoring metric by assigning each ACMG/AMP assertion a value representing the midpoint of the classification’s posterior probability e.g. Likely Pathogenic = 0.95 multiplied by 100, (Table S1)^21^. We calculated ‘conflict scores’ as the average value across all germline submissions for the variant and included in our dataset variants that scored ≥ 75 (Figure S1). This threshold was selected since it exceeds the conflict score for a variant with at least one more P/LP submission than VUS. We validated this threshold by calculating the conflict scores of variants that were previously conflicting and resolved by ClinGen VCEPs for *DICER1* and *CDH1* (Figure S2, Table S2)^10,22^. In total, 544 out of 19,341 conflicting variants in 40 TSGs were added to the final germline set. The resulting P/LP dataset consisted of 33,702 unique germline cancer variants^23,24^ on the GRCh38 reference human genome assembly (Figure 1B).

### Somatic variant data extraction and filtering

Somatic cancer variant data for the 40 TSGs was extracted from cBioPortal, a publicly available repository of genetic variation in tumors^25,26^, including large-scale tumor sequencing studies such as The Cancer Genome Atlas (TCGA). We used the cbioportalR package (https://github.com/karissawhiting/cbioportalR) to extract variant information by HUGO gene symbol from 69,276 samples in 213 non-redundant studies (Table S3) as of May 2023. To filter for somatic variants, we excluded variants having ‘mutationStatus’ of ‘uncalled’ or ‘germline’ and variants having ‘tumorAltCount’ = 0. Duplicate occurrences of variants per patient ID were dropped, resulting in 38,038 unique somatic variants in 40 TSGs on the GRCh37 reference human genome assembly. To avoid over-representation of variants from hyper-mutated tumors, we excluded samples with tumor mutation burden (TMB) greater than ten per mega base^27^. Variants were then annotated with oncogenicity classifications using OncoKB’s application program interface (API)^17,28^. We included 13,621 unique variants classified as ‘Oncogenic’ or ‘Likely Oncogenic’ (O/LO), since these are cancer-associated variants relevant to our analyses.

For replication analyses, we similarly processed and analyzed somatic data from COSMIC^29^ that was not already present in cBioPortal by removing samples if the ‘SAMPLE_NAME’ matched an ID in the ‘sampleId’ column of cBioPortal data (Figure S4A). We downloaded somatic variant data in COSMIC’s Census Genes Mutations (Cosmic_MutantCensus_Tsv_v98_GRCh38.tar, September 2023) and matched variants with patient information in the ‘Samples’ and ‘Classification’ downloads for the same 40 TSGs. We dropped duplicate occurrences of variants per individual in the remaining data from COSMIC. Variants were annotated with oncogenicity classifications using OncoKB’s API and then filtered to include the 15,909 unique ‘Oncogenic’ and ‘Likely Oncogenic’ variants in COSMIC having defined HGVS nomenclature on the GRCh38 reference human genome assembly.

### Harmonizing and comparing unique germline and somatic variants

cBioPortal somatic variants did not use HGVS nomenclature and were annotated in the GRCh37 reference human genome assembly. We constructed genomic HGVS nomenclature and used the ClinGen Allele Registry^30^ API to annotate variants with their cDNA HGVS on the GRCh38 assembly^31^. We used the Matched Annotation from NCBI and EMBL-EBI (MANE) Select transcript for 39 TSGs and MANE Plus Clinical transcript for 1 TSG (*SMARCA4*) since these represent the canonical transcripts for each gene on which somatic variant data was curated in OncoKB (Table S4). Germline and somatic variants were also annotated with their ClinGen Allele Registry ID (CAID), a standardized identifier unique to every genomic change, which was used to evaluate the proportion of unique variants that were present in both, germline and somatic sets. The Venn Diagram Generator tool (http://barc.wi.mit.edu/tools/venn/) was used to illustrate the true to size extent of overlap of variants.

### Annotating germline and somatic variants

We used the Variant Effect Predictor (VEP v112)^32^ tool from Ensembl to annotate predicted molecular consequence and location on the MANE transcript of germline and somatic variants. We collapsed molecular consequence categories to the first, most severe consequence prediction by VEP. We maintained VEP’s default severity ordering except for the ‘stop-gained, frameshift_variant’ annotations (288 P/LP germline variants and 145 O/LO somatic variants in total), which were included in the ‘frameshift_variant’ category. Variants spanning more than 50 bp (575 P/LP germline variants and 77 O/LO somatic variants in total) were annotated as ‘>=50bp_indel’.

All non-coding variant categories (Table S5) and synonymous variants were removed from further analyses since cBioPortal only contains oncogenic curations for coding variants (except the *TERT* promoter non-coding variants). To measure the overlap of germline and somatic variant datasets by protein change, we used the VEP protein HGVS annotations.

### Estimating the occurrence of germline and somatic events

Somatic variants are recorded in cBioPortal and COSMIC as many times as they are observed in tumors. This count served as our estimate of total somatic events for each variant (which ranged from 1 to 894 events per variant). In contrast, ClinVar records rarely contain information on the number of individuals with the observed germline variant. Instead, we used the number of germline origin record submitters per variant as a proxy for total germline events (which ranged from 1 to 65 submitters per variant).

### Distribution of germline and somatic events by molecular consequence

To compare the distribution of total germline and somatic events per gene by molecular consequence, we constructed contingency tables of germline and somatic event counts by VEP annotated category per TSG. There can be more than one somatic event in the same TSG per patient, so to account for independent observations, we selected one variant at random per patient. We repeated random selections five times and entered the average somatic event count per category in the contingency table. We removed molecular consequence categories containing less than 1% of both germline and somatic event counts per gene and excluded categories with less than 5 expected germline and somatic event counts if both were sampled from the same distribution. See the statistical analyses section for distribution comparison tests.

### Distribution of somatic events subset by tumor tissue types

We combined non-overlapping somatic cancer data from cBioPortal and COSMIC. O/LO somatic events from cBioPortal were matched with ‘Cancer type detailed’ assigned to the tumor sample they were detected in. This string represents the OncoTree terminology of the specific tumor type^33^. We used the OncoTree ontology to annotate parent tissue types for each event using the OncoTree ontology file (https://oncotree.mskcc.org:443/api/tumor_types.txt?version=oncotree_latest_stable, downloaded April 2024). We excluded 3,091 events whose ‘Cancer type detailed’ mapped to more than one parent tissue type in the OncoTree ontology, leaving 36,792 O/LO events with tissue type annotations.

Of the 57,147 O/LO events from COSMIC not in cBioPortal, we assigned an OncoTree parent tissue type to 34,586 events by converting National Cancer Institute Thesaurus (NCIt) codes^34^ in COSMIC to OncoTree ontology using the publicly available Ontology to Ontology Mapping Tool (https://github.com/cBioPortal/oncotree/blob/April_2025/docs/OncoTree-Mapping-Tool.md). Combining these sets from cBioPortal and COSMIC resulted in a dataset of 68,547 O/LO somatic events with parent tissue type annotations (Figure 1C). To compare the distribution of events by molecular consequence across tissue types, we subset the O/LO events by parent tissue type and constructed contingency tables per TSG. Each table comprised stop-gain and frameshift event counts from germline data and each somatic tissue type with available data (>25 total events in that tissue type). See statistical analyses for statistical tests used to compare this distribution.

### Mutation signature assignment of somatic events subset by tumor tissue types

We analyzed tumor tissue specific mutation signatures contributing to the TSG somatic mutational landscape by including all somatic events in 40 TSGs in cBioPortal data, unfiltered for oncogenicity annotations. The cumulative somatic single base substitution (SBS) profile for this data was subset by tissue type and assessed for the following tumor tissue types: skin, lung, bowel, bladder, and myeloid. We used COSMIC’s SigProfiler Assignment tool with the ‘Targeted sequencing’ option for a size of 120 kilobases covering the 40 TSGs to identify known mutation signatures contributing to these cumulative SBS profiles.

### Preferential clustering of germline and somatic events

To identify differences in distribution of germline and somatic events by location, we annotated variants using the coding DNA HGVS position on the MANE Select transcript. For frameshift variants, the position at which the genomic change begins was used and for splice site variants in intronic regions, the closest coding sequence position was used. Variants greater than 50 bp were excluded. We calculated the difference in proportions of somatic and germline events along the cDNA of each TSG using an optimized window size of 50bp moving 1bp at a time (method adapted from the calculation of oncology missense tolerance ratios^35^).

Value recorded at every window:

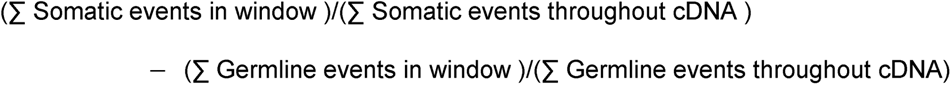

Preferential clustering was defined as regions where the window difference was greater than 2.5 standard deviations (SD) away from 0 (where 0 represents the value if germline and somatic events displayed the same distribution in the 50bp window). Preferential clusters were extended to include neighboring windows if the linked windows exceed the 2.5 SD threshold with the same sign (both positive becoming a single somatic cluster, or both negative being a single germline cluster). We plotted moving difference in fraction curves and 1-dimensional heatmaps identifying preferential clusters for each TSG using matplotlib and seaborn (see codes in data availability)^36,37^.

We analyzed the proportions of variant molecular consequence categories in each cluster and defined a predominant category driving clustering as one that represented >80% events in the cluster. To determine whether somatic predominantly missense clusters aligned with known missense hotspots, we searched cancerhotspots.org^38,39^ for recurring somatic missense variants. For genes with recurring variants in somatic predominantly frameshift clusters, we retrieved microsatellite instability status (MSI Status) from cBioPortal and coding sequences using the consensus coding sequence (CCDS) database^40^ and identified the sequences flanking clusters as well as all homopolymer runs greater than or equal to 5bp throughout the coding sequences of that TSG. We used the nonsense mediated decay (NMD) plugin for VEP to determine whether variants within germline and somatic clusters are predicted to escape NMD.

### Statistical Analyses

Chi-squared tests were performed on contingency tables prepared to compare the distribution of events by molecular consequence using chi2_contingency from SciPy^41^. Fisher’s exact tests were performed if a table did not satisfy the assumptions of the chi-squared test. Significant differences were identified at a false discovery rate of 0.05 after adjusting p-values for multiple comparisons using Bonferroni correction. To identify the categories driving significance, we calculated the adjusted standardized Pearson’s residuals (ASPR)^42^ for each table. We used the pheatmap library in R (see codes in data availability) to plot the germline ASPR across all significant TSGs for those categories where at least one TSG had an ASPR≥3. Similarly, we plotted stop-gain ASPRs for all significant TSGs when comparing stop-gain and frameshift distributions across tissue types. The heatmap includes those tissue types where at least one TSG had an ASPR≥3. Distributions of exclusive events were compared between germline and somatic preferential clusters using a Mann Whitney non-parametric test at a false discovery rate of 0.05 (performed using GraphPad Prism version 10.4.1 for macOS, GraphPad Software, Boston, Massachusetts USA, www.graphpad.com).

## Results

### Limited overlap of pathogenic germline and oncogenic somatic variants in 40 TSGs

To investigate whether the germline and somatic variant landscapes are congruent, we assessed the extent of overlap of unique variants. The extracted, filtered, and harmonized dataset described in Methods comprised 32,941 unique P/LP germline coding variants from ClinVar and 12,907 unique O/LO somatic coding variants in tumor specimens from cBioPortal (Figure 1B) in the 40 selected TSGs with sufficient data. Only 3,863 (9.2% of 41,985 total unique variants by identity) were found in both sets (Figure 2A). Germline and somatic variants were classified using distinct classification frameworks; thus, we evaluated whether the lack of variant overlap was due to discordant classification. However, only 4,491 O/LO somatic variants from cBioPortal were found in the entire ClinVar germline dataset unfiltered for pathogenicity (n=183,124 variants) (Figure 2A), with 86% of those classified as P/LP in ClinVar. Only 4.7% were conflicting variants not meeting our P/LP threshold (Methods), 8.9% were classified as VUS, and <1% as LB/B (Figure 2B). These results were replicated using somatic data from COSMIC absent in cBioPortal (Figure S4). Thus, the germline P/LP and somatic O/LO classification criteria are overwhelmingly concordant, contributing little to the limited overlap of variant data.

**Figure 2:**
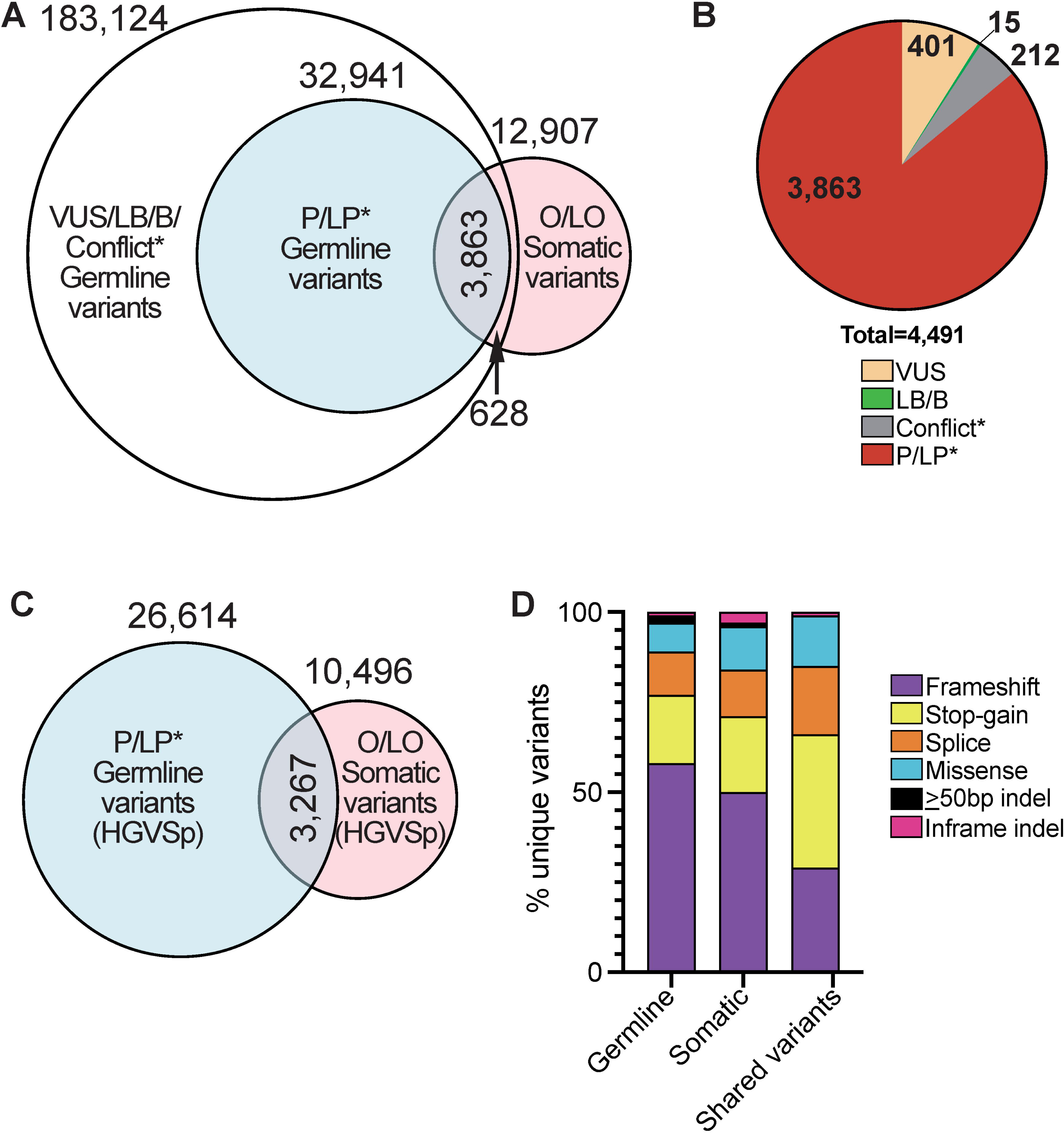
Shared unique P/LP germline and O/LO somatic variants. (A) DNA changes shared between O/LO somatic variants and germline variants. P/LP* in inner blue circle, and germline data unfiltered for pathogenicity in outer circle, containing in addition to 3863 shared P/LP* variants, only 628 VUS, LB/B, and conflict* variants in ClinVar. (B) Classification in ClinVar of shared germline and O/LO somatic variants depicted in (A). (C) Shared O/LO variants and P/LP germline variants by protein change (HGVSp). (D) Predicted molecular consequence distributions of unique P/LP variants, unique O/LO variants, and 3,863 shared P/LP and O/LO variants. P/LP*=includes P/LP and conflicting variants with conflict scores ≥75. Conflict*=conflicting variants with conflict scores <75. P/LP, pathogenic/likely pathogenic; O/LO, oncogenic/likely oncogenic; VUS, variants of uncertain significance; LB/B, likely benign/benign.

Multiple unique genomic changes can lead to the same protein change; hence, we also compared the datasets by predicted protein change (HGVSp) on the MANE transcript. There was little increase in the proportion of overlapping variants with only 3,267 of 33,843 (9.7%) unique protein changes occurring in both sets (Figure 2C).

Variants that overlapped by identity between germline and somatic sets (n=3,863) were distributed across the following mutation types: nonsense variants (37%), frameshift (29%), splice site variants (19%), and missense variants (14%) (Figure 2D). Separately, frameshift variants constituted the greatest component of germline P/LP and somatic O/LO variant sets (19,218 germline (58%) and 6,446 somatic (50%) variants), but only 1,121 (4.6% of total frameshift) were shared, partly explaining the limited overlap between datasets. Despite composing a minority of germline and somatic sets, variants resulting from single nucleotide variation (SNV) including missense, nonsense, and splice site variants, had a larger extent of overlap (n=2,712 SNVs) but still limited to only 16.9% of total SNVs (Figure 2D). Although our analytic pipeline was designed to extract cancer-associated variants with similar TSG inactivating functional consequences, this minimal overlap of unique variants was the first indication of distinct germline and somatic mutational patterns across well-characterized TSGs.

### Germline and somatic variants are distributed distinctly by molecular consequence in 18 TSGs

We assessed differences between germline and somatic molecular consequence distributions by estimating the total occurrence of each unique variant (see Methods). This ranges from 1 to 65 total germline events (using the proxy of ClinVar submitters) per P/LP germline variant and from 1 to 894 tumor events per O/LO somatic variant (Figure S3). Our dataset had a median of 796 P/LP germline events (interquartile range (IQR) = 298-1965 events) and a median of 200 O/LO somatic events (IQR = 91-626 events) per gene. However, the number of unique and total events per gene differs dramatically across the 40 TSGs (Figure 3) The largest numbers of germline events per gene were observed in *BRCA1* (n=13,487 events) and *BRCA2* (n=17,104 events), which are longstanding, established hereditary cancer associated genes frequently tested in the germline setting. On the somatic side, *TP53* had the maximum number of O/LO events (n=19,781) and other genes, e.g. *CYLD* had many fewer germline and somatic events. Given the wide range and dissimilar distributions, all subsequent analyses were performed separately for each gene.

**Figure 3:**
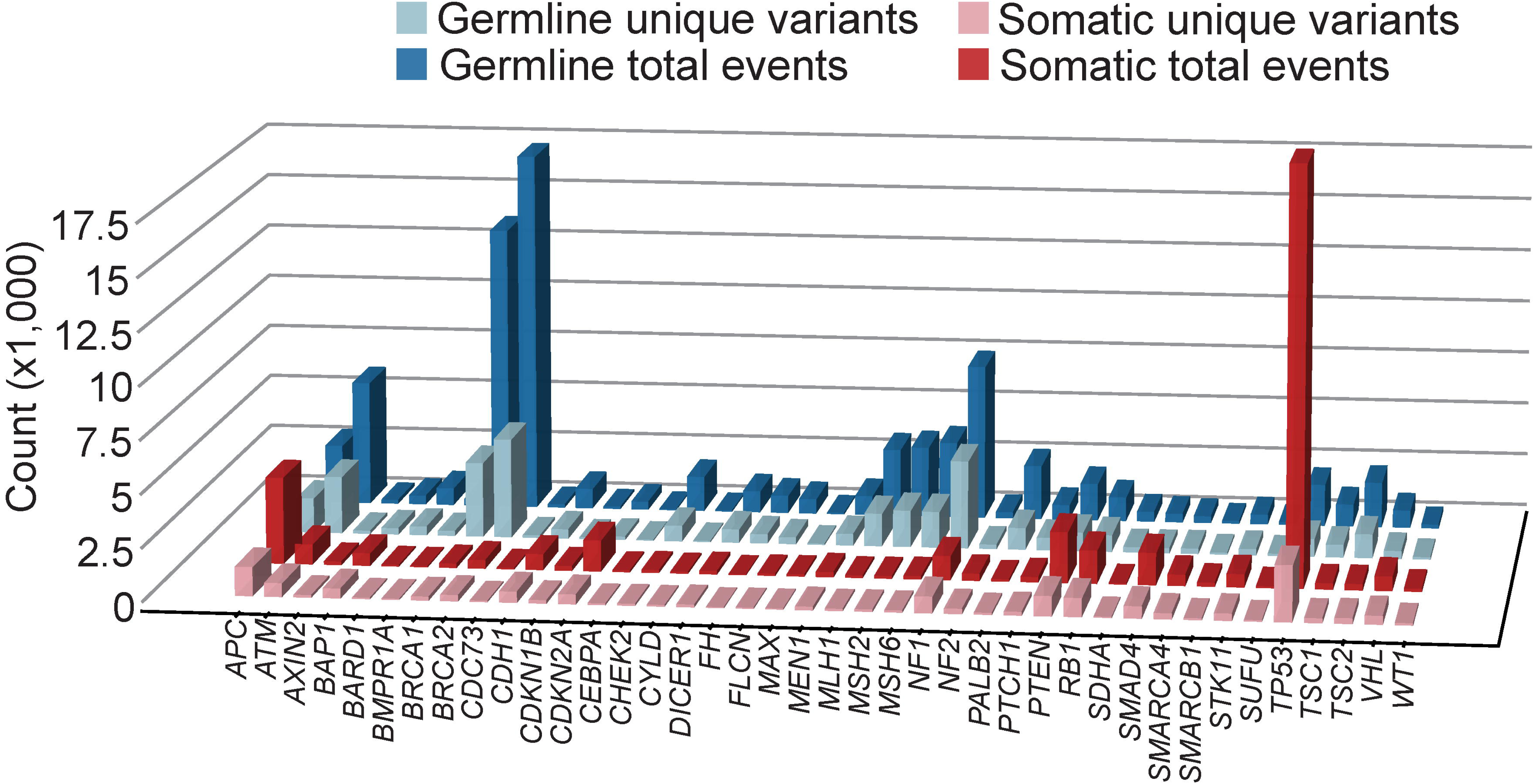
Total P/LP germline and O/LO somatic events in TSGs distribute unevenly across 40 TSGs. (For P/LP germline events, median = 796, interquartile range (IQR) = 298-1965 events, and for O/LO somatic events, median = 200, IQR = 91-626 events per gene). P/LP, pathogenic/likely pathogenic; O/LO, oncogenic/likely oncogenic.

We analyzed the events by molecular consequence per the VEP predictor and identified 21 TSGs with significantly different distributions using chi-squared or Fisher’s exact tests (Bonferroni adjusted p < 0.05, Table S6, Methods). For a replicate analysis, we compared P/LP germline events from ClinVar to O/LO somatic data in the COSMIC database absent in cBioPortal (resulting in a comparable number of total events, n = 57,147). We found 18 TSGs replicated these findings and 3 additional TSGs with significant differences only when comparing the COSMIC dataset to ClinVar (Figure S5, Table S7).

To identify the molecular consequence categories driving significance, we calculated adjusted standardized Pearson’s residuals (ASPR). Frameshift, stop-gain, splice acceptor, splice donor, missense, inframe insertion, and large indels had significant ASPRs (≥3) in at least 1 TSG and were plotted in a heatmap (as shown in the Figure 4). Overall, splice sites, inframe insertions and large indels contributed little to significant findings across the 18 TSGs with replicated findings. We found only three TSGs (*DICER1*, *TP53*, and *SMAD4)* with more somatic missense events than expected, but nine TSGs, including some of the genes with largest datasets, e.g., *APC*, and *RB1*, had more germline missense and/or somatic stop-gains than expected. Frameshifts were enriched in a similar number of genes in both the somatic and germline datasets. Thus, despite similar loss of function consequences of stop-gain and frameshift variants, germline and somatic distributions of these variant types exhibit substantial differences.

**Figure 4:**
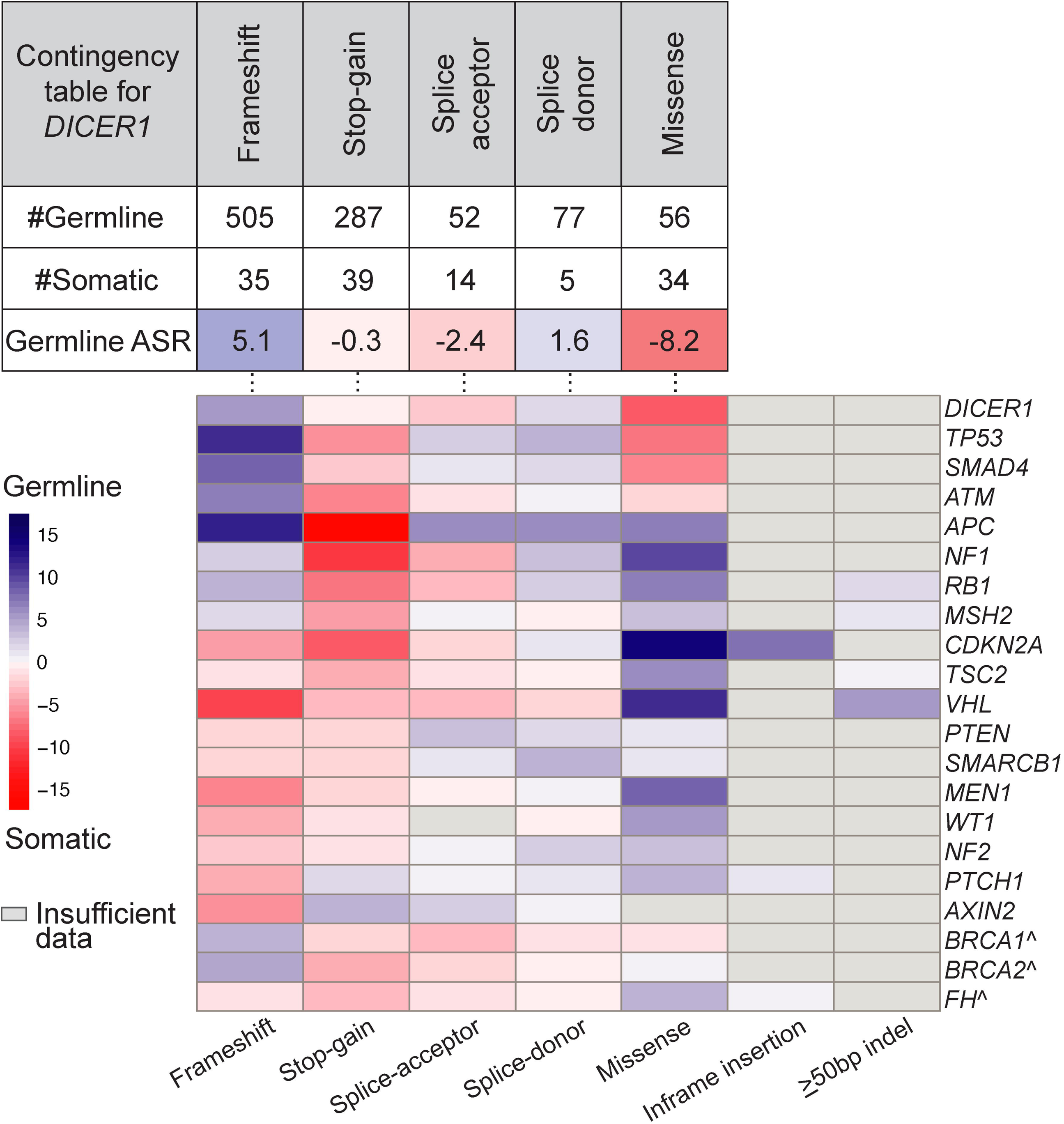
TSGs with significantly different distribution patterns of variant molecular consequence for total P/LP germline and O/LO somatic events. At the top of the figure is an example of the *DICER1* contingency table, with observed germline and somatic event counts (chi-squared test q<0.05) along with calculated adjusted standardized Pearson’s residuals with respect to germline data (Germline ASR) (above), followed by a heatmap of Germline ASRs for 21 TSGs with significant findings (below). ^ indicate 3 TSGs with significant findings that did not meet the significance threshold when comparing non-overlapping somatic data from COSMIC but displayed the same pattern. P/LP, pathogenic/likely pathogenic; O/LO, oncogenic/likely oncogenic.

### Tissue-specific mutational mechanisms contribute to the distinct somatic variant profile

The differences between frameshift and stop-gain distributions might be a result of distinct underlying mutational mechanisms shaping the respective mutational profiles. O/LO somatic events occur in tumors from a wide variety of underlying tissues. Tissue types have different susceptibilities to somatic mutagenesis based on age, cell replication rate, DNA repair efficiency, and exposure to external mutagens^43–46^. We characterized the effect of tumor tissue type in driving the distinct somatic mutational consequence profiles. O/LO somatic data from unique samples in cBioPortal and COSMIC (Methods) were combined to increase sample size and existing tissue type ontologies were used to harmonize the datasets. We identified that the largest proportion of O/LO somatic events (n=68,547) derived from bowel cancers (n=17,135), followed by lung (n=8,225), stomach (n=6,990), and other cancer types (Figure S6A, per gene distribution in Figure S6B). We focused the analysis on germline and somatic frameshift and stop-gain events and found that 19 of the 36 TSGs with sufficient data had significant tissue differences (Bonferroni adjusted p < 0.05) (Figure 5A, Table S8). These included 16 TSGs already identified when analyzing the combined somatic dataset (figure 4) and *STK11*, *CDKN1B*, and *CDH1* which were not captured previously.

**Figure 5:**
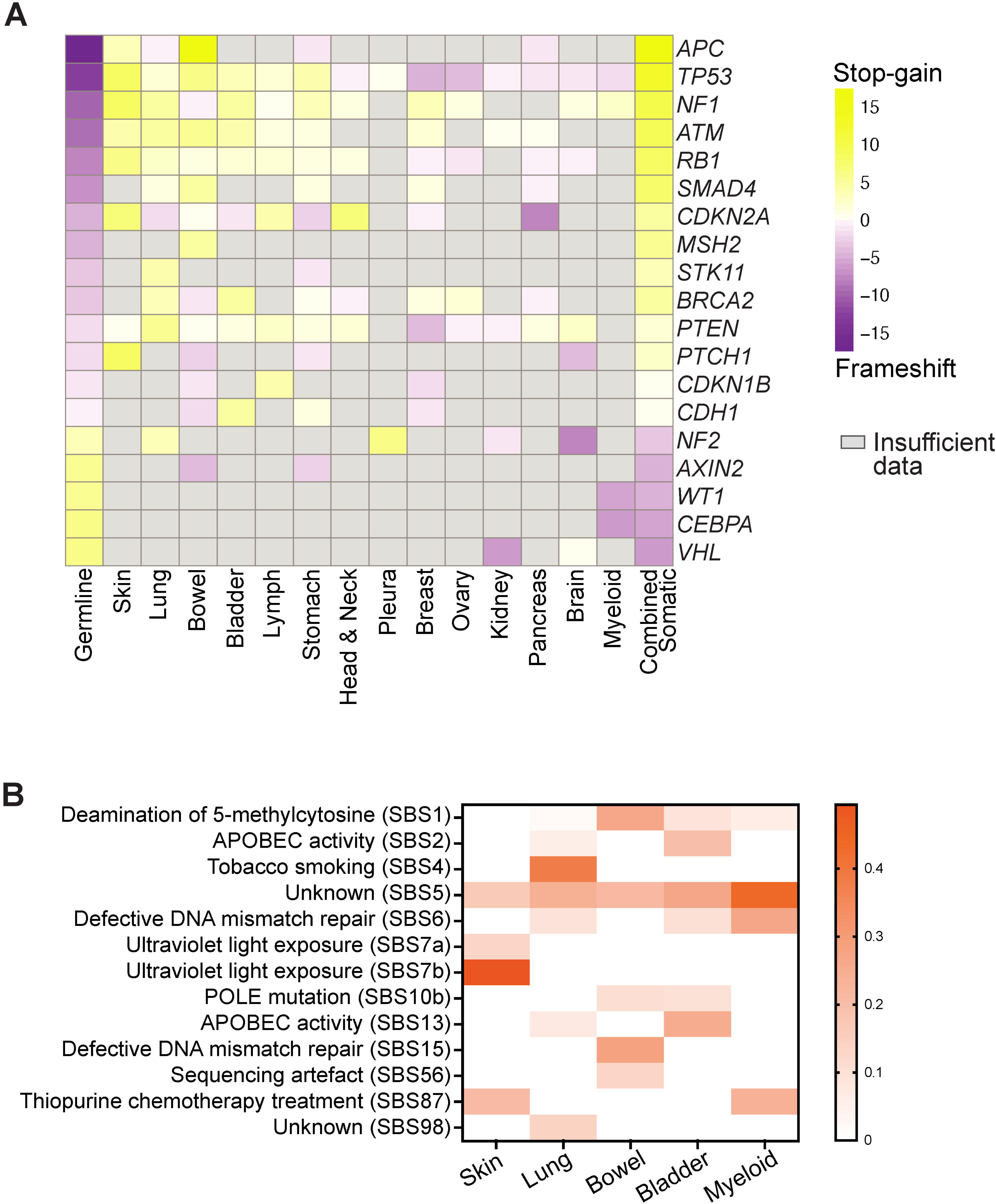
Tissue-specific susceptibilities to somatic mutagenesis contribute to the distinct distributions of O/LO somatic frameshift and stop-gain events. (A) Heatmap of adjusted standardized Pearson’s residuals with respect to stop-gain events (stop-gain ASR) calculated for the distribution of frameshift and stop-gain events across germline and somatic parent tissue types for 19 TSGs with significant findings. Stop-gain ASR for germline (left), each tumor parent tissue type in decreasing order of number of significant TSGs with excess stop-gain ASR and stop-gain ASR for somatic data combined across tissues compared to germline (right). (B) Known COSMIC mutation signatures indicating tissue-specific environmental factors contributing to the cumulative single base substitution (SBS) profile for somatic skin, lung, bowel, bladder, and myeloid tumors. O/LO, oncogenic/likely oncogenic.

To identify the tissue types driving significance here, we again calculated and plotted the ASPR for each significant TSG (Figure 5A). Skin as the tumor tissue type had the greatest number of TSGs (7 of 19) with an enrichment of somatic stop-gains (ASPR≥3), followed by lung (6 of 19 TSGs), bowel (5 of 19 TSGs), and bladder (5 of 19 TSGs). Tumor development in these tissues has been extensively reported to demonstrate exposure-related base substitution mutagenesis mechanisms that could explain the excess of somatic stop-gains compared to indel frameshifts (see for example^45,47^).

To assess exposure mechanisms, we performed a modified mutational signature analysis using COSMIC’s SigProfiler Assignment tool. We identified the SBS mutational signatures specifically contributing to the cumulative SBS mutational profile across all tumor samples in a given tissue type for all 40 TSGs (as opposed to the more typical analysis of all somatic mutations in one tumor sample) (see Methods). Indeed, we found expected exposure mechanisms such as ultraviolet light exposure signatures SBS7a and 7b contributing only to skin tumors (>50% signature contribution to cumulative skin mutational profile), tobacco smoking signature SBS4 seen only in lung tumors (38% contribution), and defective DNA mismatch repair SBS15 along with deamination of 5-methylcytosine SBS1 prominent in bowel tumors (28.1% and 26.4% contribution, respectively) (Figure 5B, Table S9). In addition, we see APOBEC activity signatures SBS2 and SBS13 in the context of bladder tumors (44.9% total contribution). These mutational signatures are tissue specific and not observed in tissue types, e.g. myeloid malignancies less likely to be influenced by exogenous factors. Exogenous mutagenesis mechanisms are a distinguishing characteristic driving enrichment of somatic stop-gain events in tumor types exposed to environmental or chemical mutagens compared to germline variation.

Of the 19 TSGs with significant findings, 5 TSGs had excess somatic frameshifts, including *WT1* and *CEBPA* in myeloid tumors. P/LP germline variants in *WT1* are associated with an increased risk of Wilms tumor, a rare type of kidney cancer, which constituted only 7 of 125 (5.6%, Figure S6B) *WT1*-mutated tumors in our somatic dataset. Somatic data for *WT1* largely comprised the more common myeloid malignancies (77 of 125 tumors, 61.6%) (Figure S6B). Thus, different tumor types associated with germline variants from that observed in somatic data for the same gene is another underlying mechanism driving distinct somatic variant profiles for some TSGs such as *WT1*.

### Germline and somatic events preferentially cluster in 39 TSGs

We compared the distribution by location of P/LP germline and O/LO somatic total events by a moving difference in fraction of data along TSG cDNA (Methods, Figure 6A). For this, we performed parallel analyses using cBioPortal and COSMIC data independently. Regions enriched equally in both germline and somatic data cancel each other out such as the cancer hotspot at codon 233 (arginine) in *PTEN*, and only those regions with preferential enrichment clustering (>±2.5 SD) in one or the other dataset are captured (Figure 6B). We identified a total of 103 clusters of varying widths across the 40 TSGs. Apart from *CDKN1B*, 39 TSGs had at least 1 cluster and up to a maximum of 9 (in *NF1*) reflecting potential differences in analysis power with large datasets. Counts and cluster positions varied across the TSGs (see select TSGs in Figure 6D, all genes depicted in Figure S7). We found over 3 times more somatic clusters than germline (Figure 6C), ranging in size from 50 bp (cluster [1] in *MLH1*, *CYLD* and cluster [2] in *ATM*) to 643 bp (cluster [3] in *APC*) (Table S10). Somatic clusters also make up 75% of the clusters identified with replicate analysis comparing germline data to COSMIC somatic data (Figure S8, Table S11).

**Figure 6:**
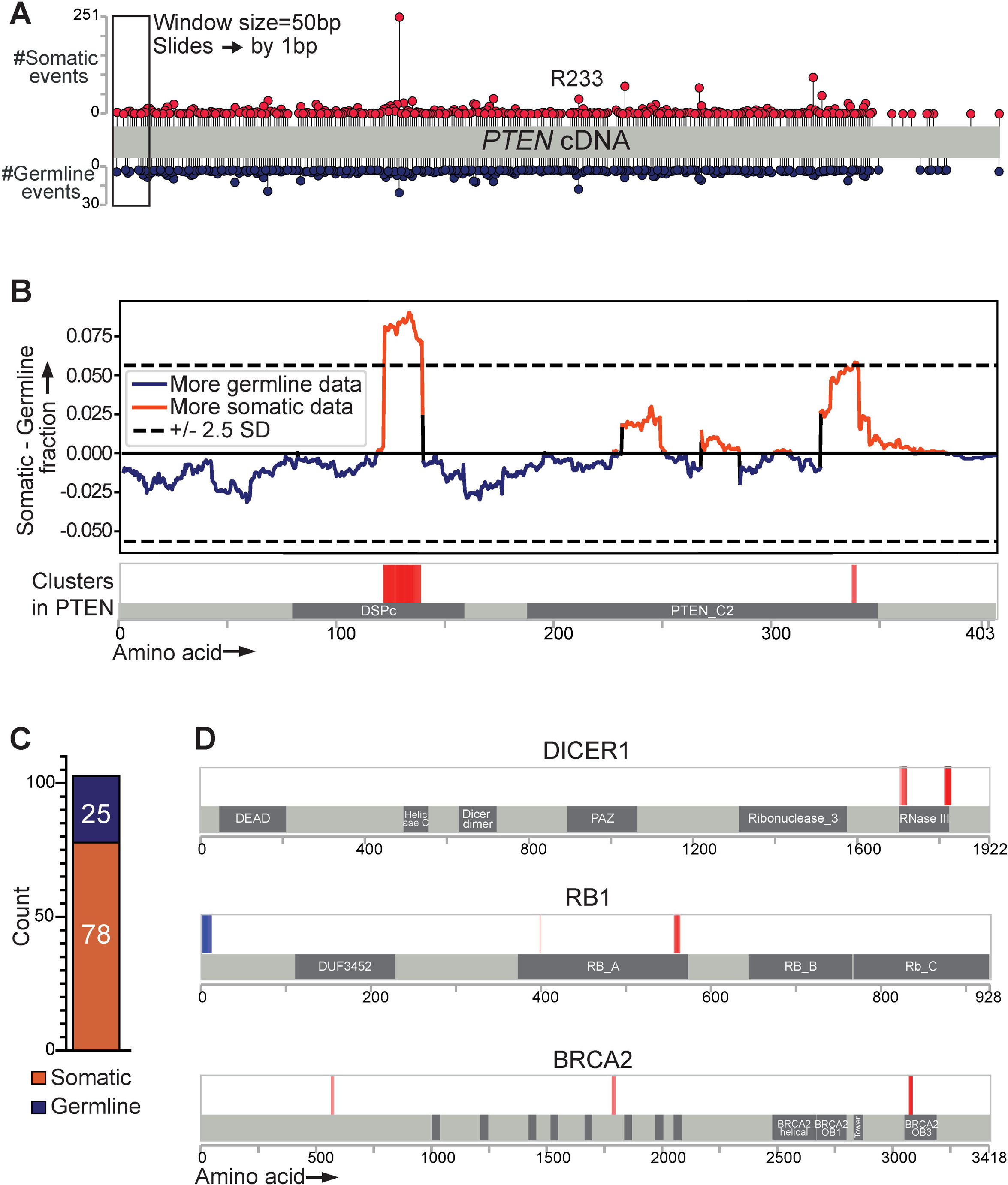
Preferential clustering of P/LP germline and O/LO somatic events across the TSG cDNA. (A) Lollipop plot of total O/LO somatic events (above) and total P/LP germline events (below) along the coding sequence of PTEN. (B) Plot of the moving difference in fraction of O/LO somatic and P/LP germline events across the cDNA capturing preferentially clustered regions ±2.5 SD from 0. Shown below is a one-dimensional heatmap denoting preferentially clustered regions related to amino acid sequence. Red=somatic cluster, blue=germline cluster. No preferential clustering is detected at the known hotspot locus, R233 (https://www.cancerhotspots.org/#/home) shown in (A), demonstrating how the moving difference in fraction algorithm captures regions enriched in only germline or somatic data, but not both. (C) Total count of germline and somatic preferential clusters. (D) One-dimensional heatmaps for *DICER1, RB1,* and *BRCA2*, illustrating variable cluster counts, loci, and window lengths with predominance of somatic clusters versus germline clusters. See Figure S7 for the remaining TSGs. P/LP, pathogenic/likely pathogenic; O/LO, oncogenic/likely oncogenic.

### Multiple mutational mechanisms drive preferential clustering

We explored the molecular consequence categories found in clusters, evaluating category proportions (stop gain, frameshift, etc.) in each of the 103 clusters (germline or somatic using the cBioPortal data) (Figure 7A). Somatic clusters were found to contain single predominant molecular consequences defined as >80% of the events within an individual cluster more often than germline (Somatic=33/78 vs germline=2/25), (Figure 7B). Somatic stop-gains are predominant in only 2 clusters. Somatic missense events were predominant in 8 clusters, with 7 containing known missense cancer hotspot variants in *DICER1*, *SMAD4*, *TP53*, *SMARCA4*, and *NF1* all recorded in cancerhotspots.org (Table 2). *DICER1*, *SMAD4*, and *TP53* correlate with the gene-wide result of having significantly more somatic missense events compared to germline (Figure 4). Of note, there are many more hotspots recorded in cancerhotspots.org, including missense and stop-gain changes in the TSGs under study. However, our cluster analysis focuses on differences between somatic and germline distributions. This demonstrates that hotspots are rarely exclusive to somatic data and can be valuable for germline variant classification as per ClinGen’s recommendations^7^. In contrast, frameshift events are the molecular consequence in 23 of 33 (70%) somatic clusters with a predominant variant type. Of these, 20 clusters identified in 16 TSGs contain recurring single bp deletion or duplication frameshift variants in homopolymer regions (Table 3). The recurring events are less likely to represent sequencing artefacts since they were identified in only 20 of the 87 homopolymer regions found in these 16 TSGs.

**Figure 7:**
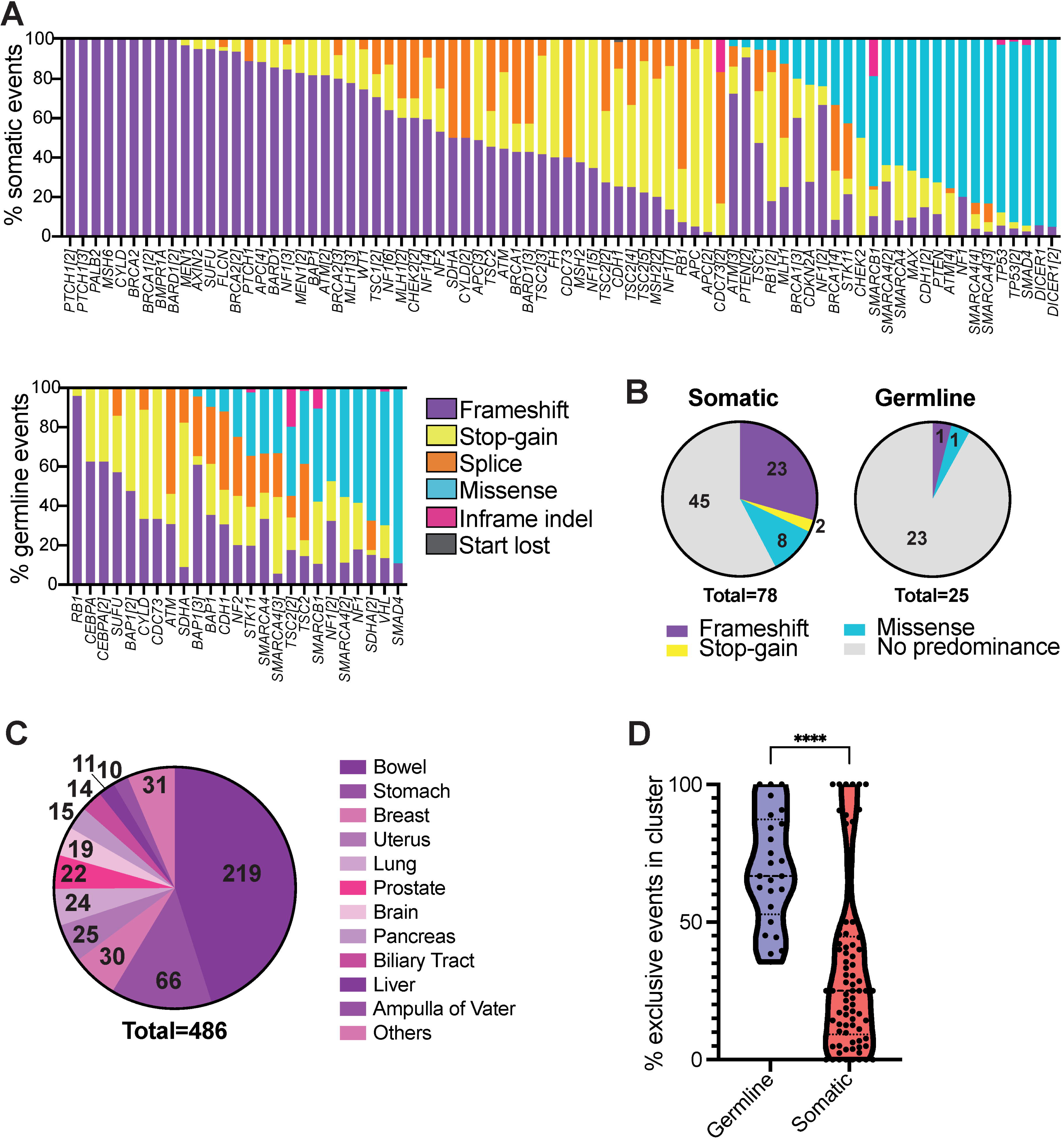
Types of P/LP germline and O/LO somatic variants driving preferential clustering. (A) Proportions of molecular consequence categories represented in 78 somatic clusters (above) and 25 germline clusters (below). (B) Count of somatic and germline clusters grouped by predominant category, defined as >80% events in the cluster. More somatic clusters are composed of a predominant category, mainly frameshift events. (C) Parent tissue types in which recurring O/LO somatic frameshift clusters in homopolymer runs were detected. Bowel and stomach are the top tissue types represented and are known to be associated with a microsatellite unstable mutational mechanism. (D) Distribution of percent exclusive variants in 25 germline clusters compared to 78 somatic clusters. Data are represented in a violin plot depicting the distribution of individual points. Mann Whitney test, p<0.0001. P/LP, pathogenic/likely pathogenic; O/LO, oncogenic/likely oncogenic.

**Table 2.**
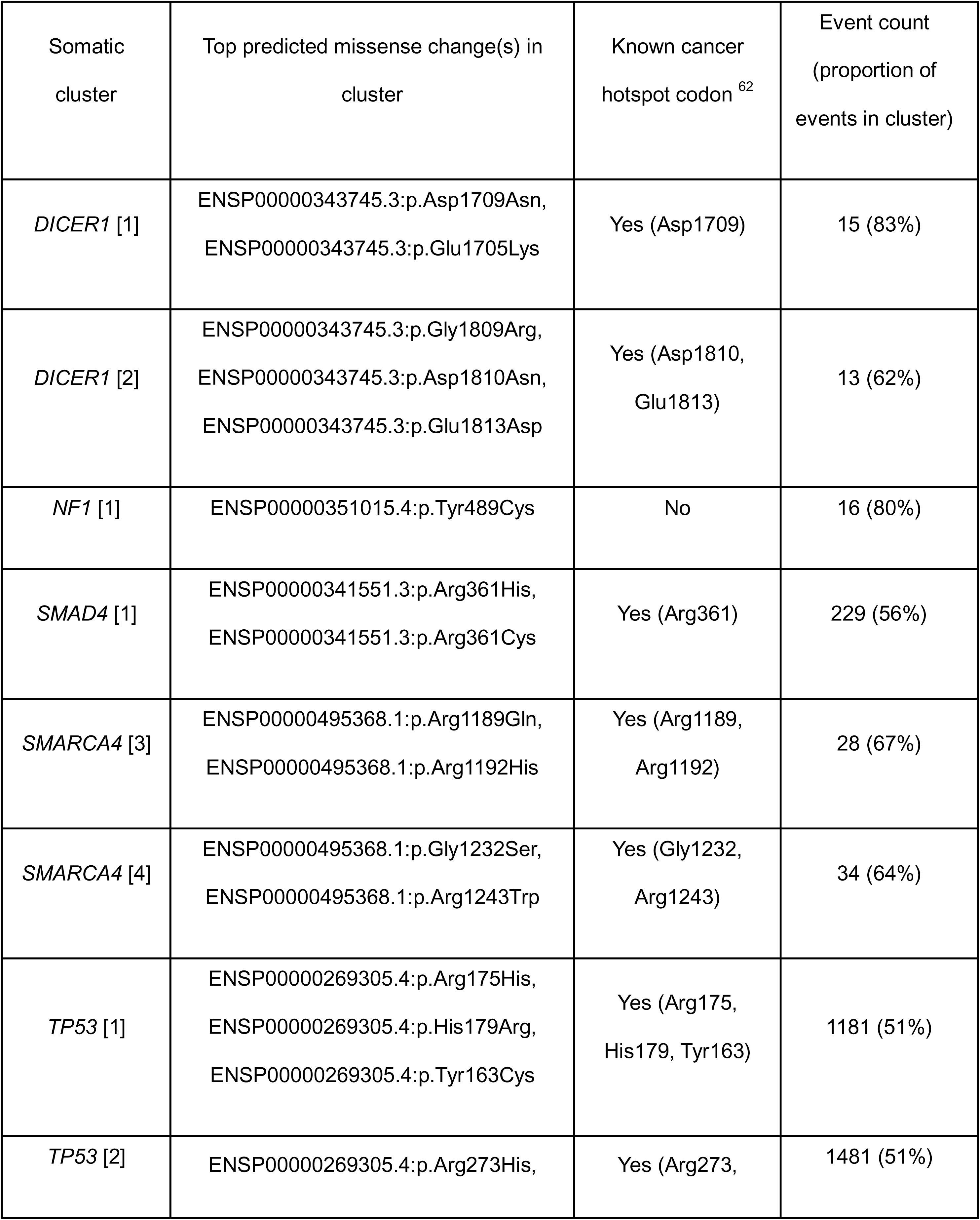
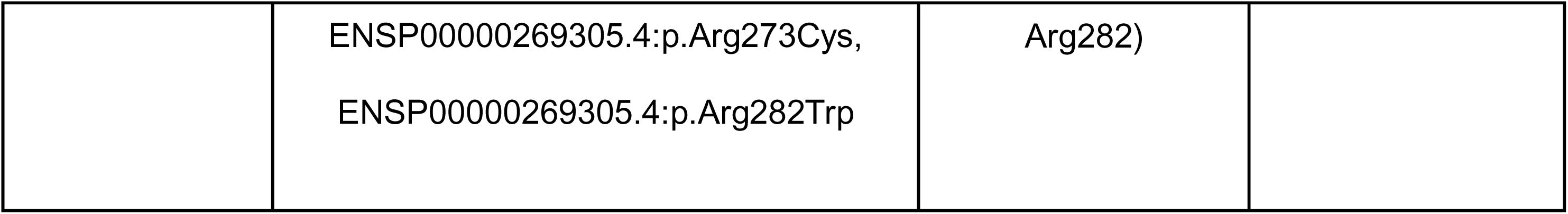
Top oncogenic/likely oncogenic missense variants in predominantly missense somatic clusters.

| Somatic cluster | Top predicted missense change(s) in cluster | Known cancer hotspot codon <sup>62</sup> | Event count (proportion of events in cluster) |
| --- | --- | --- | --- |
| *DICER1* [1] | ENSP00000343745.3:p.Asp1709Asn,<br>ENSP00000343745.3:p.Glu1705Lys | Yes (Asp1709) | 15 (83%) |
| *DICER1* [2] | ENSP00000343745.3:p.Gly1809Arg,<br>ENSP00000343745.3:p.Asp1810Asn,<br>ENSP00000343745.3:p.Glu1813Asp | Yes (Asp1810,<br>Glu1813) | 13 (62%) |
| *NF1* [1] | ENSP00000351015.4:p.Tyr489Cys | No | 16 (80%) |
| *SMAD4* [1] | ENSP00000341551.3:p.Arg361His,<br>ENSP00000341551.3:p.Arg361Cys | Yes (Arg361) | 229 (56%) |
| *SMARCA4* [3] | ENSP00000495368.1:p.Arg1189Gln,<br>ENSP00000495368.1:p.Arg1192His | Yes (Arg1189,<br>Arg1192) | 28 (67%) |
| *SMARCA4* [4] | ENSP00000495368.1:p.Gly1232Ser,<br>ENSP00000495368.1:p.Arg1243Trp | Yes (Gly1232,<br>Arg1243) | 34 (64%) |
| *TP53* [1] | ENSP00000269305.4:p.Arg175His,<br>ENSP00000269305.4:p.His179Arg,<br>ENSP00000269305.4:p.Tyr163Cys | Yes (Arg175,<br>His179, Tyr163) | 1181 (51%) |
| *TP53* [2] | ENSP00000269305.4:p.Arg273His, | Yes (Arg273, | 1481 (51%) |
|  | ENSP00000269305.4:p.Arg273Cys,<br>ENSP00000269305.4:p.Arg282Trp | Arg282) |  |

**Table 3.**
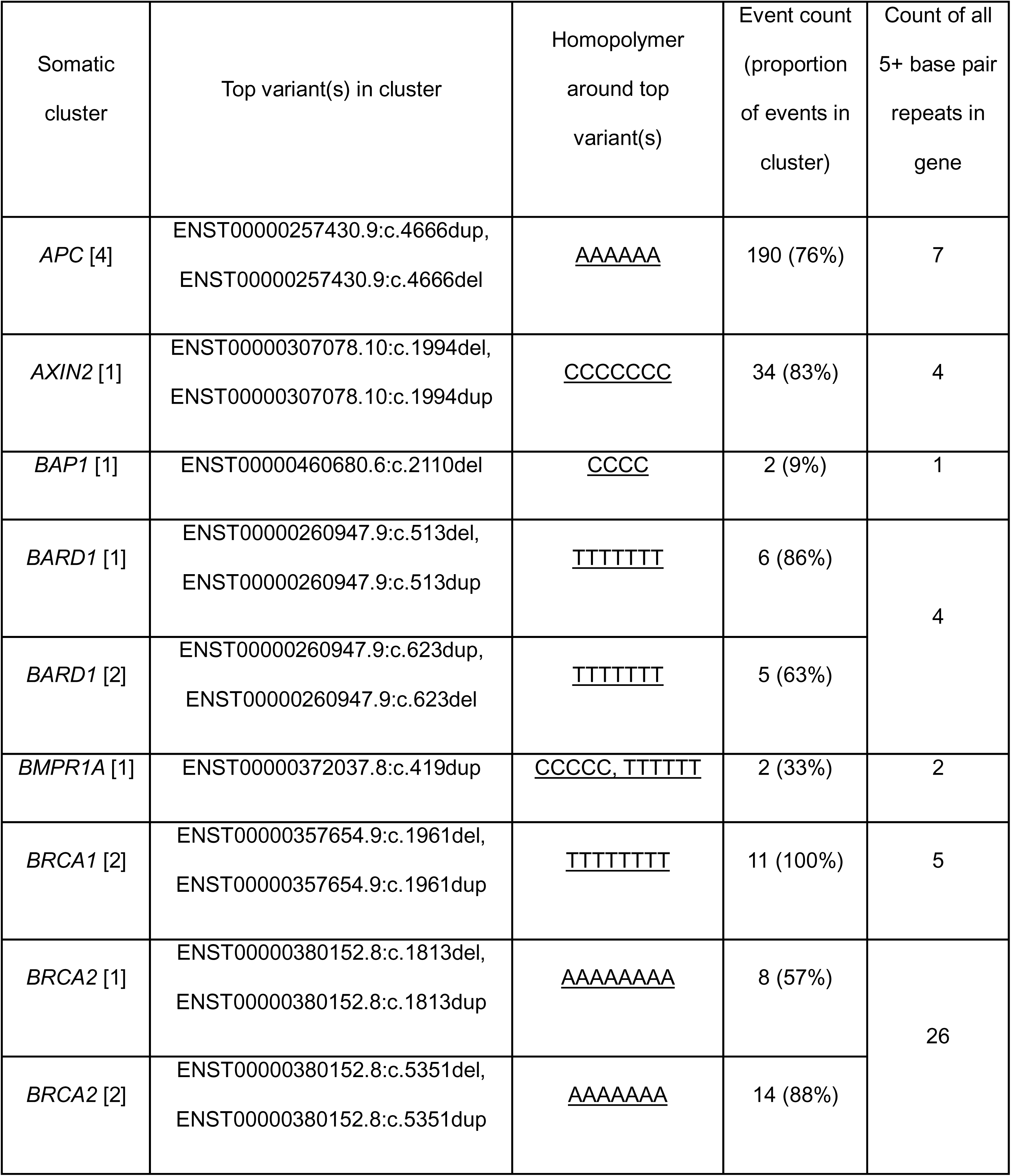
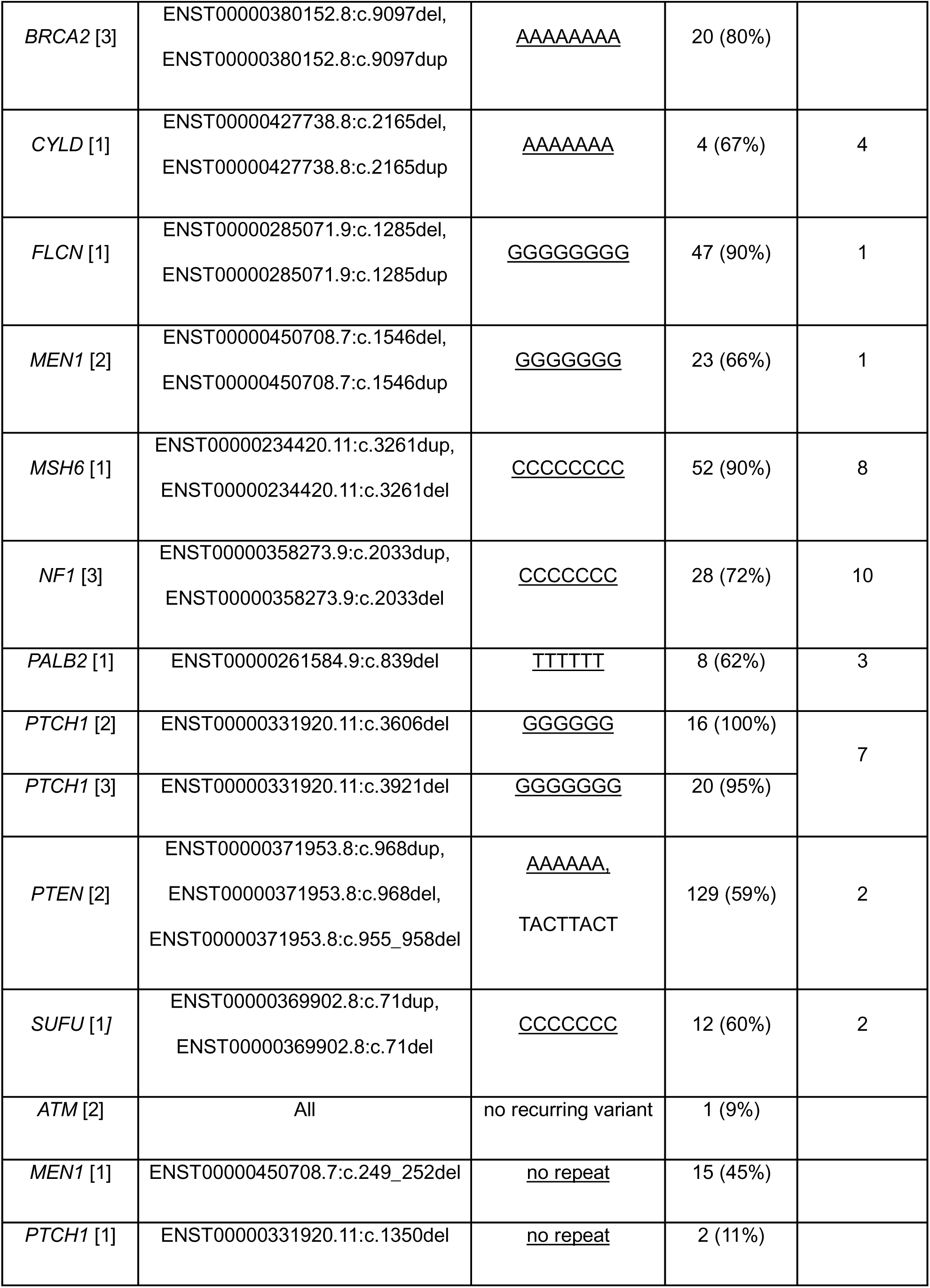
Top recurring variants in somatic predominantly frameshift clusters with the homopolymer they lie in and total count of 5+ base pair homopolymer repeats in the gene.

To assess whether recurrent frameshift variants in homopolymers arise in select tumor tissue types we categorized the 486 available tissue type annotations described earlier and found a majority (70%) of annotations in bowel, stomach, breast, and uterine cancers (Figure 7C), which are tissues having known microsatellite instability (MSI) mutagenesis mechanisms^48^. Eighty-five of 104 tumor samples (81.7%) with these variants and having MSI status information provided in cBioPortal were MSI high (Figure S9), despite eliminating tumors with high TMB (>10) from this analysis (Methods). This reinforces tissue-type specific mutagenesis mechanisms underly distinct somatic mutational profiles.

The majority (66%) of all clusters comprise combinations of different molecular consequence categories without a predominant category. We evaluated if there was a more wide-spread selection bias across all clusters of regions resistant to nonsense mediated decay (NMD) but found only 11 of 78 somatic clusters contain NMD escaping variants (Figure S10). Similarly, only 8 of 23 germline clusters had NMD escaping variants (including two in the single-exon gene *CEBPA*). Overall, P/LP variants or O/LO variants that are predicted to escape NMD do not make major contributions to clusters.

We additionally assessed whether clustering is driven by variants that are exclusively in germline or somatic datasets (non-overlapping by identity) given the initial finding that only 9.2% of variants are shared between the two datasets. Germline clusters are comprised of significantly larger proportions of non-overlapping or exclusive variants (median=66.67% events per cluster, IQR=52.78%-87.3%), while variants in somatic clusters are less likely to be exclusive (median=25% events per cluster, IQR=9.24%-44.7%) (Mann Whitney test, p<0.0001) (Figure 7D). Many exclusive variants in germline clusters are observed in genes such as *SMAD4* and *SMARCA4* which are classical tumor suppressor genes, but specific P/LP germline variants in these genes can also be associated with phenotypes other than cancer predisposition, such as Myre Syndrome (*SMAD4*) or Coffin Siris Syndrome (*SMARCA4*). This highlights a difference in clinical presentations driven by P/LP germline variants in pleiotropic TSGs represented in ClinVar, while somatic data is limited to cancer driving variants in tumors.

## Discussion

We systematically characterized the landscape of 32,941 P/LP germline and 12,907 O/LO somatic variants in tumors in 40 well described TSGs, using variants extracted from ClinVar, cBioPortal, and COSMIC. Given the shared loss of function impact on the gene’s tumor suppressor function, we expected significant overlap in germline and somatic datasets. Despite large-scale clinical germline cancer predisposition testing and the more recent extensive somatic tumor profiling data available, a surprisingly small number (∼9%) of P/LP germline or O/LO somatic variants were shared when compared by genomic change and predicted protein change. The standardized criteria used to classify germline and somatic variants as P/LP and O/LO, respectively, were largely concordant with a majority of shared variants (86%) being classified as P/LP and O/LO. The minimal overlap of germline and somatic cancer variants is similar to a previous report comparing missense variants in cancer predisposition genes, though not limited to TSGs^49^.

The lack of overlap further motivated a detailed assessment of the correspondence of variant distributions. Unlike oncogenes which are frequently activated by somatic missense hotspots in cancers, only three TSGs (*DICER1*, *TP53*, and *SMAD4*) had more somatic missense events than expected, including the previously noted pattern in *DICER1* with exclusively somatic missense hotspots in the RNase IIIB domain^8^. The recurring somatic missense clusters were previously reported as cancer hotspots. In contrast, many of the TSGs with discordant molecular consequence patterns have more somatic stop-gain events than expected. Although frameshift and stop-gain events can lead to a similar outcome of LoF through either NMD or truncation, we found that the differential distribution arose from external mutagens shaping tissue-specific mutational spectra. Somatic mutational profiles were consistent with known COSMIC mutation signatures attributed to tissue-specific mutagens (such as C>T substitutions in skin driven by ultraviolet light exposure^50^, and T>A substitutions in lung driven by cigarette smoking^51^). Excess somatic stop-gain events might also reflect negative selection of frameshift LoF variants by immune system clearance of tumor cells since frameshifted peptides are more immunogenic^52,53^. Integrating these results may improve probabilistic models that predict which variants are likely of germline or somatic origin from tumor-only or liquid biopsy sequencing^54,55^.

Beyond the molecular consequence distributions, we showed that somatic events cluster more often than germline with few clusters driven by NMD escaping variants. Instead, a larger subset of somatic clusters were comprised predominantly of frameshift variants in homopolymer runs suggesting DNA replication slippage^56^. Interestingly, these frameshift events were detected in tumor samples with low tumor mutation burden (ten or less mutations per mega base) but still represent microsatellite-unstable tissue types^48^. Although high TMB was not reached, inefficient repair mechanisms persist, revealing a vulnerability that can likely be exploited by therapies targeting genomic instability^57^. This mechanism appears specific to somatic variation as no germline variant clusters resulted from recurring homopolymer frameshift variants presumably due to high fidelity of DNA replication^58^.

In contrast to the enrichment of stop-gain events in somatic data, we showed that missense events were enriched in germline data for 11 TSGs. Additionally, variants found in germline clusters are more often absent in the existing tumor data. The differences in prevalence of tumor types driven by germline variants compared to somatic data may explain some distinct mutational patterns. For example, patients with von Hippel Lindau disease carrying germline P/LP *VHL* missense variants (Type 1 VHL disease) are less likely to develop renal cell cancer compared to those with frameshift variants^59^. Given that *VHL* somatic variant data in this analysis are predominantly derived from kidney tumors (95% of tumors with tissue annotations) which is associated with Type 2 VHL disease, it may explain the distinct variant patterns observed. Similarly, the *WT1* TSG displays germline tumor predisposition to Wilms Tumor but somatic mutations are frequently observed in the much more common myeloid tumors such as acute myelogenous leukemia compared to kidney tumors such as Wilms tumor, leading to different variant sets in germline and somatic data. Another feature responsible for a small proportion of the unique germline variant landscape is the reporting in ClinVar of germline alleles associated with non-cancer phenotypes in pleiotropic TSGs. For example, a few specific *SMARCA4* variants are associated with the neurodevelopmental disorder Coffin Siris syndrome, apart from other germline LoF variants causing rhabdoid tumor predisposition syndrome 2.

In this study, we leveraged rich resources of germline and somatic variation. ClinVar is a unique resource of germline variation reflecting over 25 years of clinical sequencing from ∼2800 laboratories, while cBioPortal housed over 200 tumor sequencing studies across multiple tumor types at the time of data extraction. Although the All of Us Research Program is another large resource of germline variation per individual in the United States population, the frequency of P/LP cancer predisposing variants is very low (<3%)^60,61^. Hence, due to lack of a second germline dataset of P/LP variants similar in size to ClinVar, we were able to replicate findings using only an alternate somatic dataset from non-overlapping tumor sequencing studies in COSMIC. We were further limited by lack of germline variant frequency in ClinVar, and hence, we estimated this by counting the number of labs that submitted the variant. This likely underestimates the true range of frequencies of P/LP germline variants. Additionally, although germline data contained non-coding variants, we focused our analyses only on coding variants since somatic datasets lacked curations for non-coding variants in our list of TSGs.

Altogether, we demonstrate that the landscape of germline and somatic cancer genetic coding variants in TSGs are fundamentally different in identity, molecular consequence, and locational clustering adding a layer of complexity to the two-hit hypothesis. We found a combination of factors, e.g., environmental mutagens, functional consequences of missense events, and differences in germline and somatic selection pressures are substantially shaping distinct germline cancer predisposition and somatic cancer mutational landscapes summarized in Table 4.

**Table 4.**
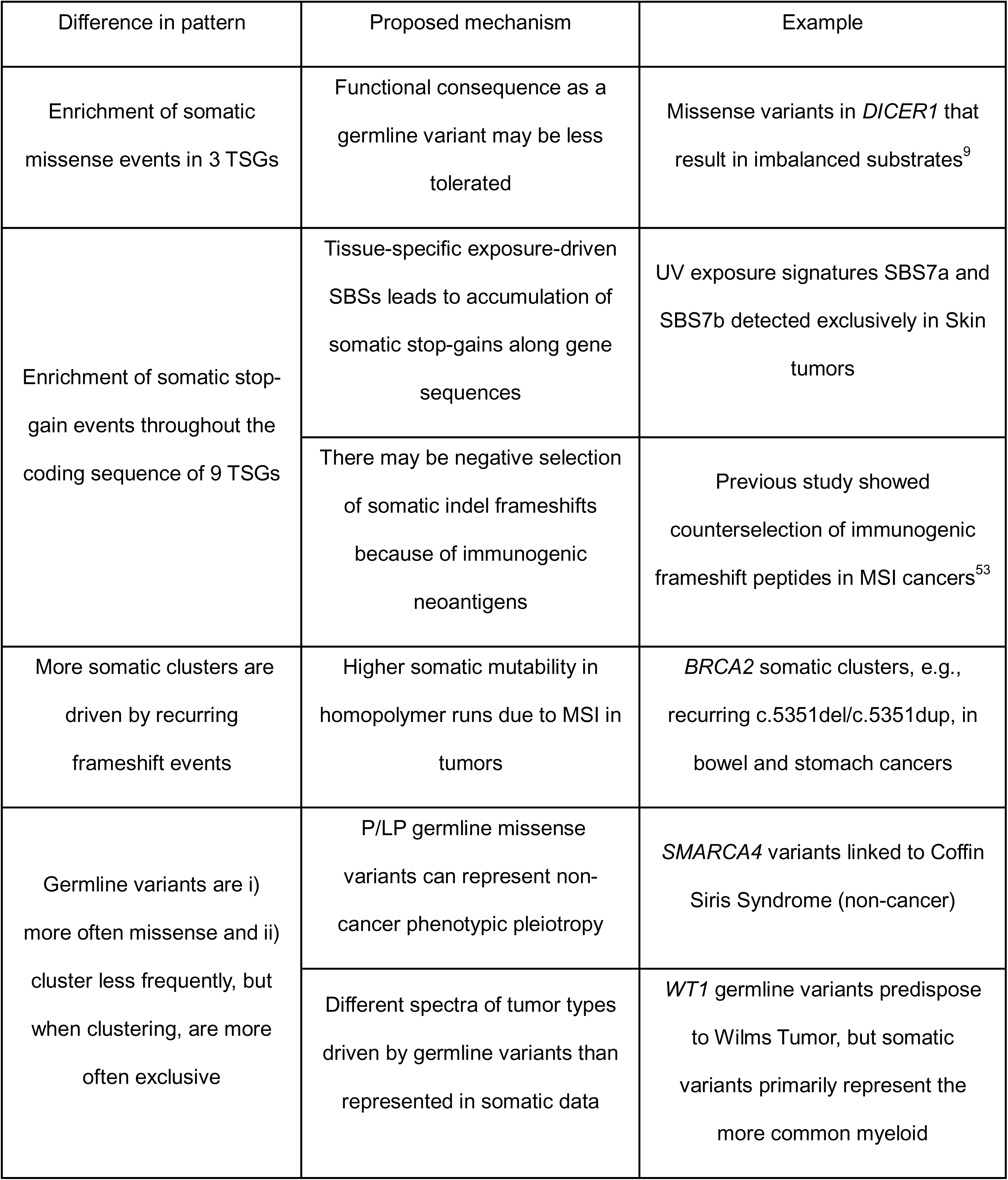
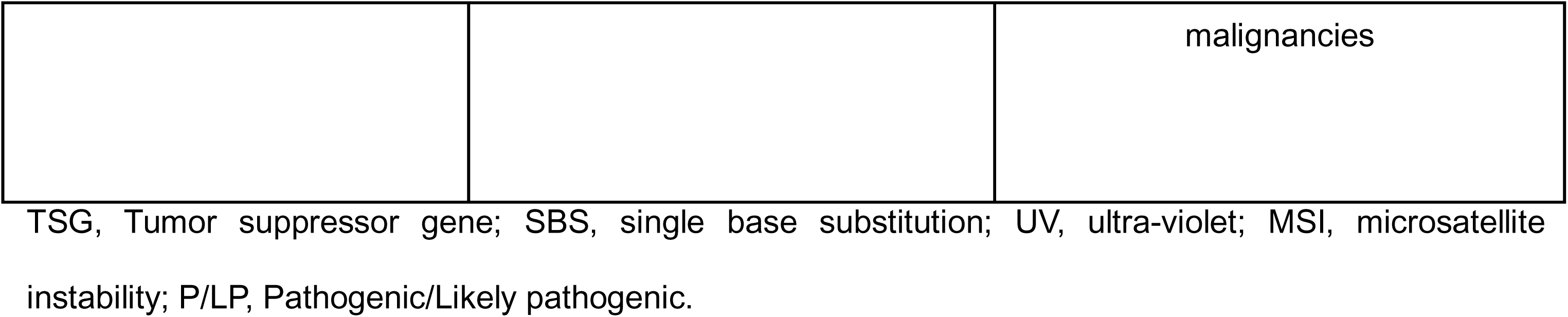
Key differences between germline and somatic variant patterns in tumor suppressor genes and proposed underlying mechanisms.

Although we focused our analyses on well-described TSGs with substantial data, the analyses pipelines that we developed can be applied to other cancer genes as additional data is incorporated from new and expanding cancer genetic studies. From the perspective of clinical diagnostics, the lack of germline and somatic variant overlap supports the need for distinct classification frameworks for cancer associated germline and somatic variants. Insights from tumor-specific selective constraints will be valuable in refining germline variant curation. Further interrogation of the unique distribution patterns described here, on a gene-by-gene basis, will expand understanding of the biology underlying cancer predisposition versus tumorigenesis, aid in cancer variant classification, and support the development of novel cancer therapies.

## Declaration of interests

S.E.P. is a member of the Scientific Advisory Panel of Baylor Genetics.

S.L. completed curricular practical training through the Google Summer of Code 2025 program.

## Acknowledgements

Research reported in this publication was supported by the National Human Genome Research Institute of the National Institutes of Health under Award Number U24HG009649 (to S.E.P.). The content is solely the responsibility of the authors and does not necessarily represent the official views of the National Institutes of Health. Additionally, we would like to thank Susan G. Hilsenbeck for expert feedback on statistical analyses.

## Author contributions

Conceptualization by S.E.P. D.I.R., C.K., and S.L. Data curation, formal analysis, investigation, and writing-original draft by S.L. Resources and funding acquisition by S.E.P. Writing – review and editing by S.E.P., D.I.R., C.K., and S.L.

## Web resources (see Methods section for dates accessed)

ClinGen, https://clinicalgenome.org/

COSMIC Cancer Gene Census, https://cancer.sanger.ac.uk/census

OMIM, https://www.omim.org/

OncoKB, https://www.oncokb.org/

Ensembl, https://useast.ensembl.org/index.html

ClinVar, https://www.ncbi.nlm.nih.gov/clinvar/

NCBI, https://www.ncbi.nlm.nih.gov/

cBioPortal, https://www.cbioportal.org/

COSMIC Cancer Mutation Census, https://cancer.sanger.ac.uk/cosmic/download/cancer-mutation-census

ClinGen Allele Registry, https://reg.clinicalgenome.org/redmine/projects/registry/genboree_registry/landing

OncoTree, https://oncotree.mskcc.org/?version=oncotree_latest_stable&field=NAME

Cancer hotspots, https://www.cancerhotspots.org/

COSMIC SigProfilerAssignment, https://cancer.sanger.ac.uk/signatures/assignment/

CCDS, https://www.ncbi.nlm.nih.gov/projects/CCDS/CcdsBrowse.cgi

NIAID Visual & Medical Arts. (10/7/2024). DNA. NIAID NIH BIOART Source, https://bioart.niaid.nih.gov/bioart/123

NIAID Visual & Medical Arts. (10/7/2024). Generic Cells. NIAID NIH BIOART Source, https://bioart.niaid.nih.gov/bioart/172

Venn Diagram Generator, Bioinformatics and Research Computing, Whitehead Institute, http://barc.wi.mit.edu/tools/venn/

Lucid, https://lucid.co/

GraphPad Prism, https://www.graphpad.com/

## Data and code availability

The datasets/codes generated during this study are available at the Germline-Somatic-AnalysisPaper repository on GitHub (https://github.com/seplon/Germline-Somatic-AnalysisPaper/tree/main).

---

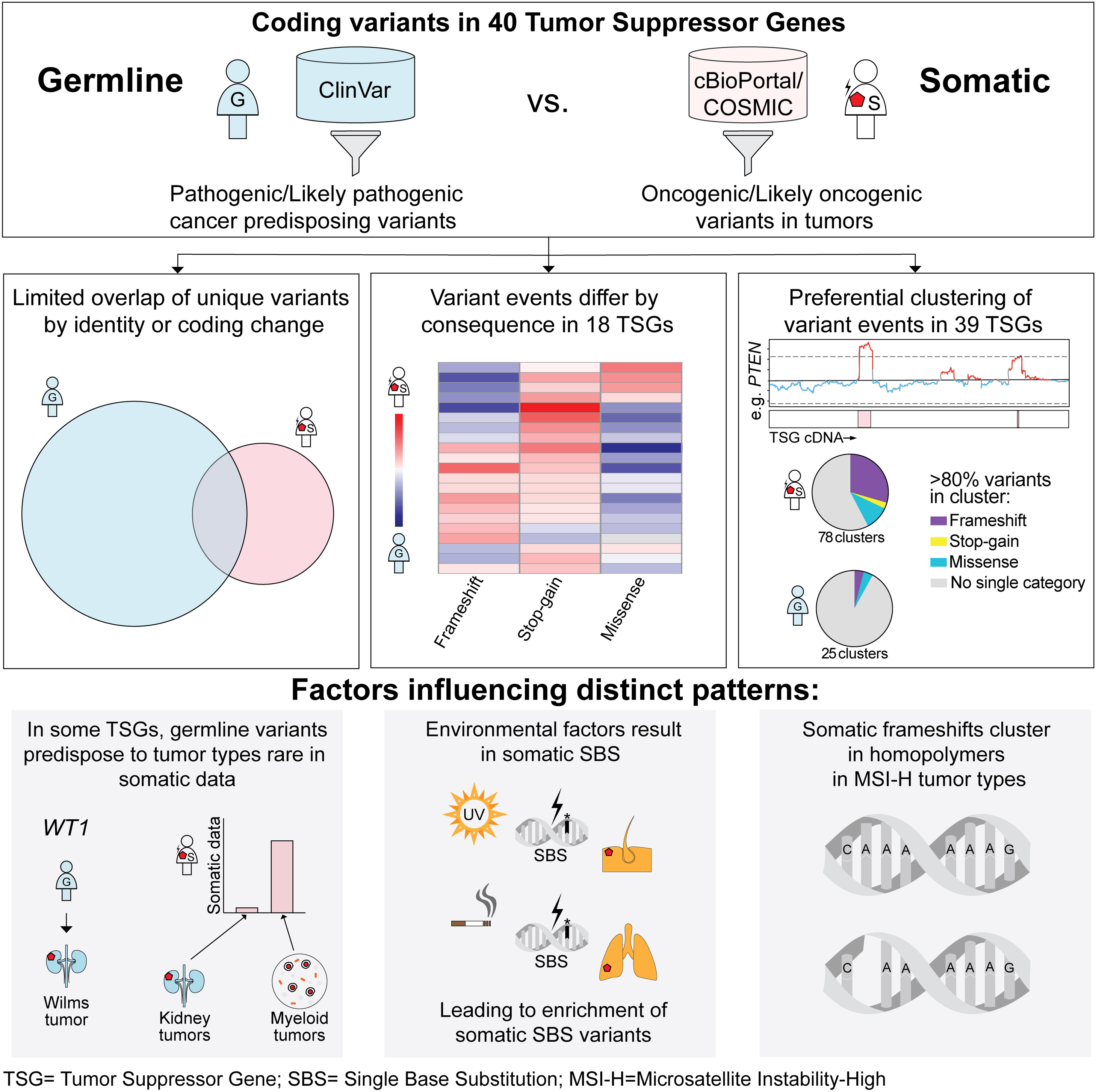

<u>Permissions:</u> None

<u>Supplemental files:</u> Figures (S1-S10) and Tables (S1-S11).

