## Supplemental Figures S1-S10 for "Distinct mutational landscapes when comparing germline and somatic cancer variants in forty tumor suppressor genes"

Supplemental figures and legends:

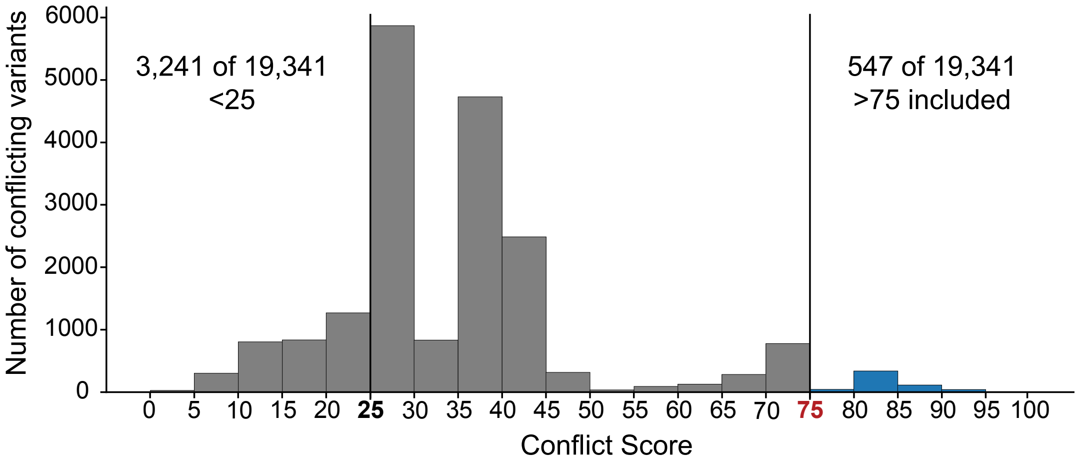


**Figure S1. Distribution of conflict scores for germline conflicting variants in 40 tumor suppressor genes in ClinVar.** Each ACMG/AMP assertion in ClinVar was assigned a value representing the midpoint of the classification's posterior probability e.g. Likely Pathogenic = 0.95 multiplied by 100, (Table S1). The conflict score is the average value across all germline submissions for each variant, calculated for 19,341 variants with conflicting classifications in ClinVar. Variants with a conflict score ≥75 were included in analyses.

ACMG/AMP, American College of Medical Genetics and Genomics and the Association for Molecular Pathology.

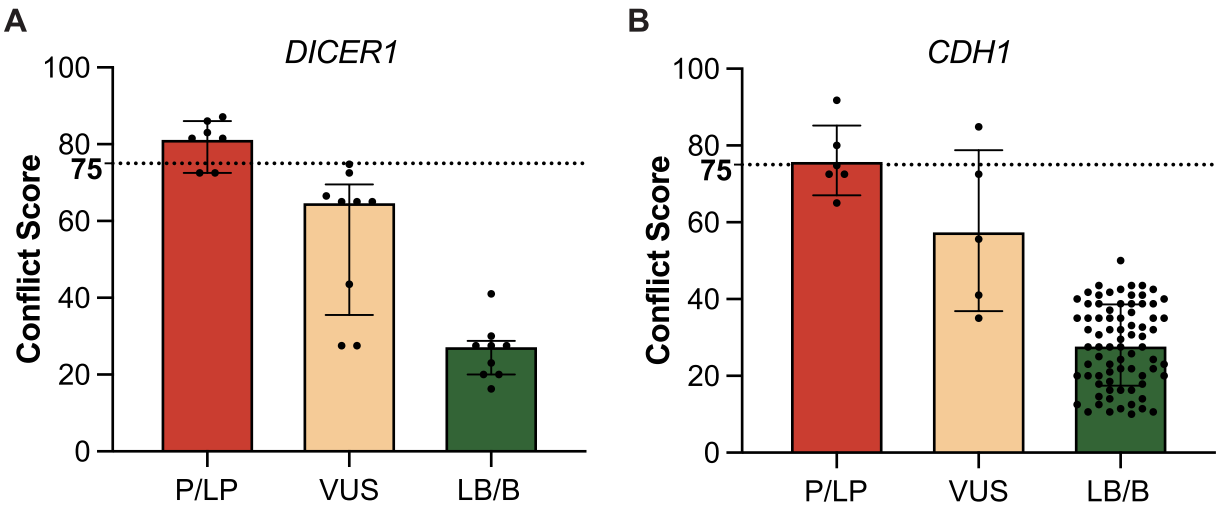


**Figure S2. Distribution of conflict scores for germline variants resolved by ClinGen Variant Curation Expert Panels (VCEPs).** Conflict scores for variants classified as P/LP or VUS or LB/B that previously had conflicting ACMG/AMP classifications across submissions for (A) *DICER1*, and (B) *CDH1*. Data are represented as individual data points within bar plots of mean±SD of conflict scores. All variants classified as LB/B by the ClinGen VCEPs had conflict scores $<75$, while 6 of 7 variants with conflict scores $\geq 75$ were classified as P/LP by the ClinGen VCEPs.

ClinGen, Clinical Genome Resource; P/LP, pathogenic/likely pathogenic; VUS, variants of uncertain significance; LB/B, likely benign/benign; AMP, American College of Medical Genetics and Genomics and the Association for Molecular Pathology; SD, standard deviation.

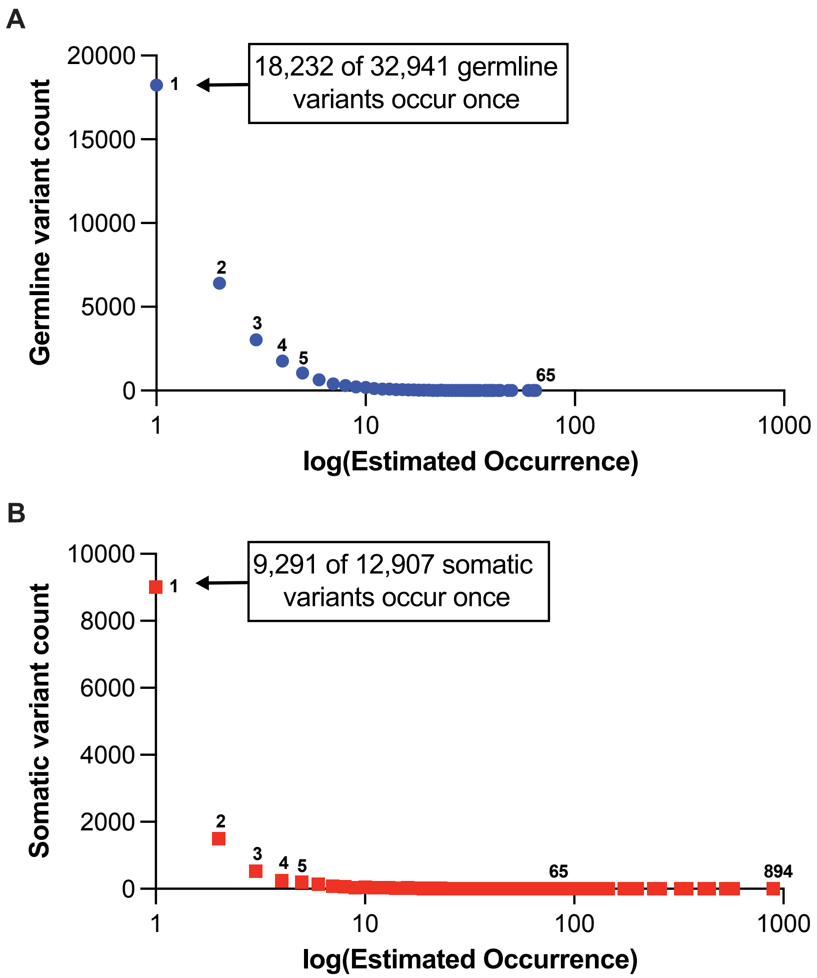


**Figure S3. Distribution of variant frequency by estimated occurrence.**
(A) P/LP germline (ClinVar) variant occurrence for 32,941 unique variants ranges from 1 (18,232 variants occurring once each) to 65 (1 variant occurring 65 times), totaling 77,247 germline events in 40 TSGs.
(B) O/LO somatic (cBioPortal) variant occurrence for 12,907 unique variants ranges from 1 (9,291 variants occurring once) to 894 (1 variant occurring 894 times), totaling 39,781 somatic events in 40 TSGs.
P/LP, pathogenic/likely pathogenic; O/LO, oncogenic/likely oncogenic; TSG, tumor suppressor gene.

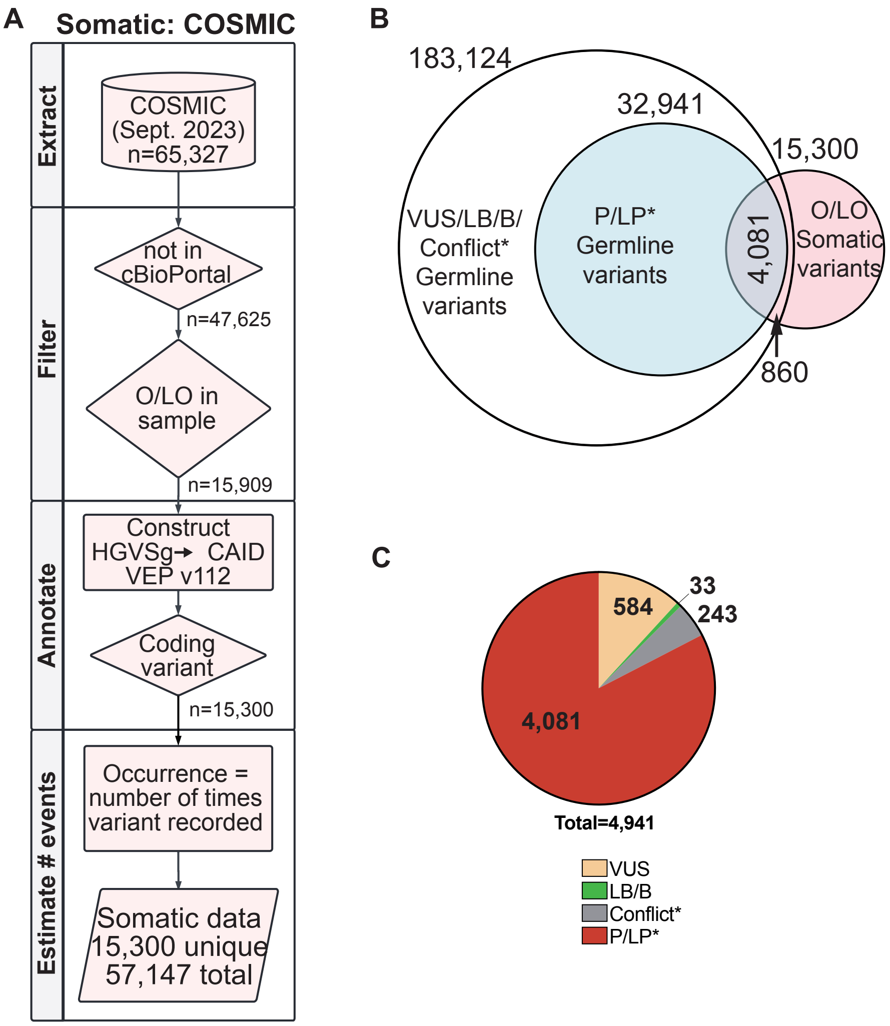


**Figure S4. Extraction and processing of somatic data from COSMIC and overlap with P/LP germline variants from ClinVar.**

(A) Somatic data from COSMIC was filtered to 15,909 unique O/LO variants from samples that were not already included in cBioPortal data. Variants were annotated with their CAID and molecular consequence using VEP (v112) and 15,300 coding variants were retained. Total occurrence per variant was estimated resulting in 57,147 somatic events from COSMIC. Chart created in Lucid (lucid.co).

(B) DNA changes shared between O/LO somatic variants and germline variants. P/LP* in inner blue circle, and germline data unfiltered for pathogenicity in outer circle, containing in addition to 4,081 shared P/LP* variants, only 584 VUS, 33 LB/B, and 243 conflict* variants in ClinVar.

(C) Classification in ClinVar of shared germline and O/LO somatic variants depicted in (B). P/LP*=includes P/LP and conflicting variants with conflict scores ≥75. Conflict*=conflicting variants with conflict scores <75. P/LP, pathogenic/likely pathogenic; O/LO, oncogenic/likely oncogenic; VUS, variants of uncertain significance; LB/B, likely benign/benign; CAID, Clinical Genome Resource Allele Registry ID; VEP, Variant Effect Predictor; COSMIC, Catalogue of Somatic Mutations in Cancer.

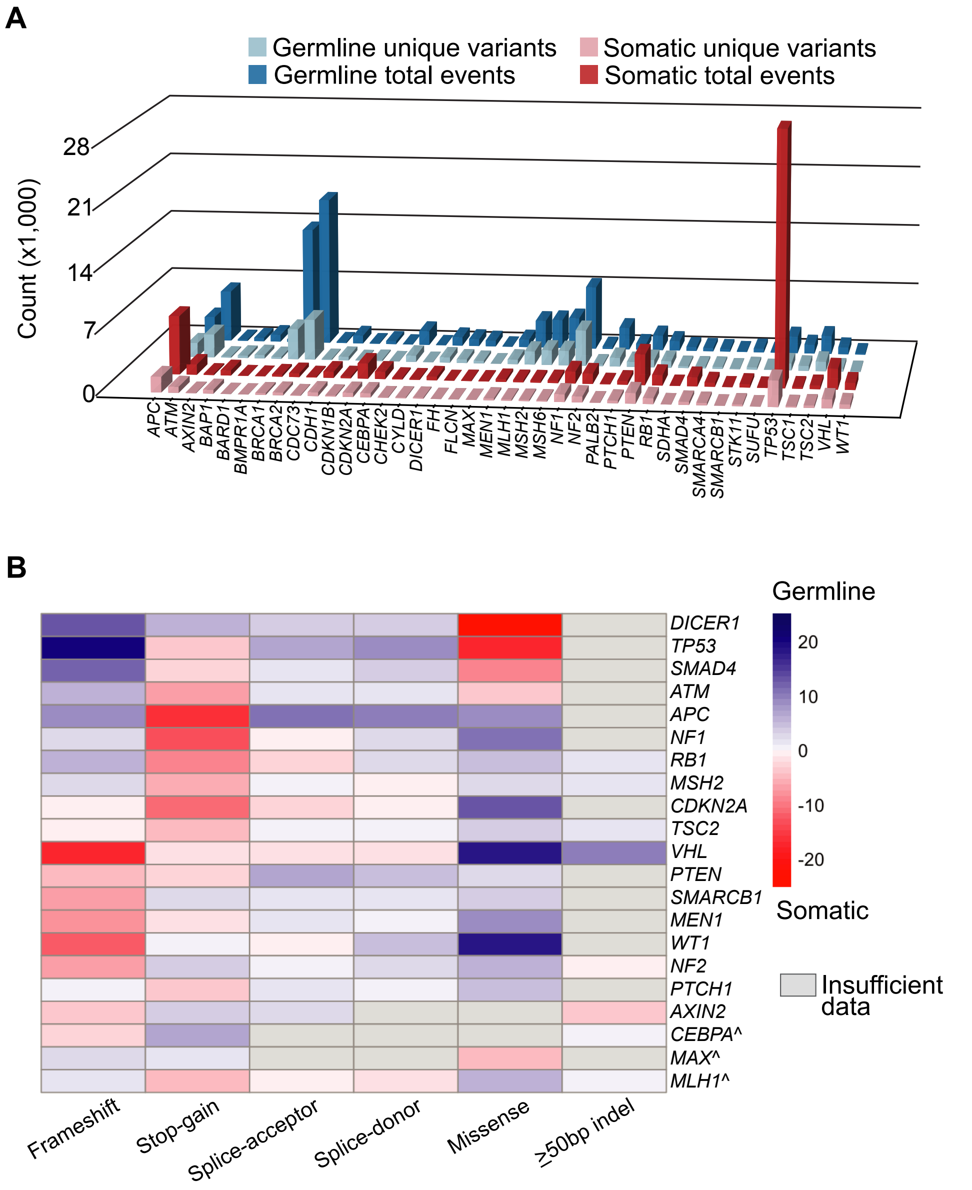


**Figure S5. Total P/LP germline and O/LO COSMIC somatic events have distinct distribution patterns per TSG.**

(A) Unique P/LP variants in ClinVar data and O/LO variants in COSMIC data, and their estimated total events distribute unevenly across the 40 TSGs.

(B) Heatmap of adjusted standardized Pearson's residuals with respect to germline data (Germline ASR) for 21 TSGs with significant findings, 18 of which matched with comparison of datasets from cBioPortal and ClinVar. ^ indicate TSGs that were not significant when comparing non-overlapping somatic data from cBioPortal. P/LP, pathogenic/likely pathogenic; O/LO, oncogenic/likely oncogenic; TSGs, tumor suppressor genes; COSMIC, Catalogue of Somatic Mutations in Cancer.

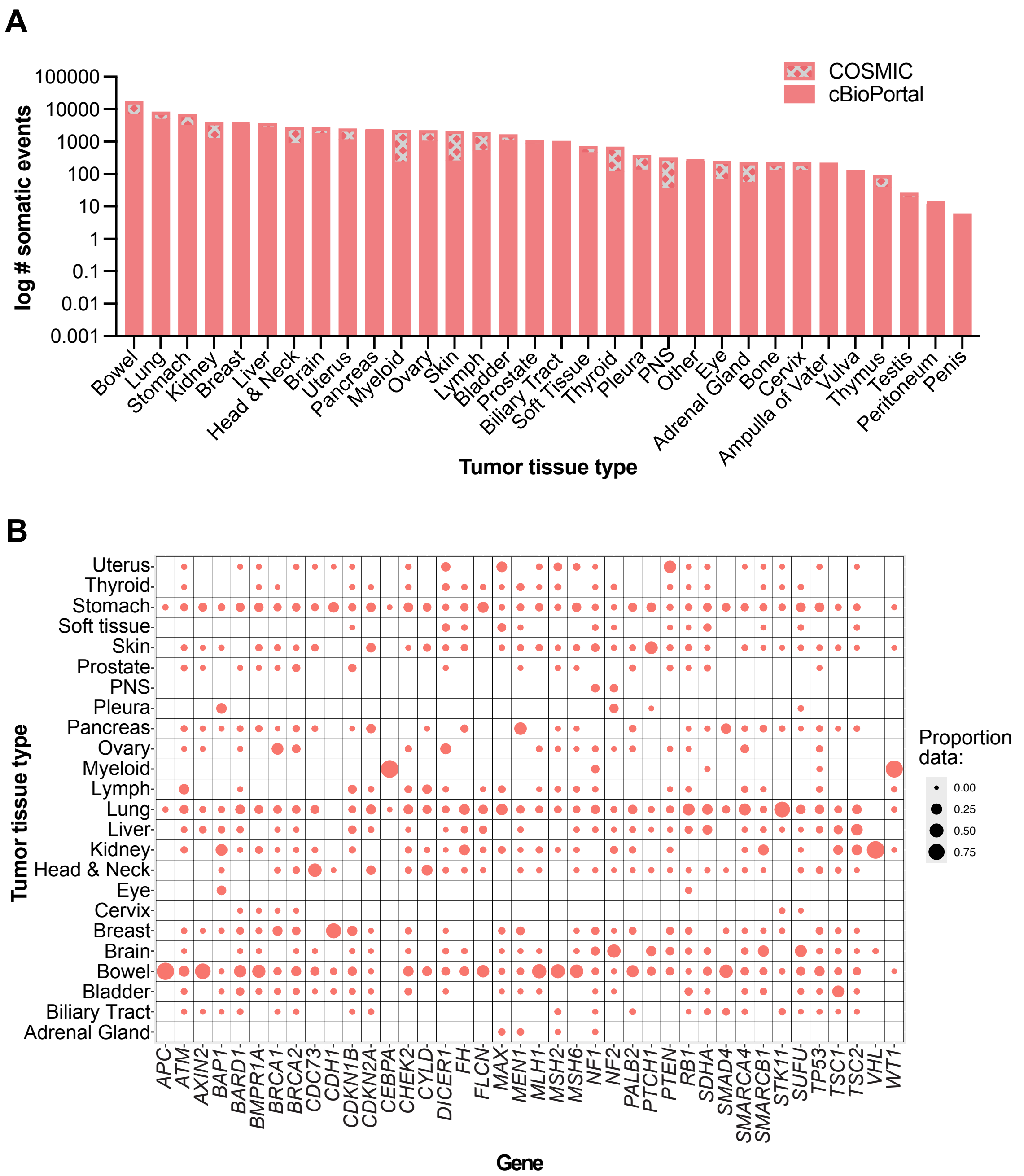


**Figure S6. Distribution of O/LO somatic events by tumor tissue type they were observed in.**

(A) Total O/LO events combined across 40 TSGs and subset by tissue type and the database they were extracted from.

(B) Proportion of O/LO somatic events from cBioPortal and COSMIC combined in each gene by the tumor tissue type they were observed in.

O/LO, oncogenic/likely oncogenic; COSMIC, Catalogue of Somatic Mutations in Cancer; PNS, peripheral nervous system.

cBioPortal

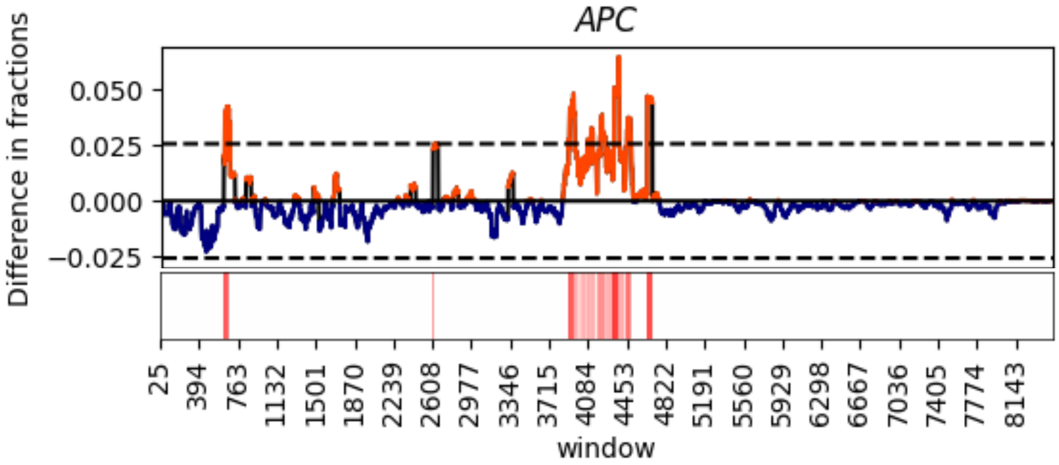


COSMIC

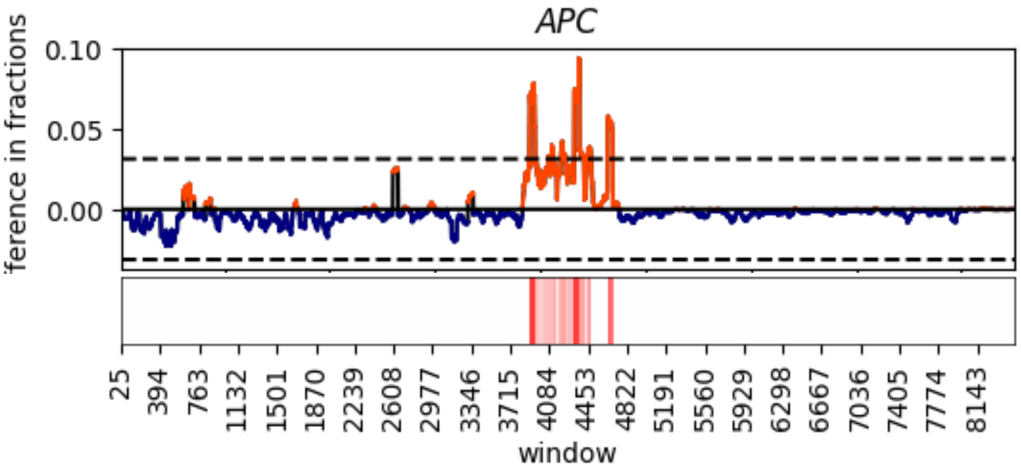

cBioPortal

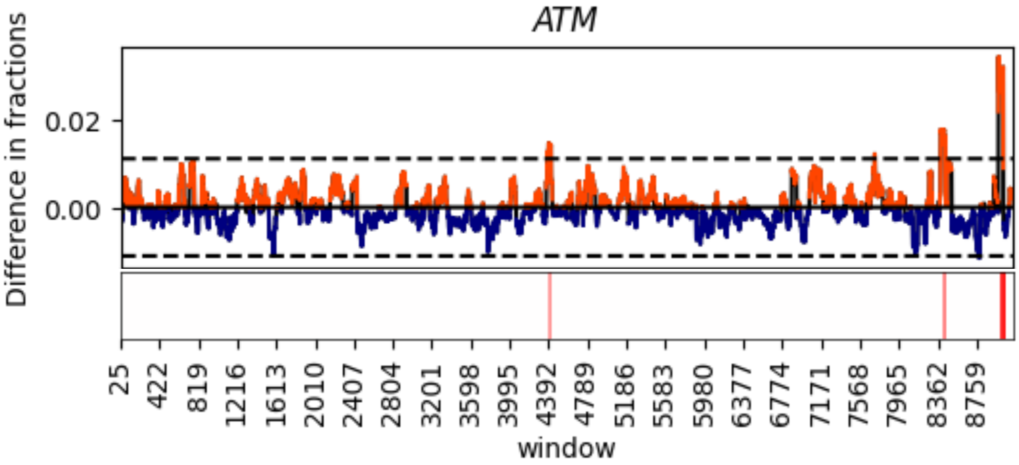


COSMIC

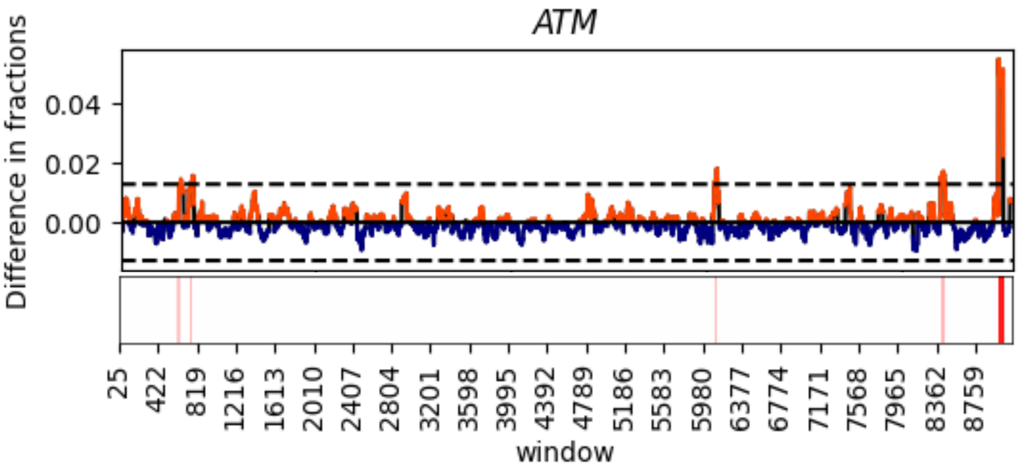

cBioPortal

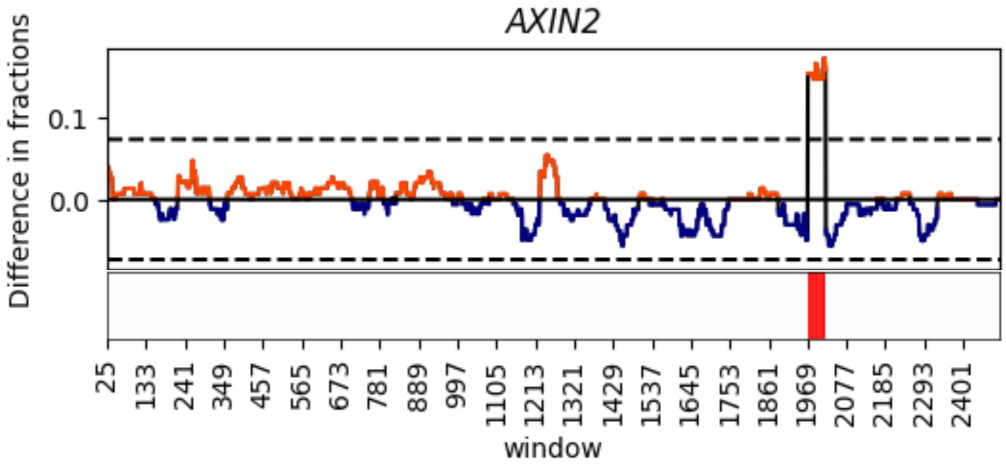


COSMIC

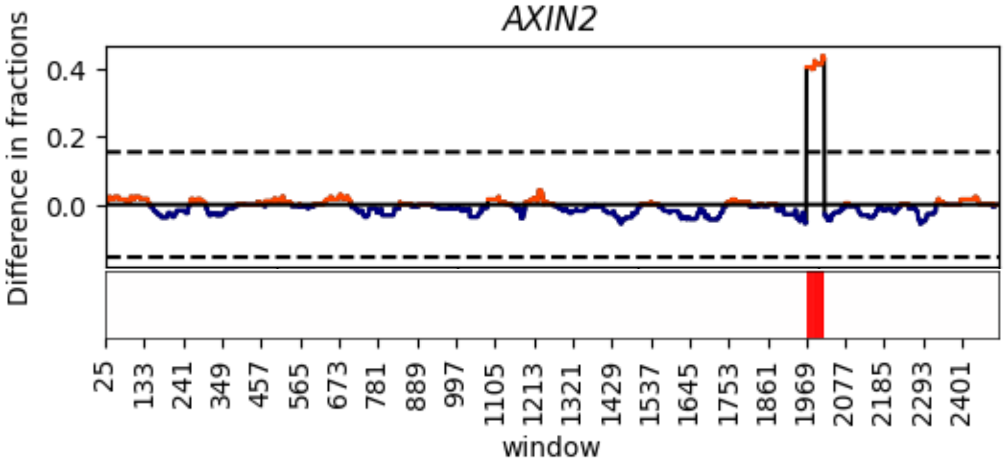

cBioPortal

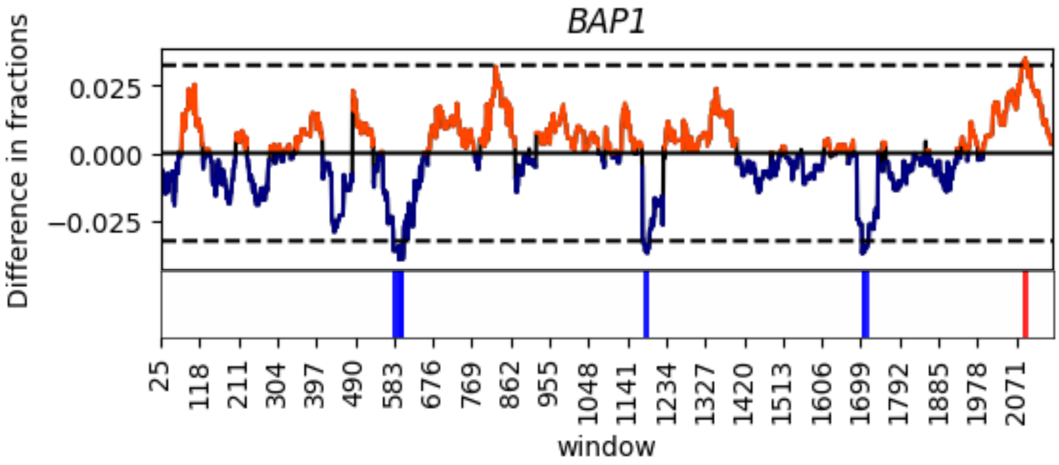


COSMIC

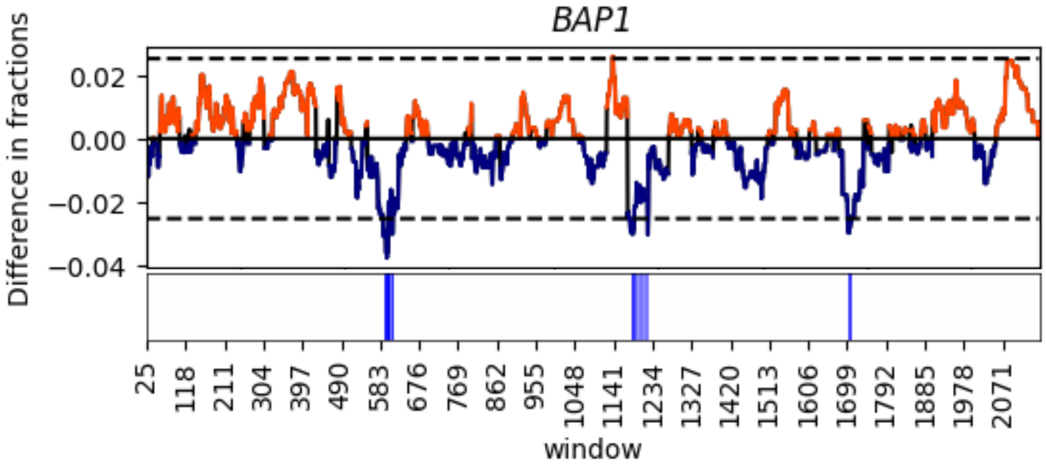

cBioPortal

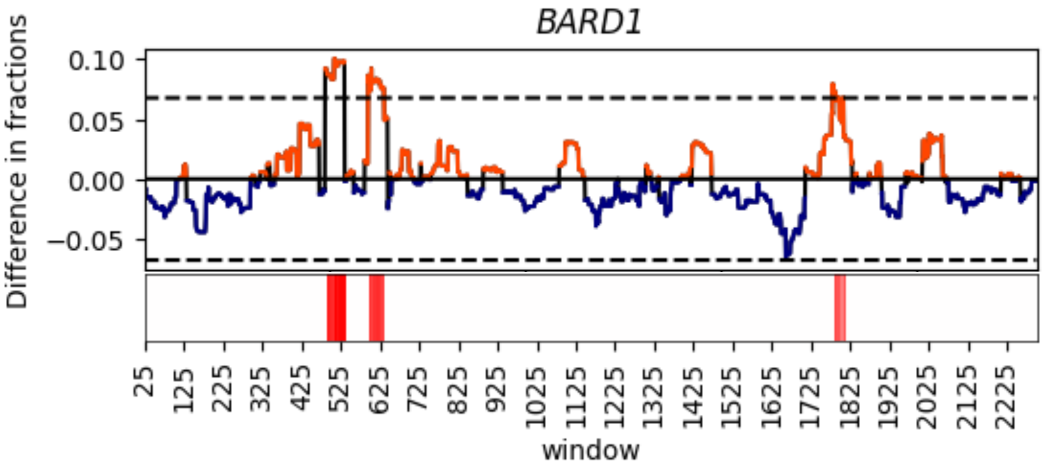


COSMIC

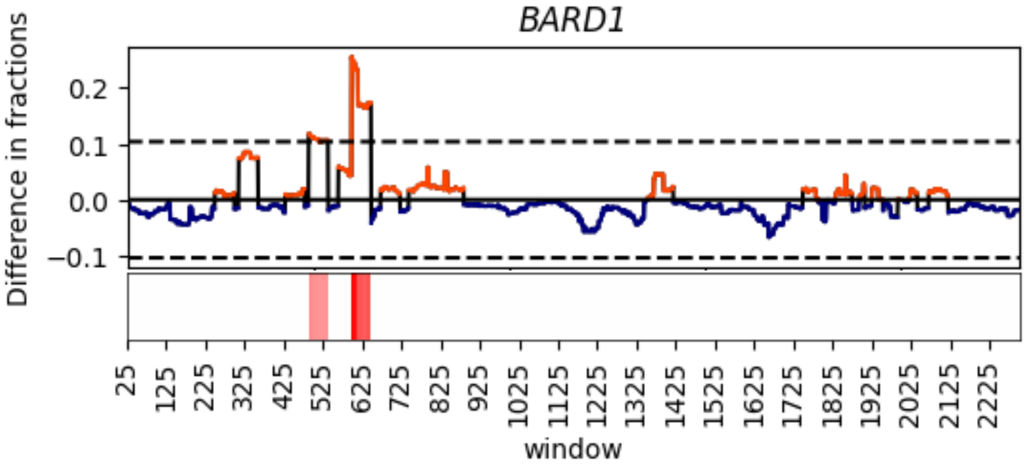

cBioPortal

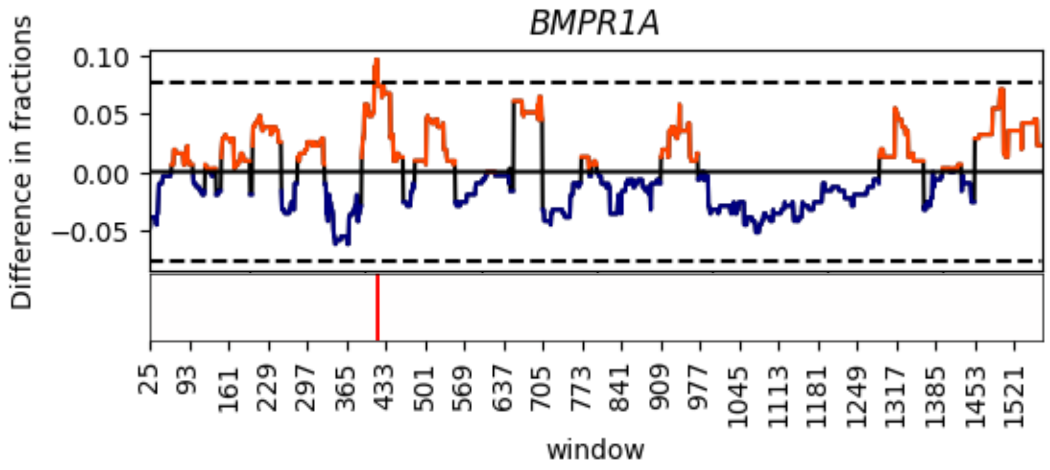


COSMIC

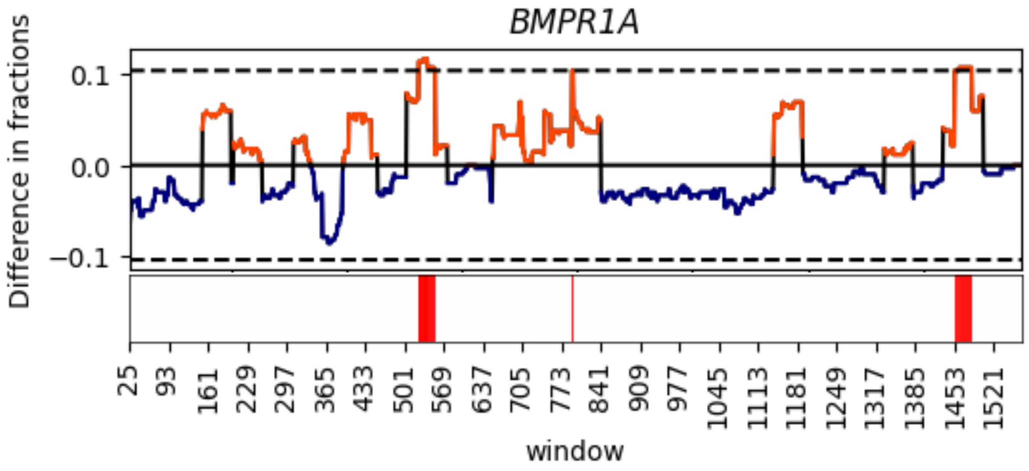

cBioPortal

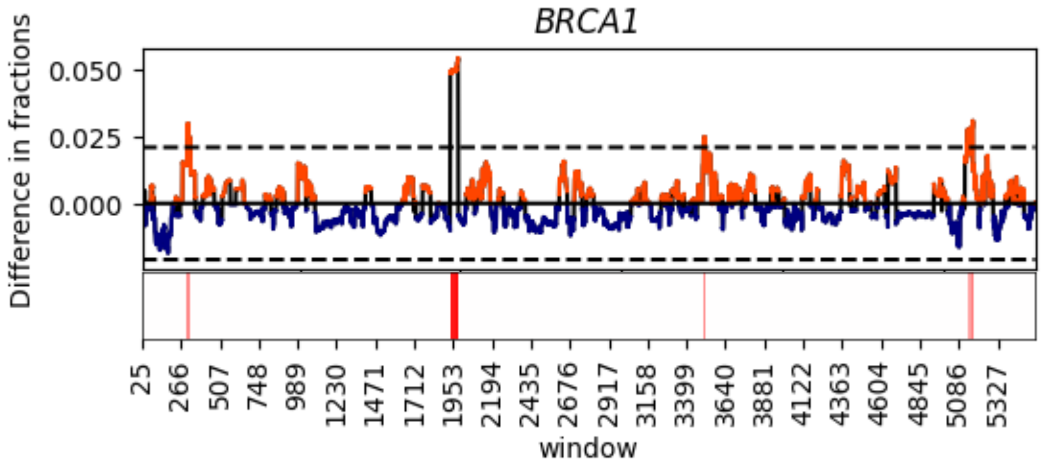


COSMIC

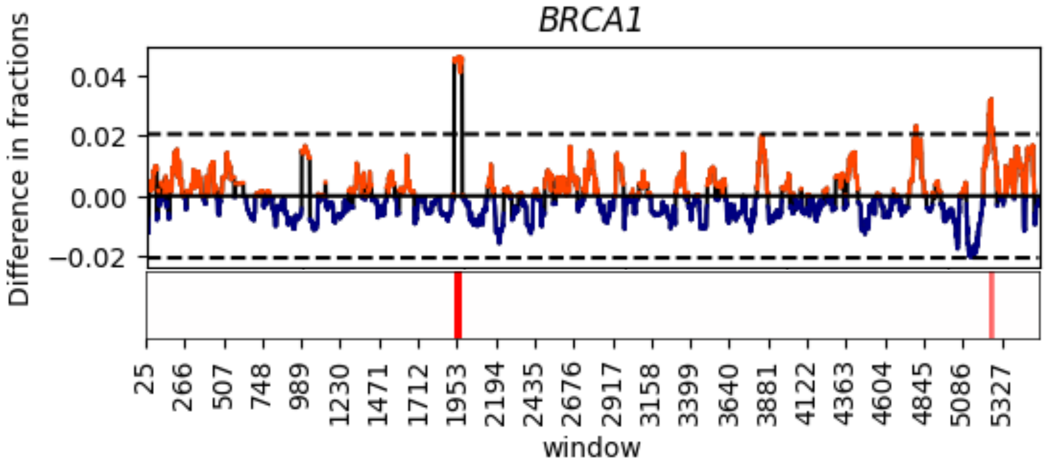

cBioPortal

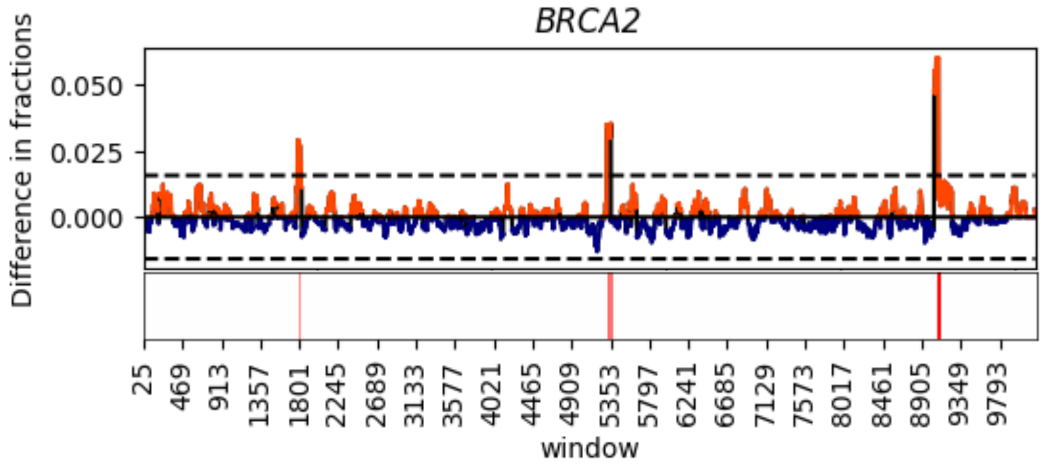


COSMIC

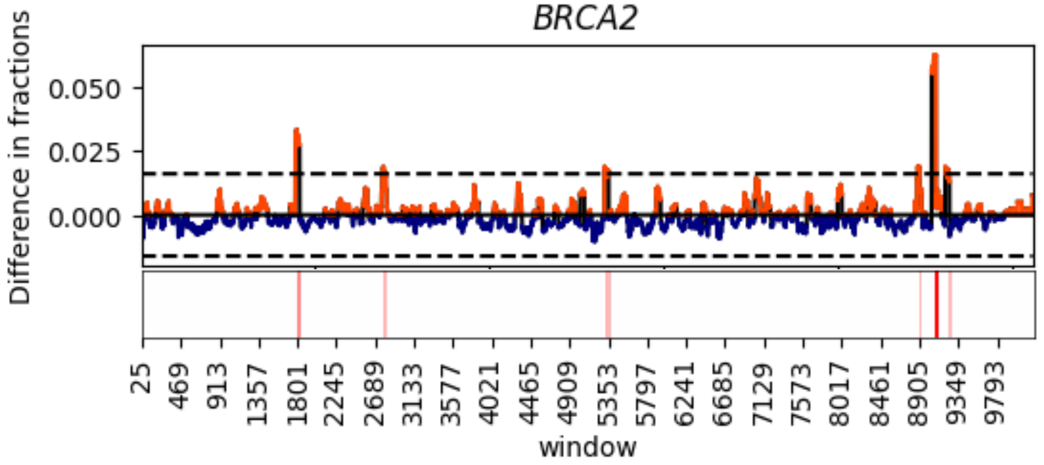

cBioPortal

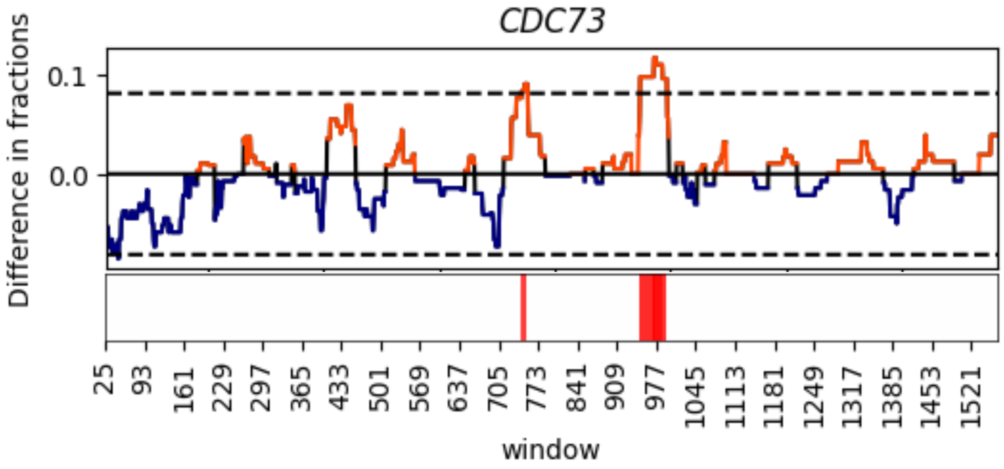


COSMIC

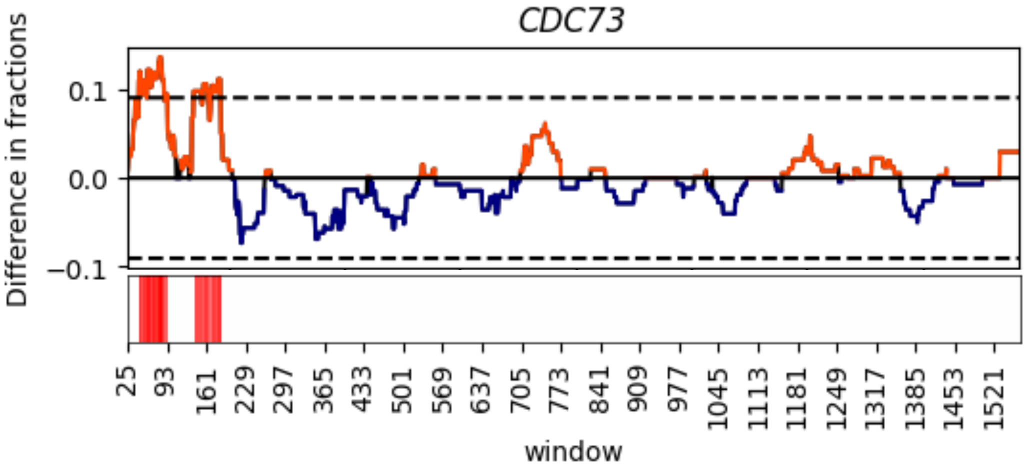

cBioPortal

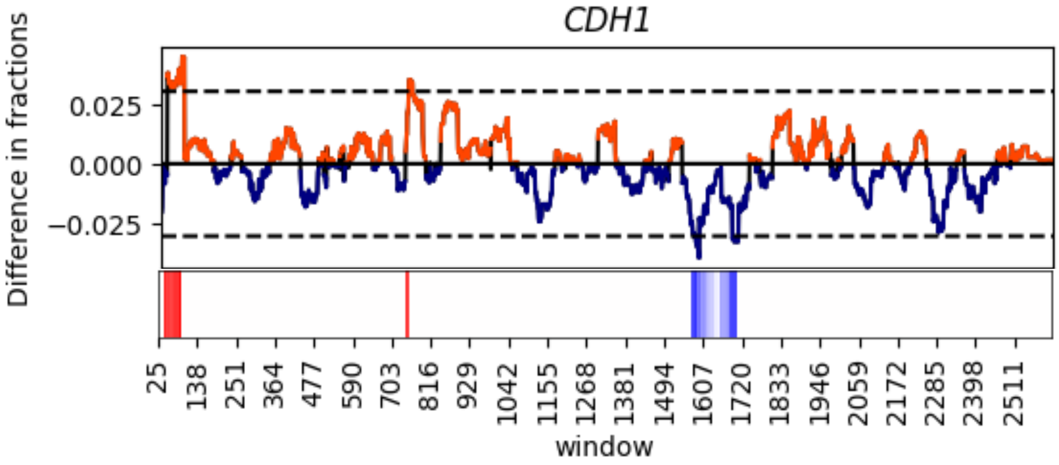


COSMIC

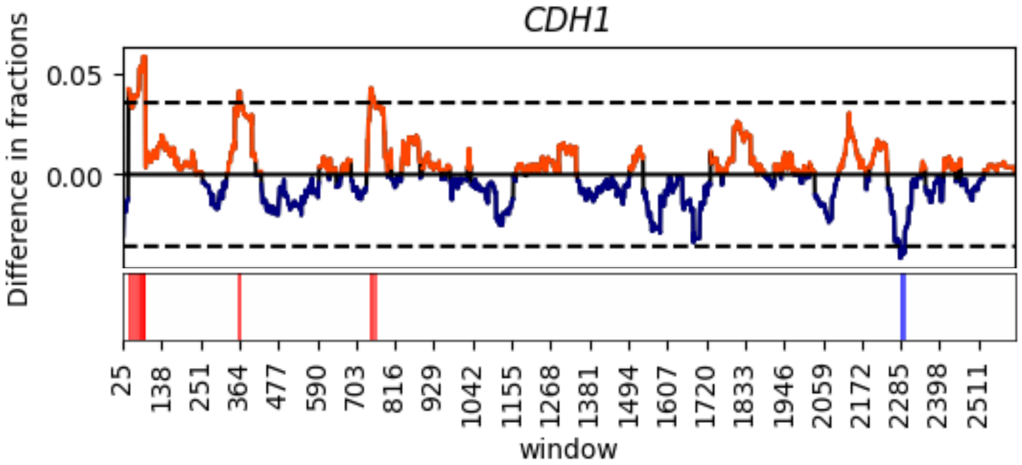

cBioPortal

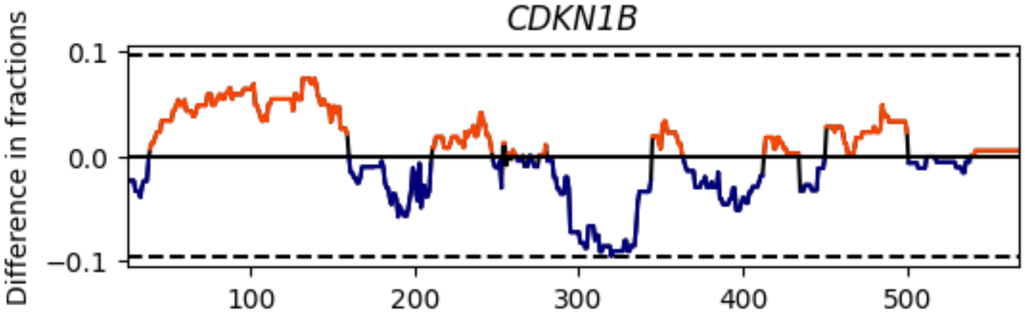


COSMIC

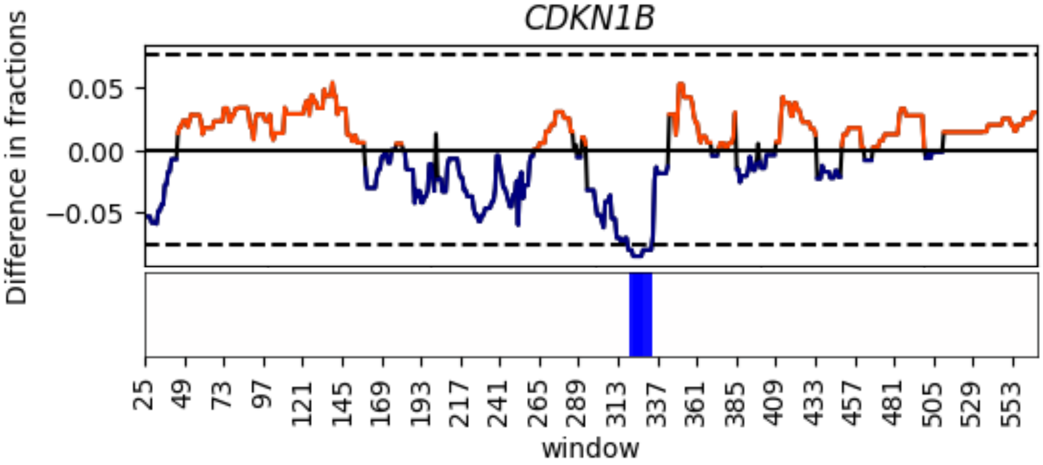

cBioPortal

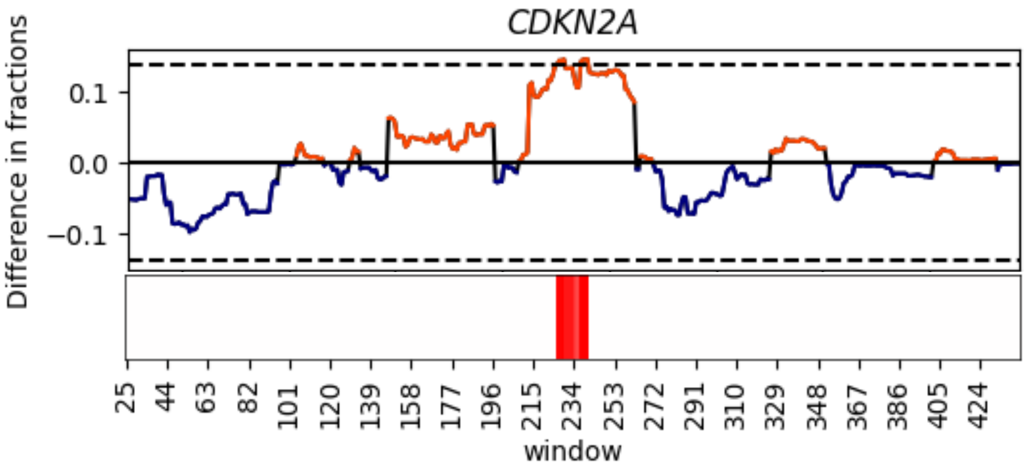


COSMIC

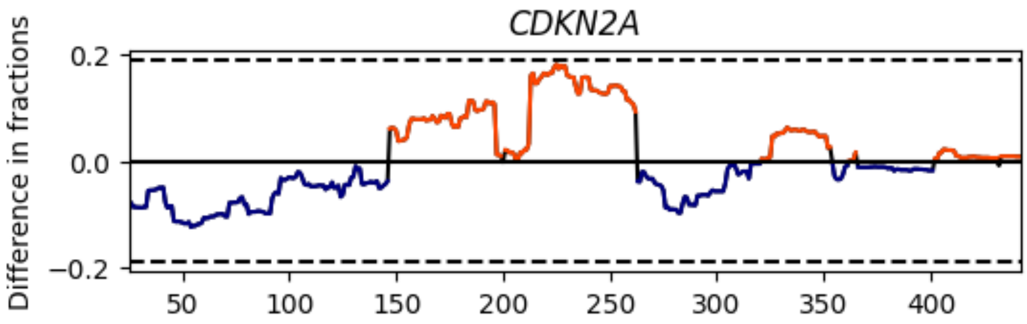

cBioPortal

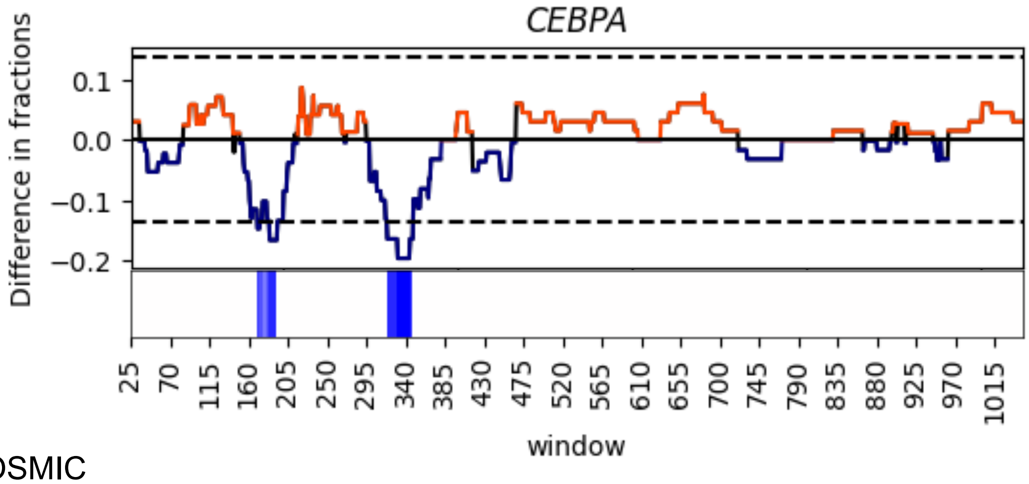


COSMIC

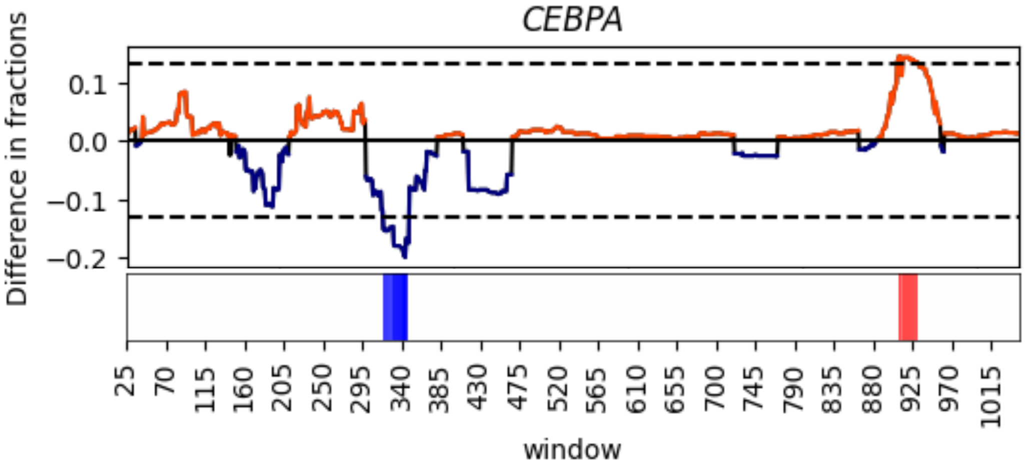

cBioPortal

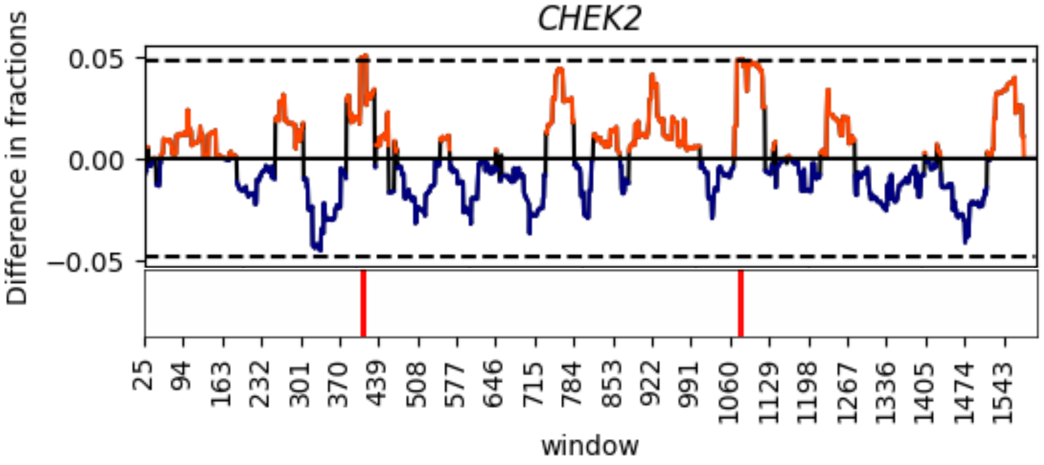


COSMIC

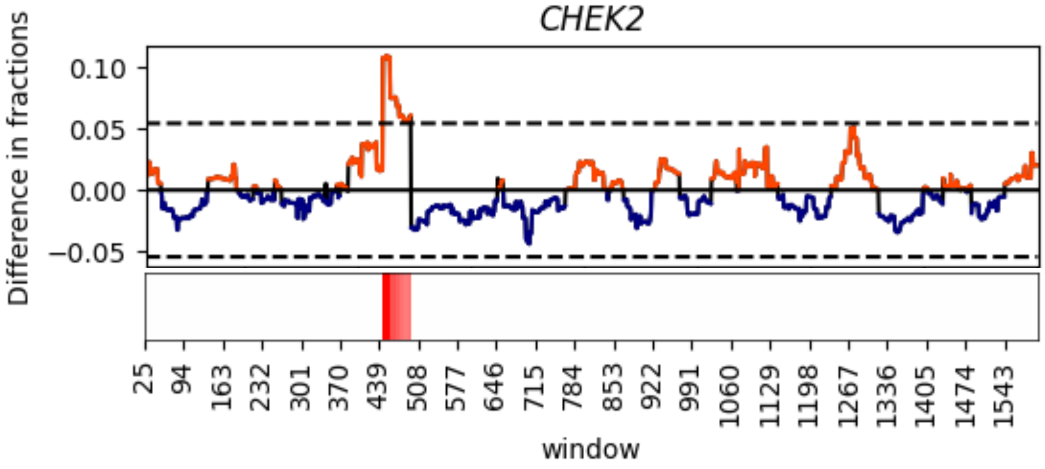

cBioPortal

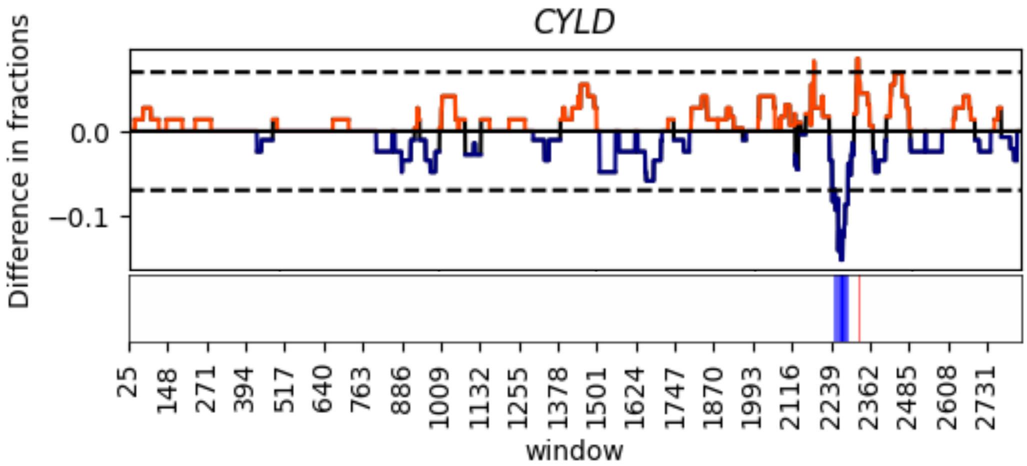


COSMIC

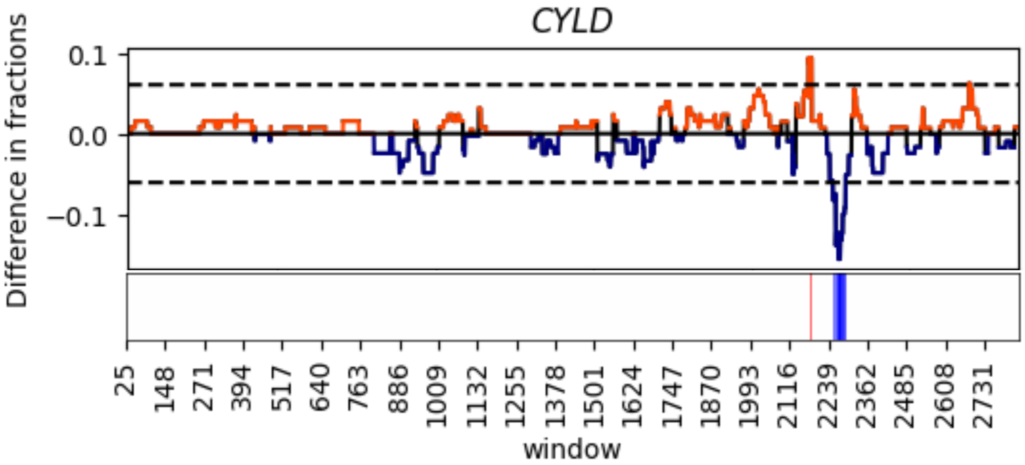

cBioPortal

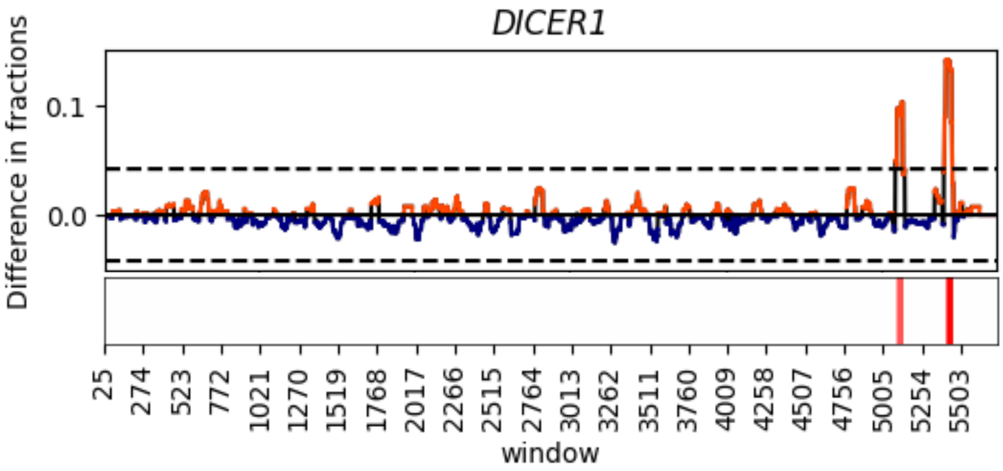


COSMIC

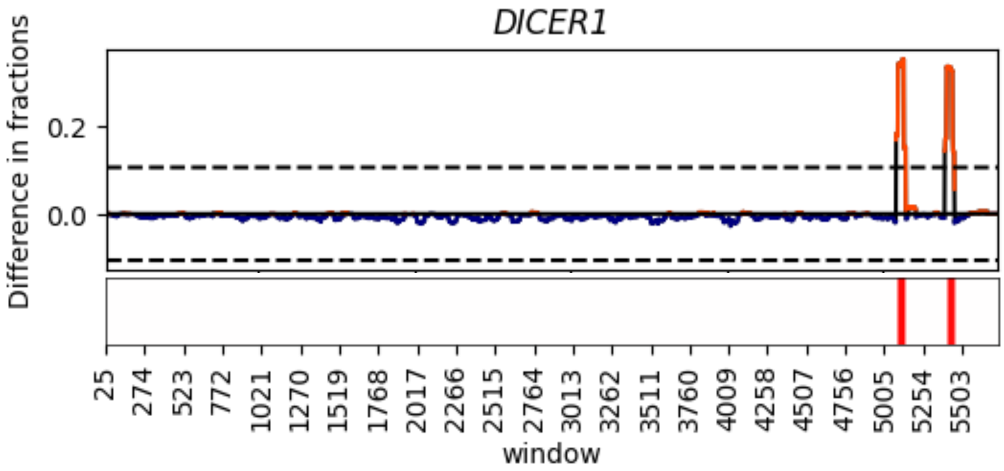

cBioPortal

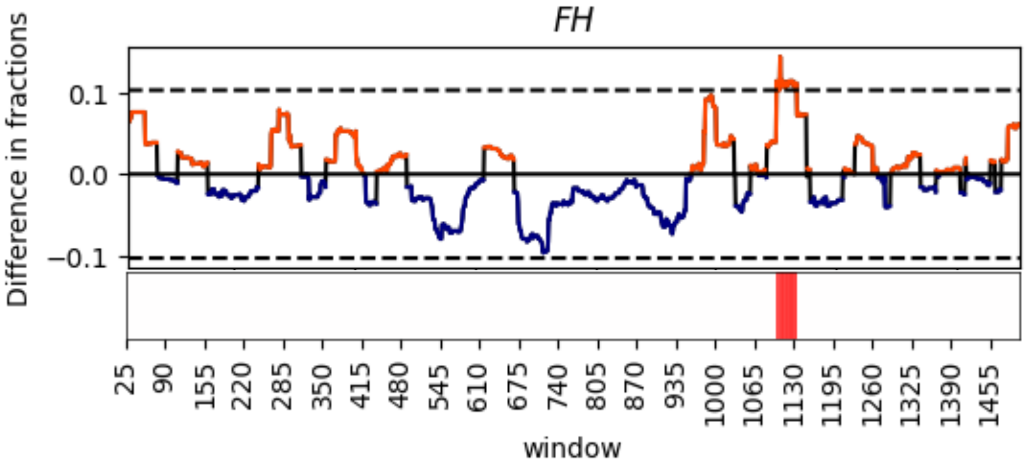


COSMIC

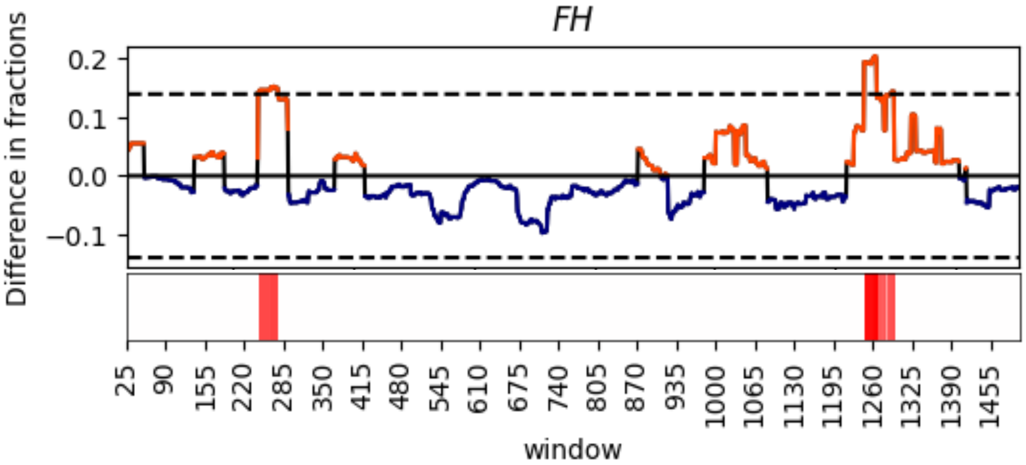

cBioPortal

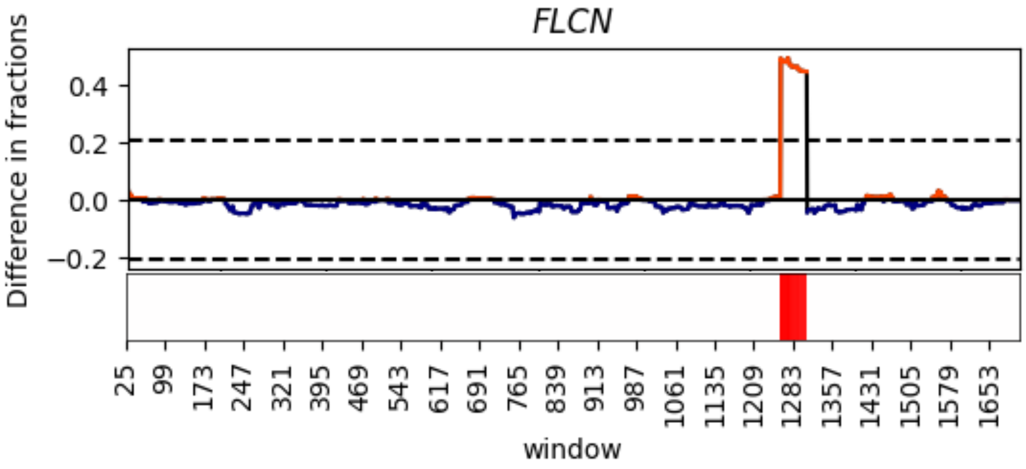


COSMIC

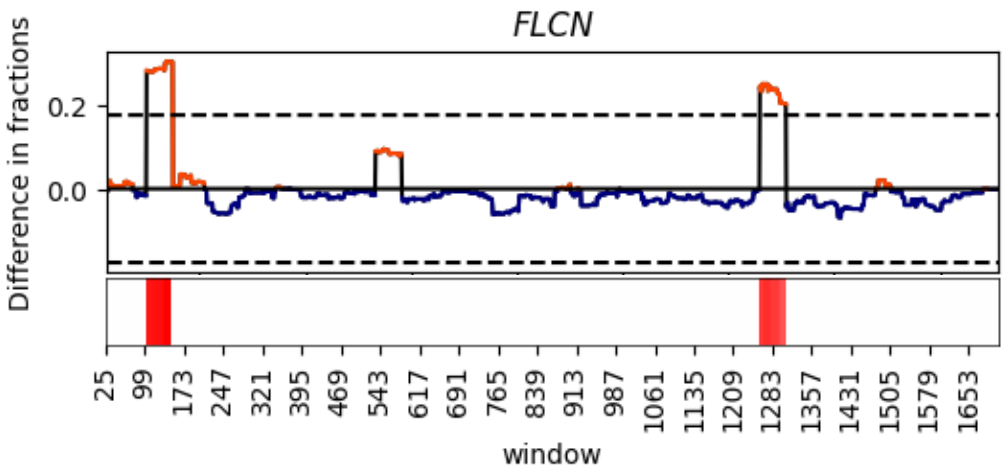

cBioPortal

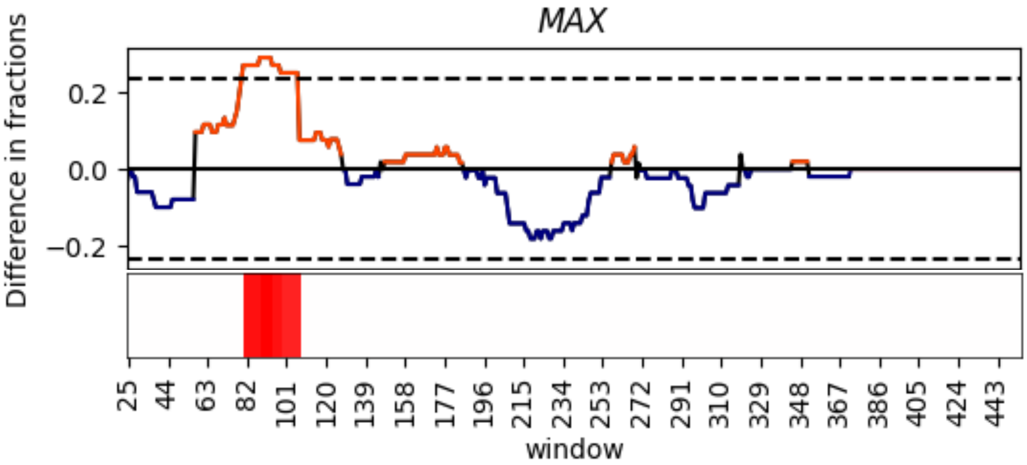


COSMIC

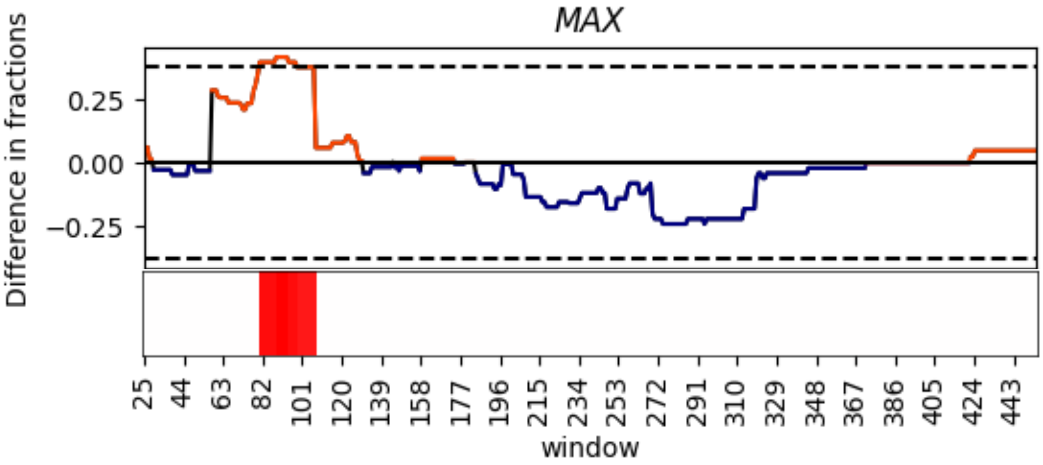

cBioPortal

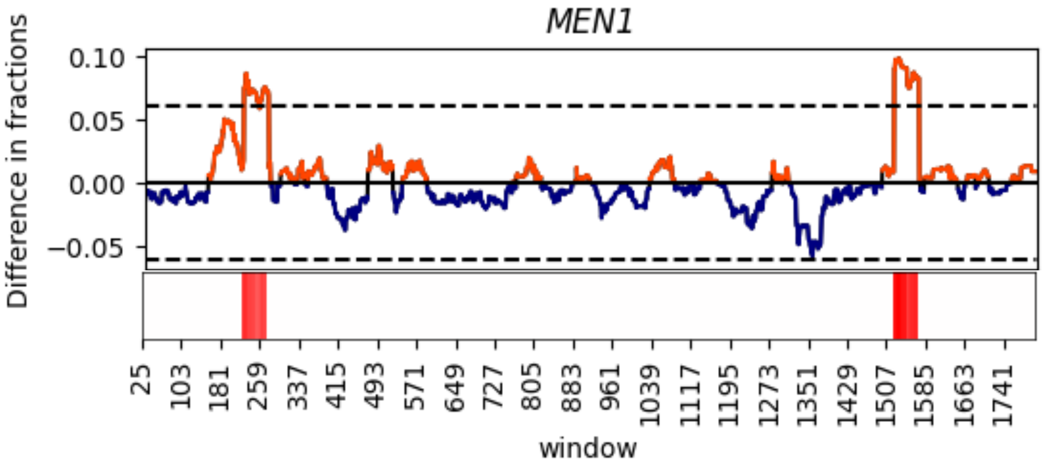


COSMIC

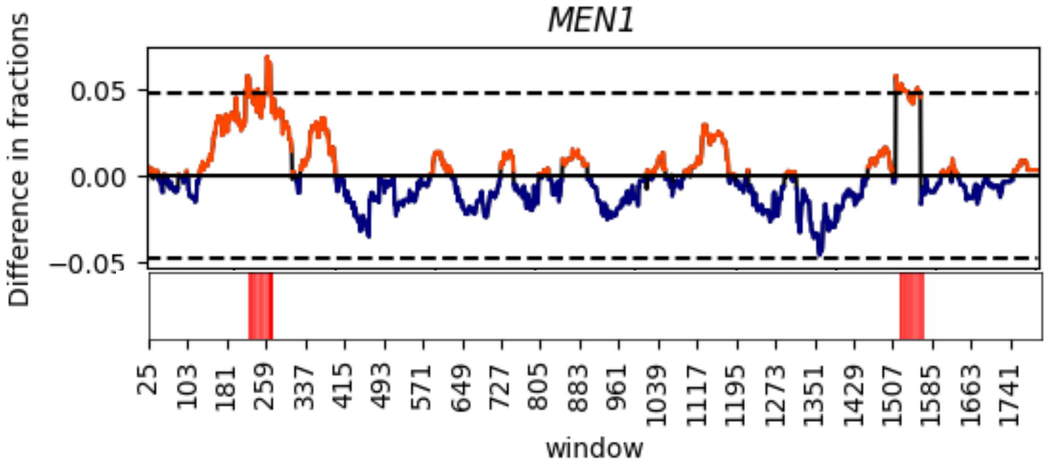

cBioPortal

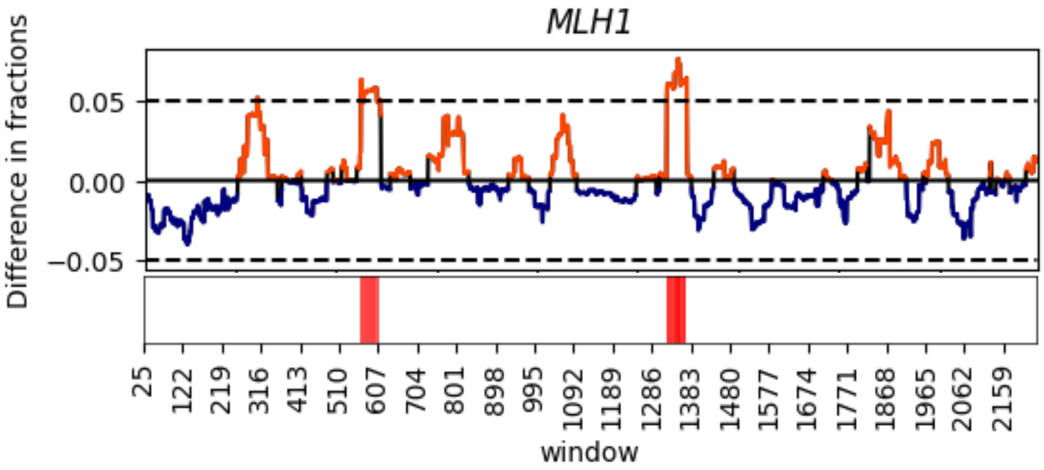


COSMIC

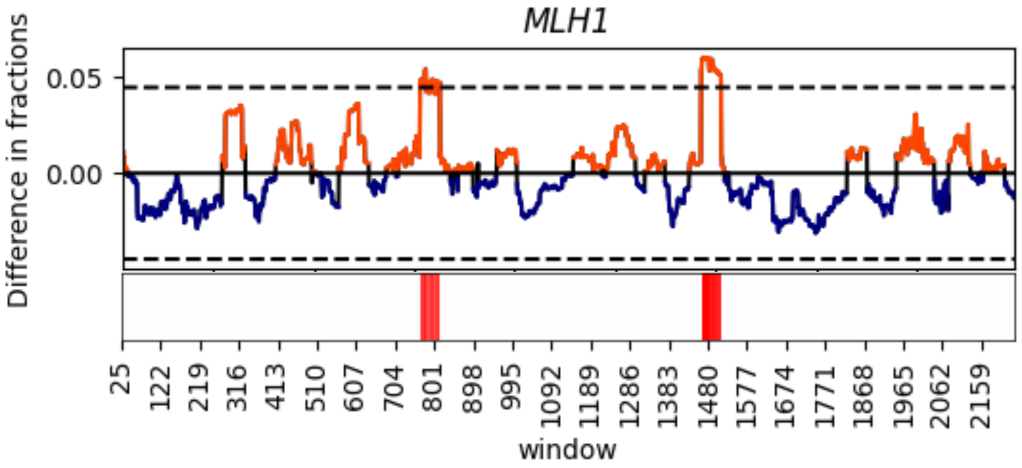

cBioPortal

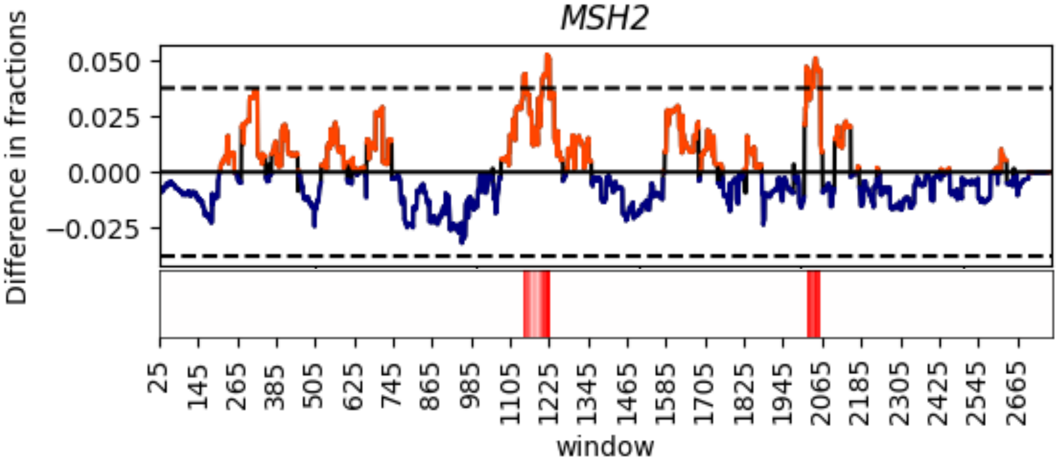


COSMIC

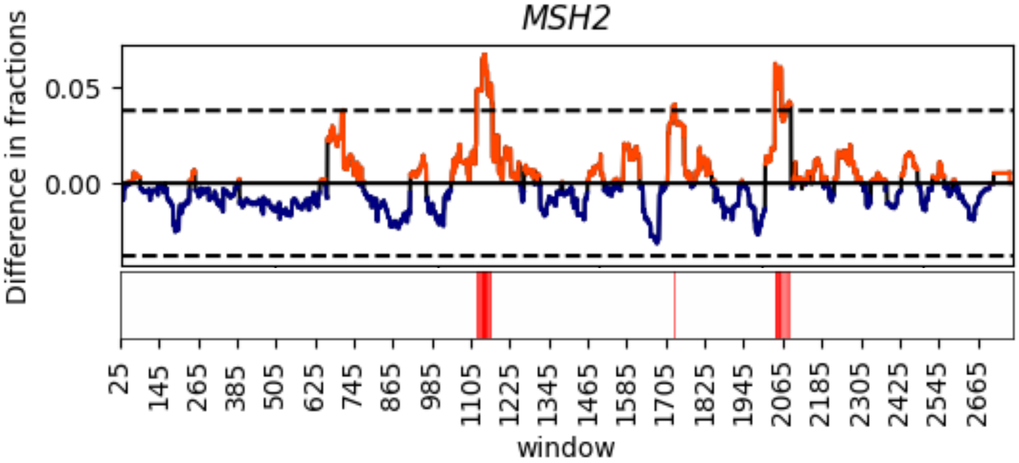

cBioPortal

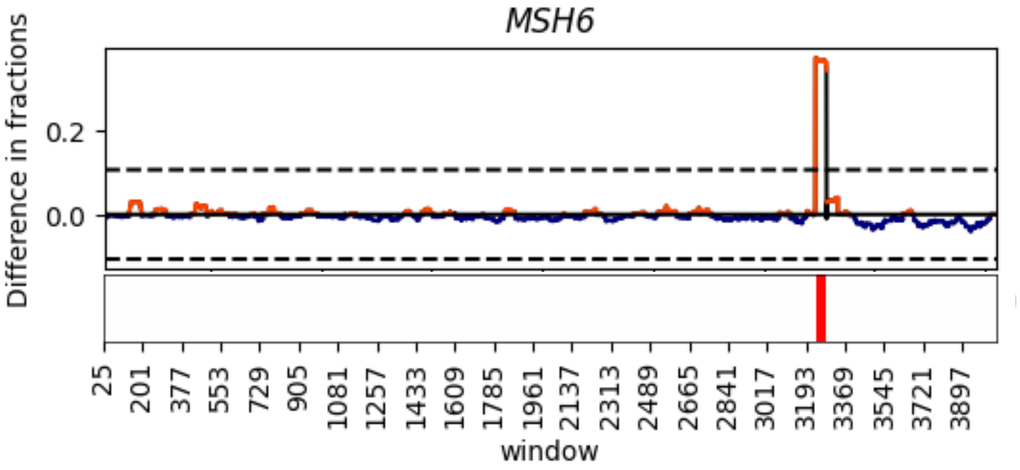


COSMIC

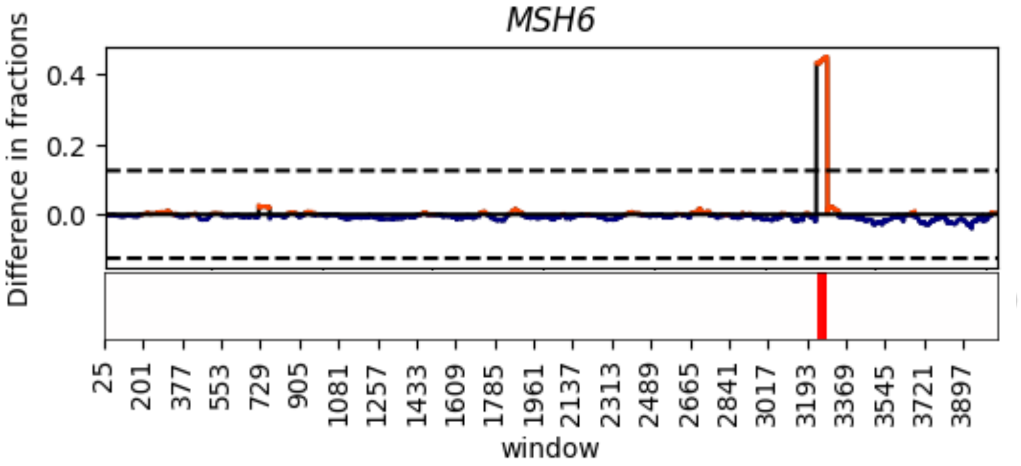

cBioPortal

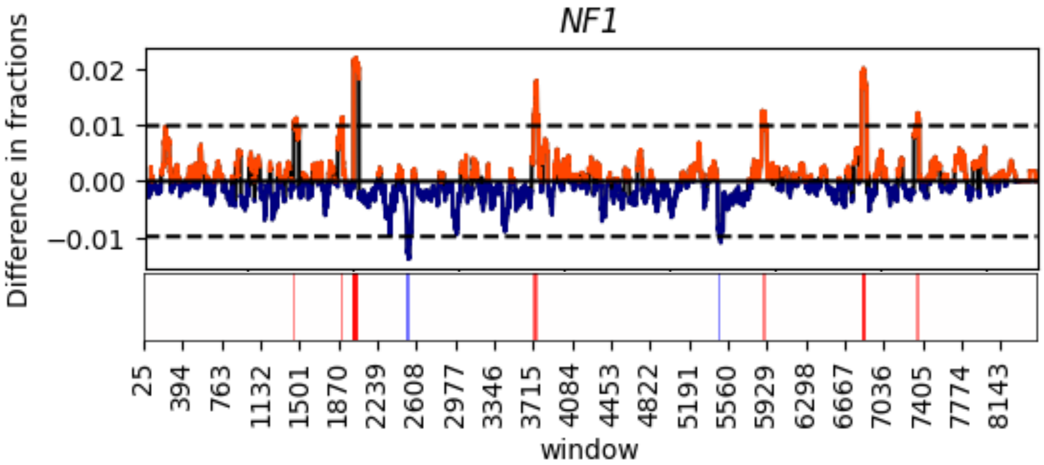


COSMIC

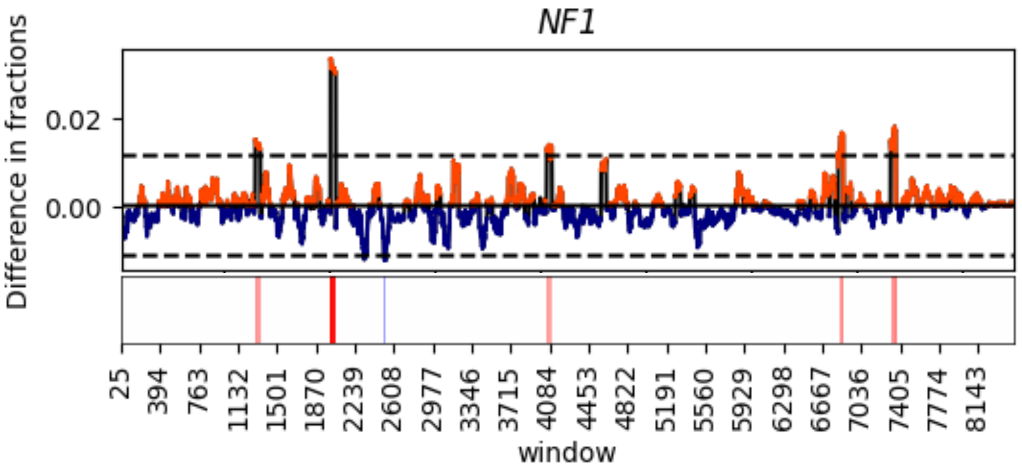

cBioPortal

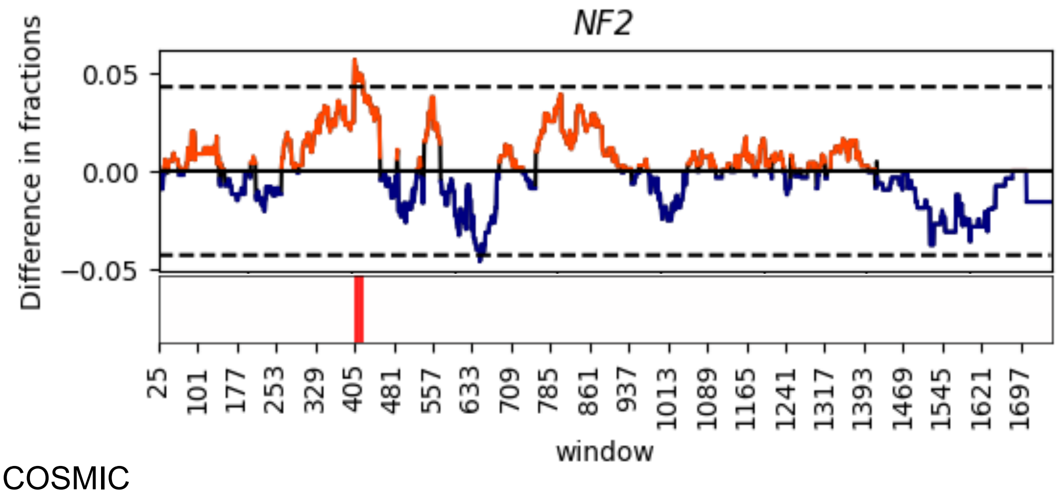


COSMIC

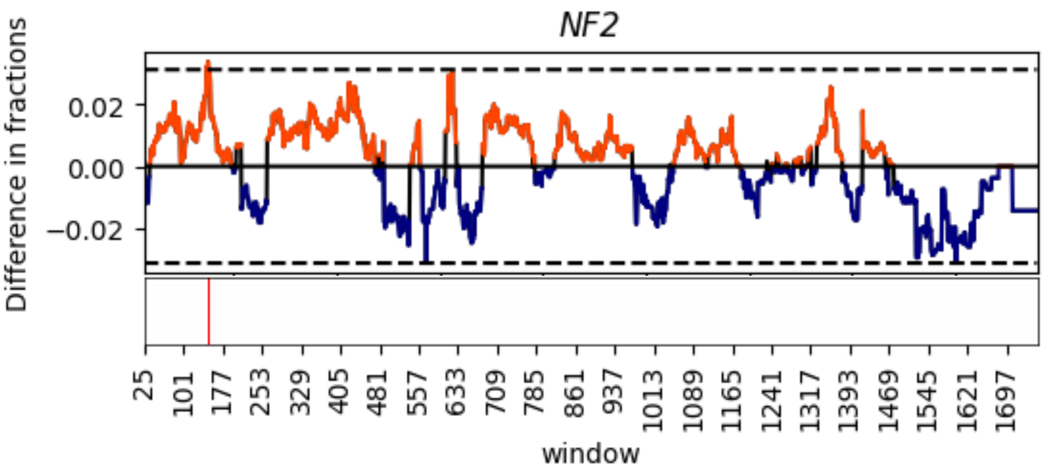

cBioPortal

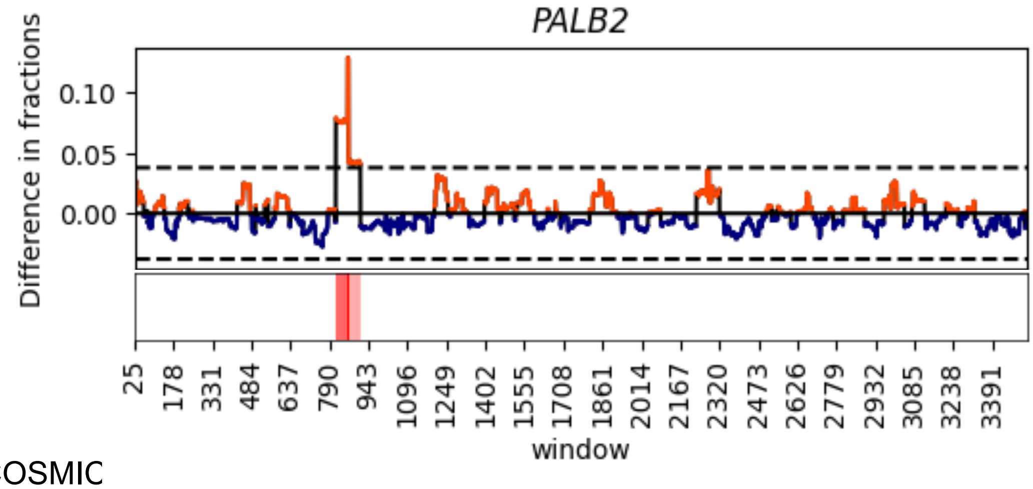


COSMIC

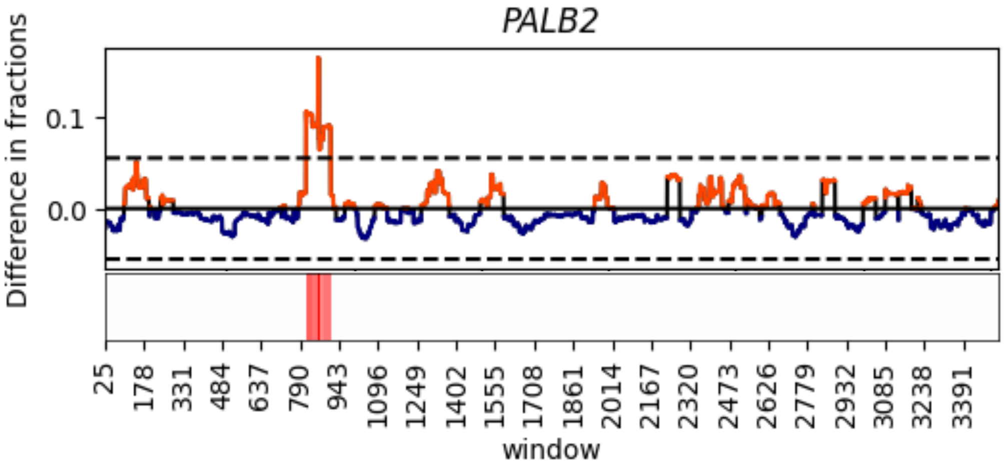

cBioPortal

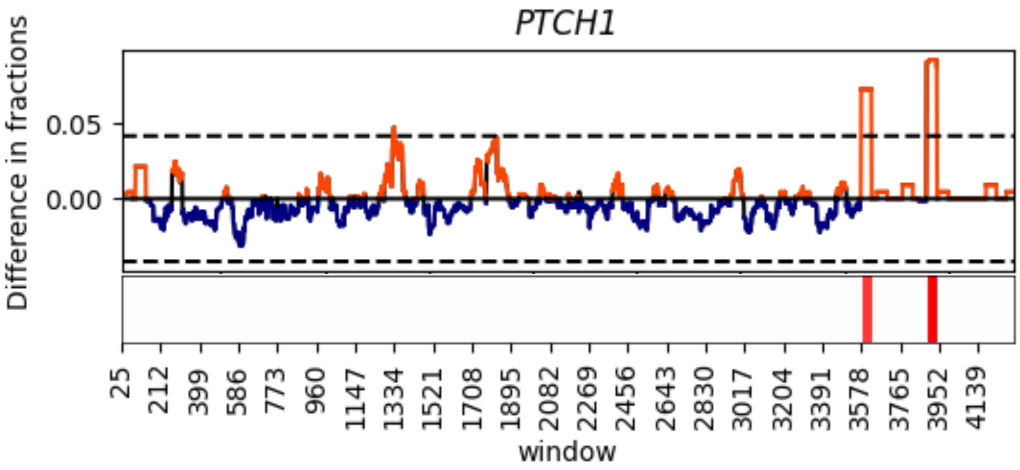


COSMIC

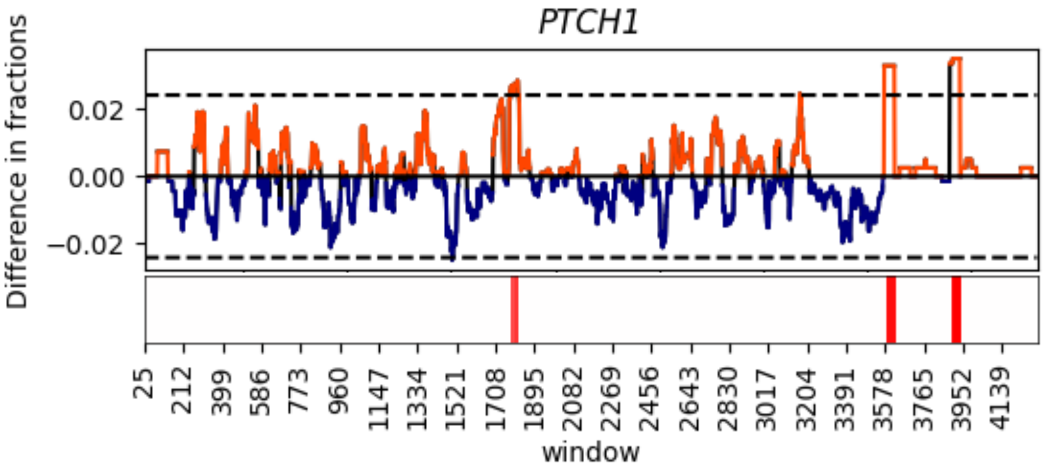

cBioPortal

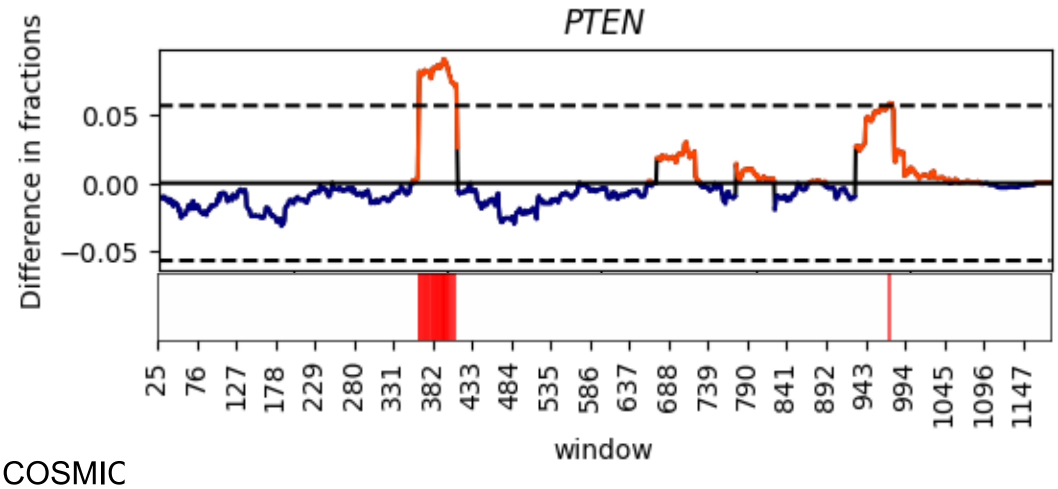


COSMIC

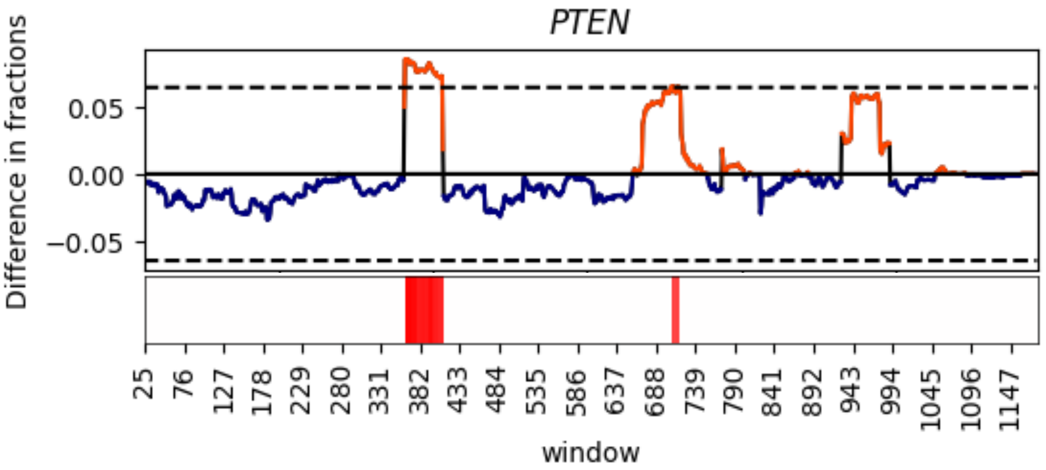

cBioPortal

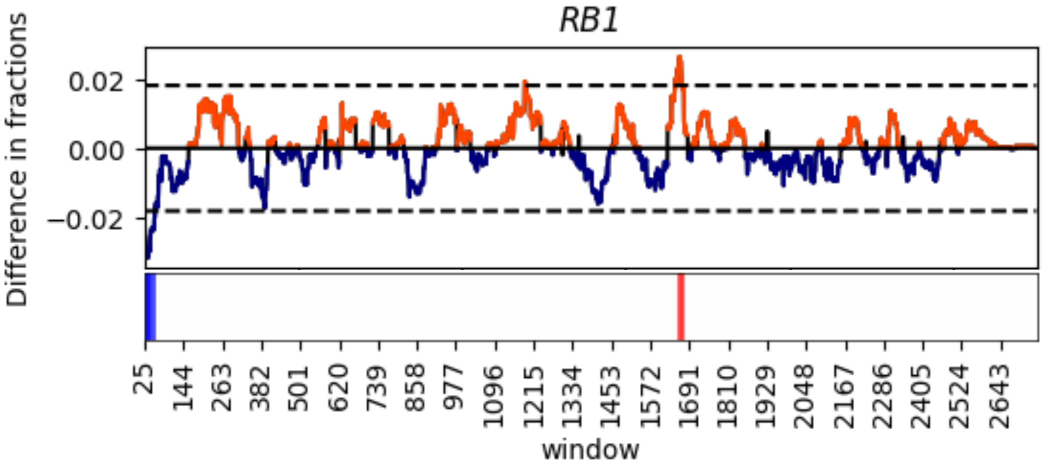


COSMIC

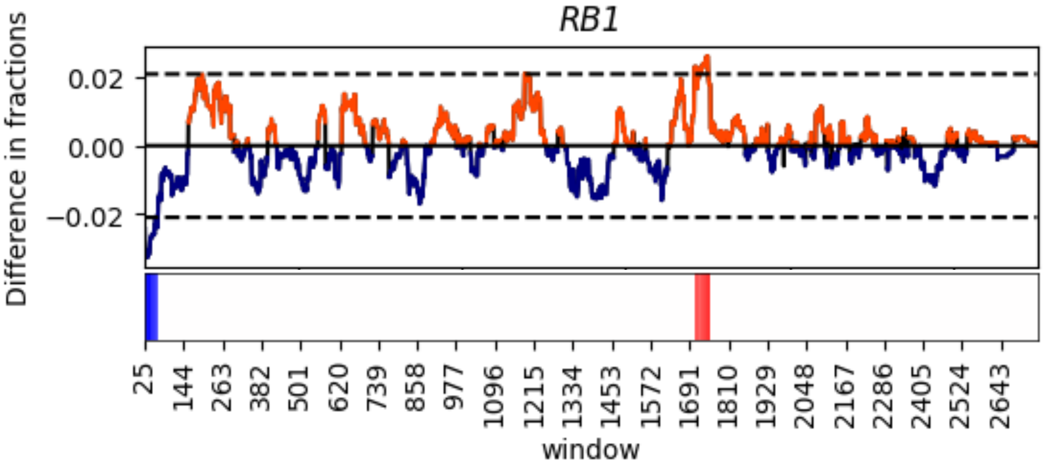

cBioPortal

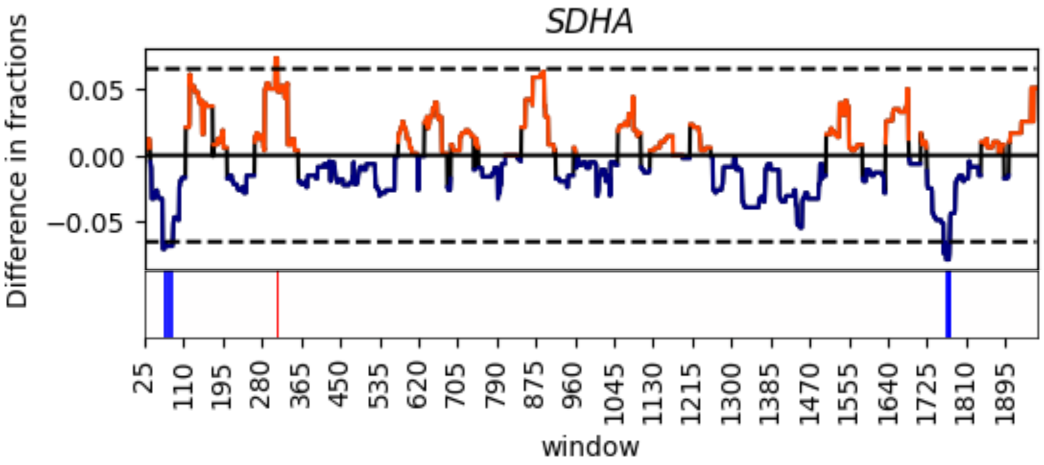


COSMIC

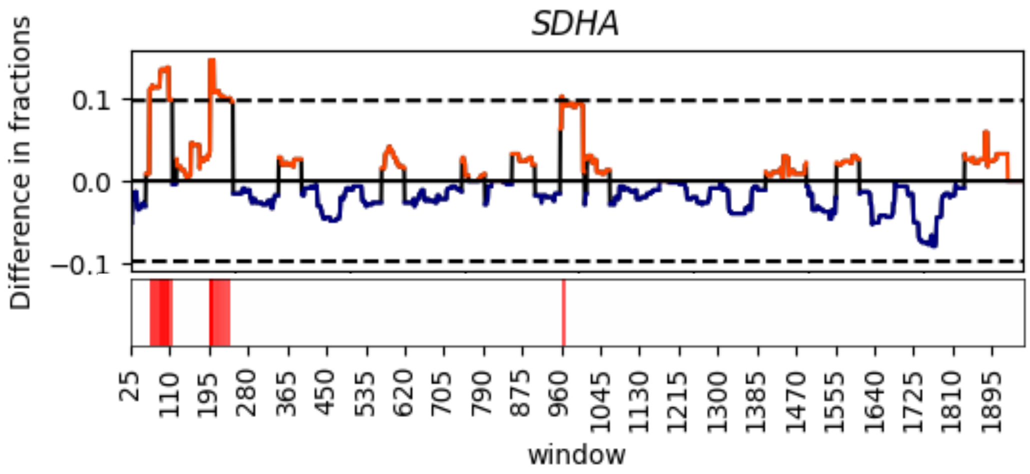

cBioPortal

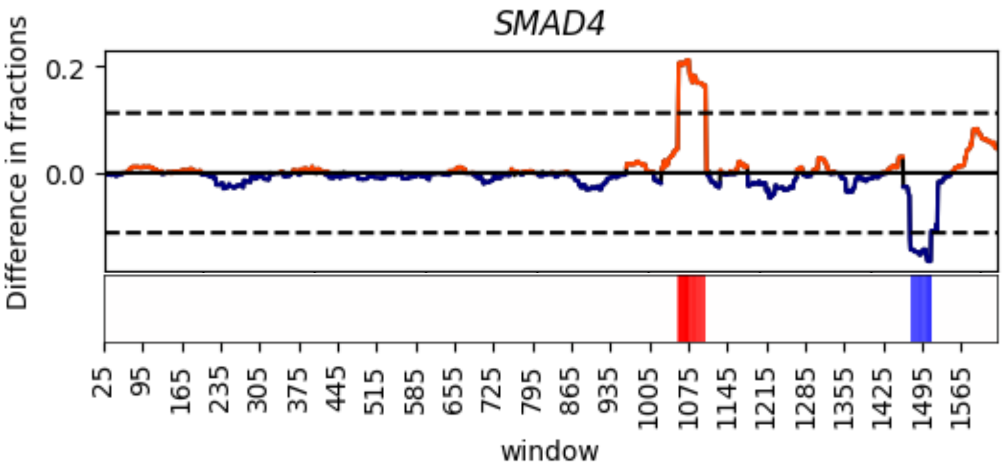


COSMIC

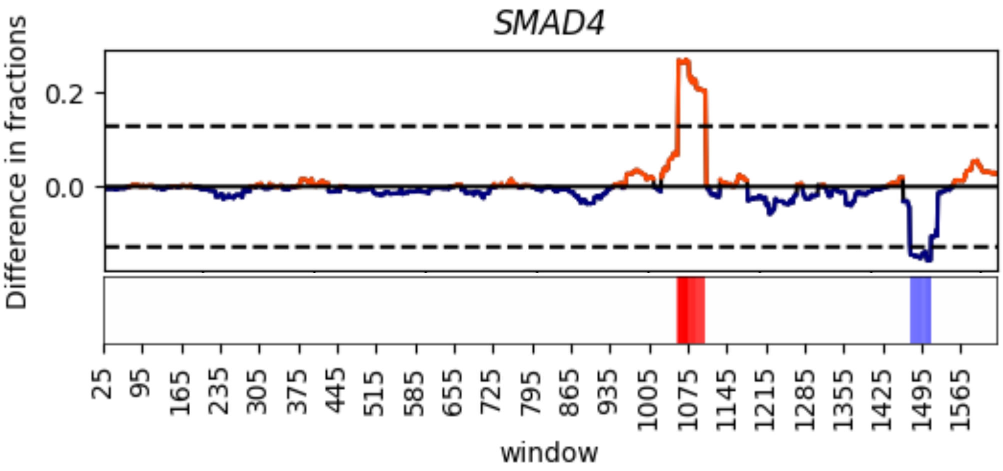

cBioPortal

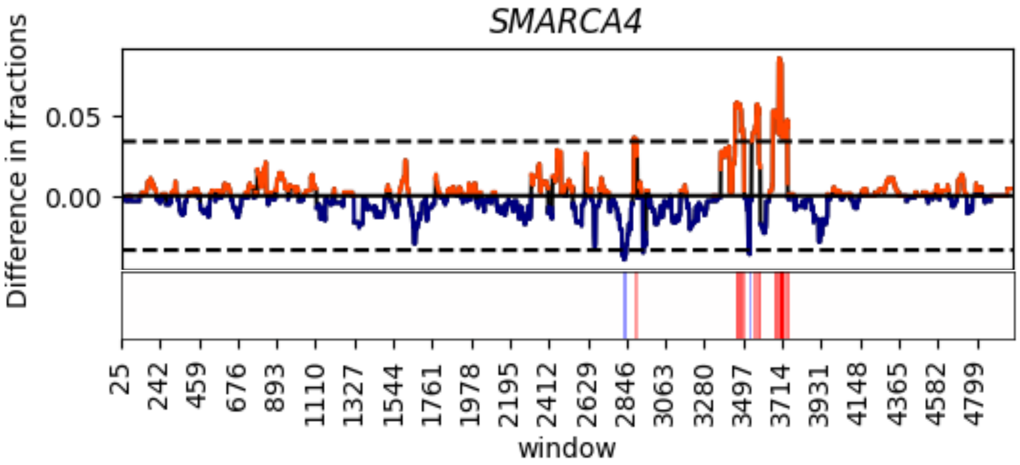


COSMIC

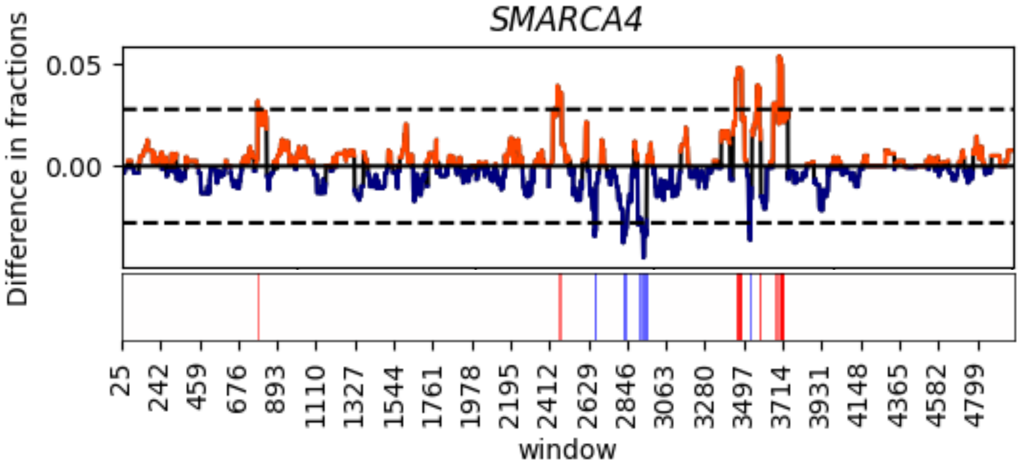

cBioPortal

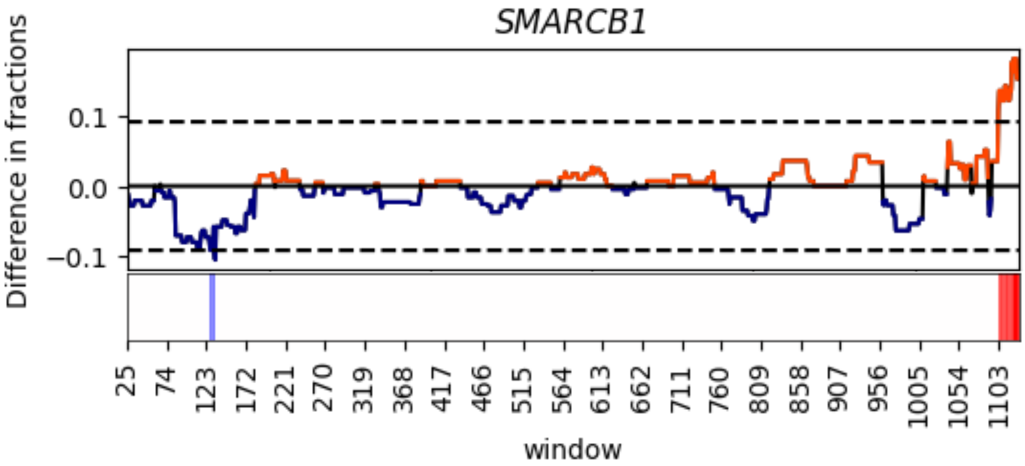


COSMIC

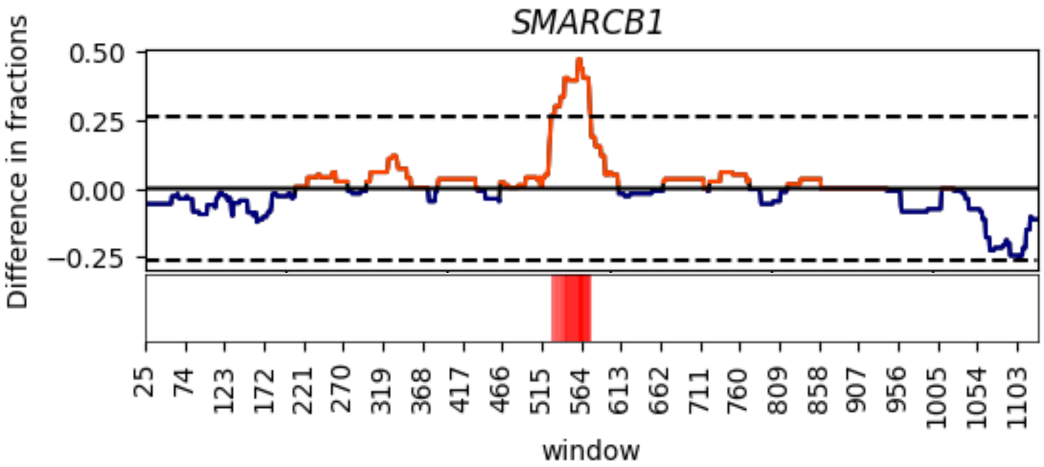

cBioPortal

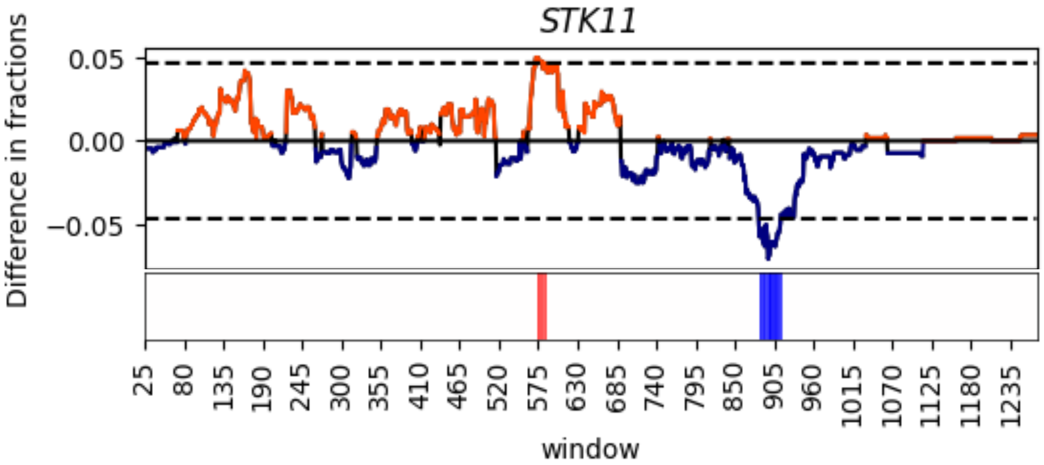


COSMIC

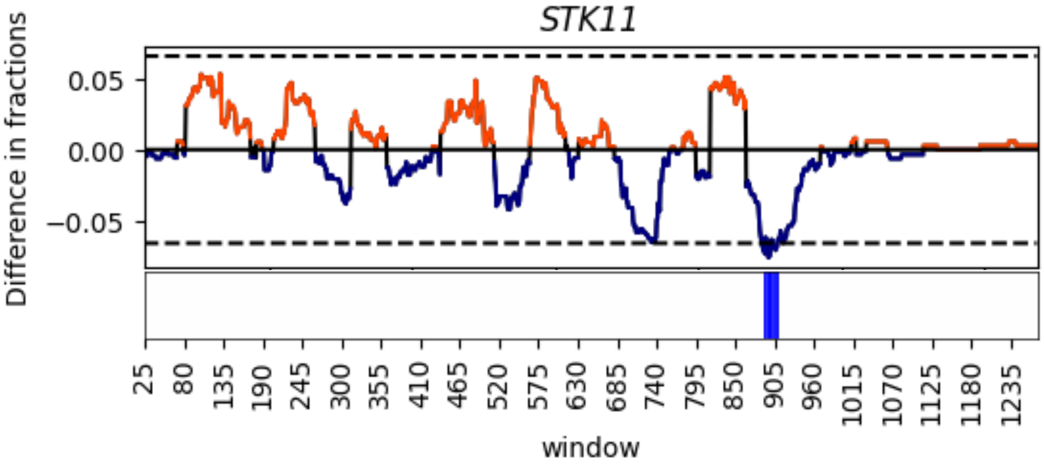

cBioPortal

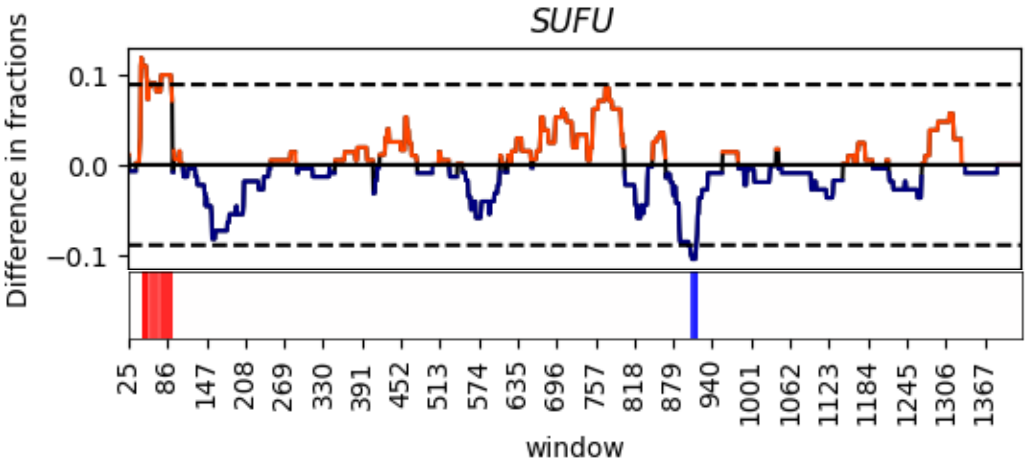


COSMIC

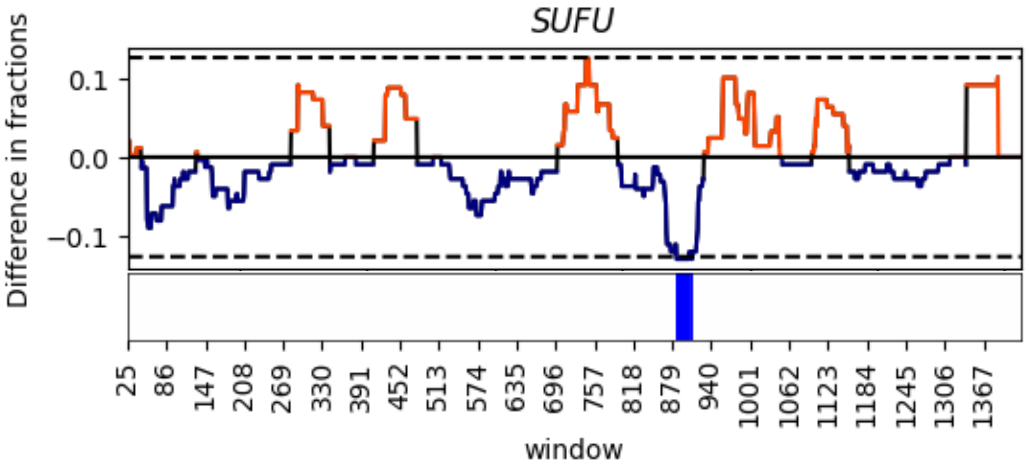

cBioPortal

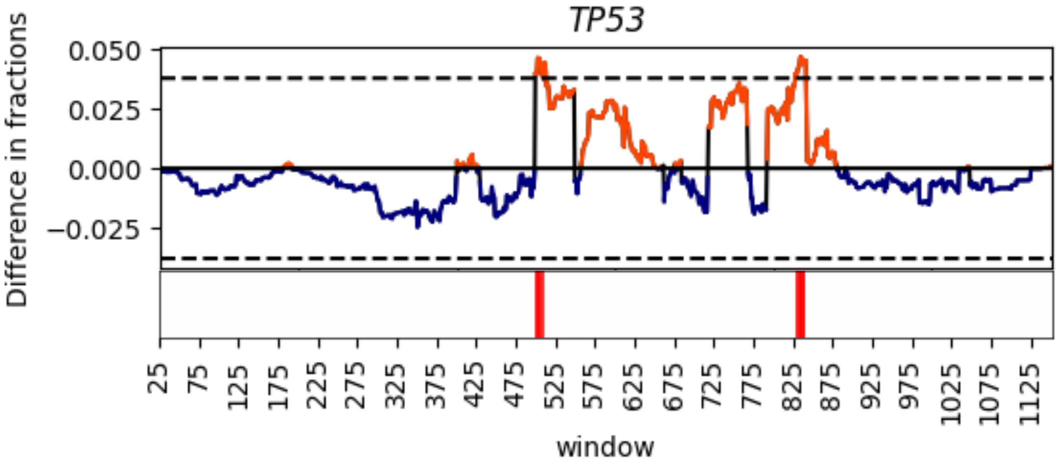


COSMIC

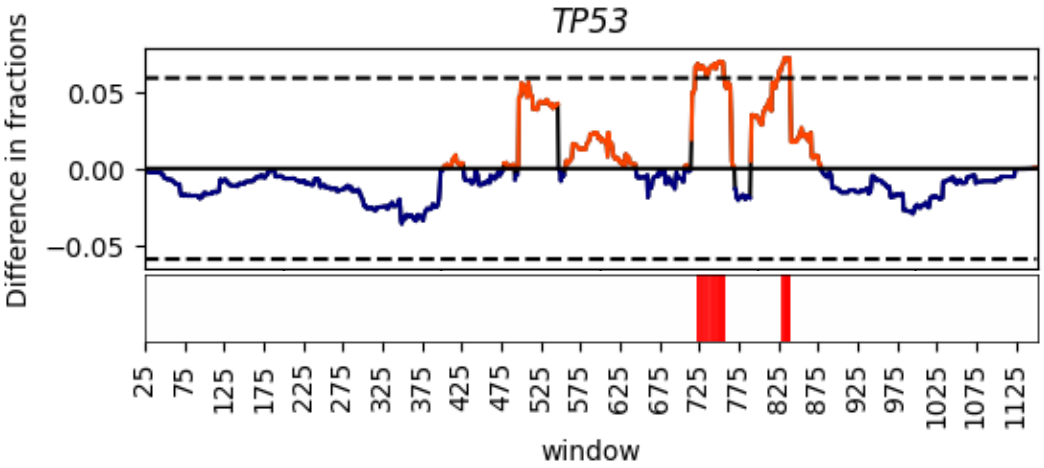

cBioPortal

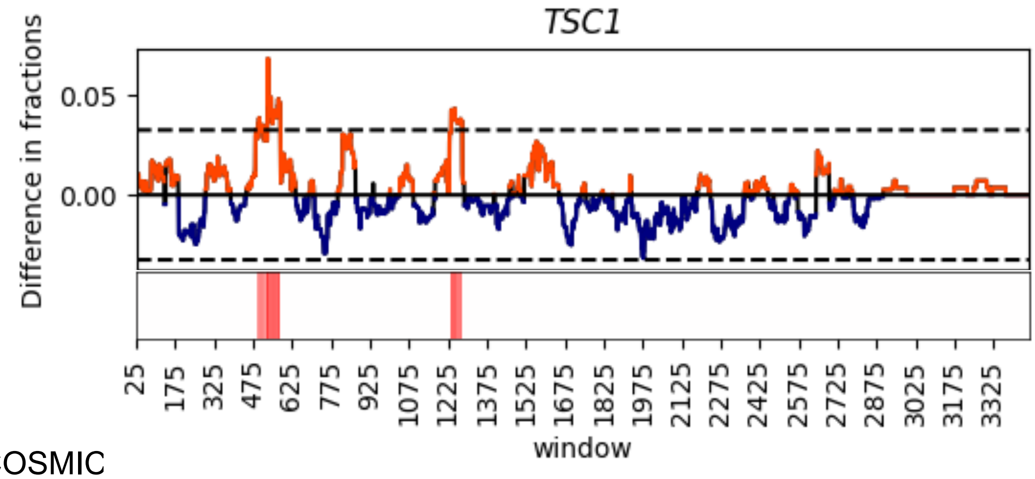


COSMIC

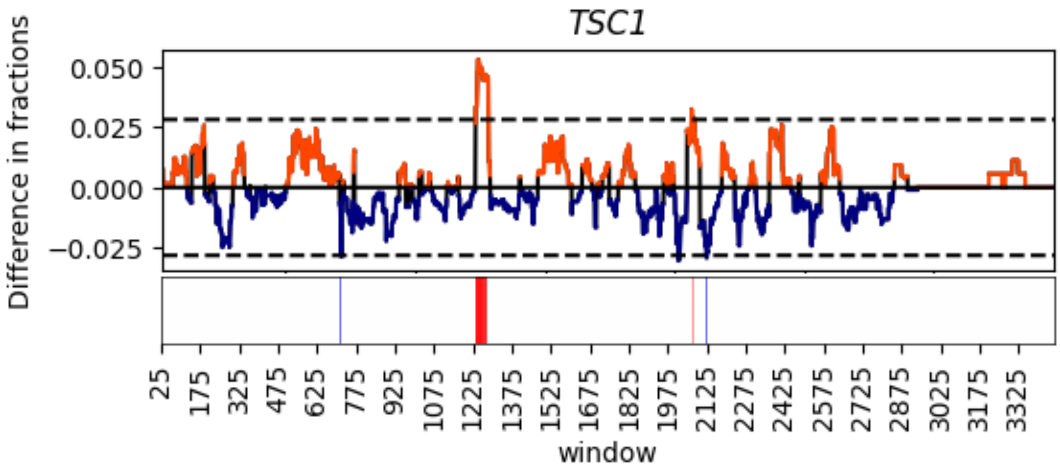

cBioPortal

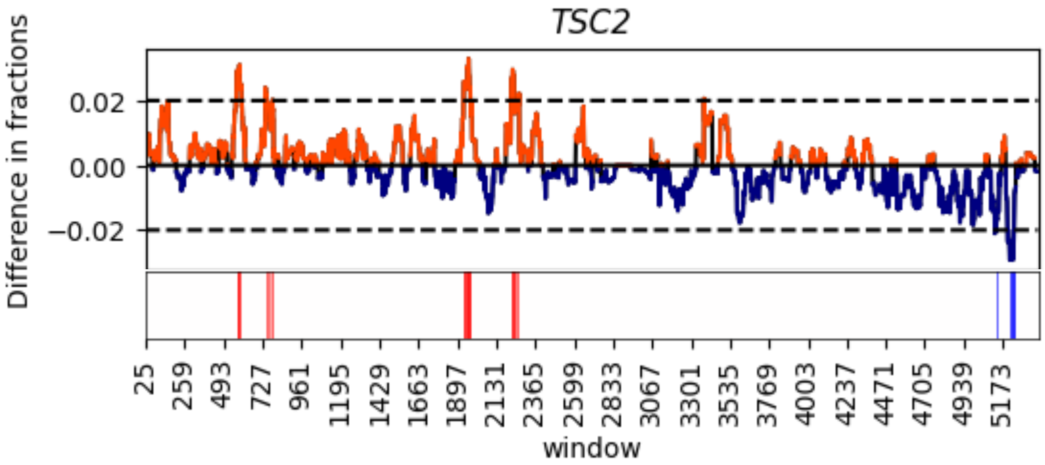


COSMIC

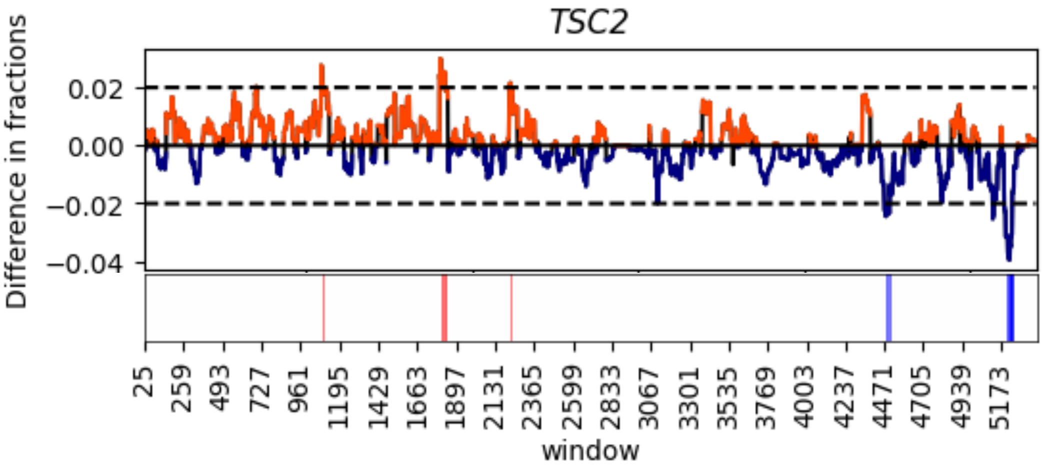

cBioPortal

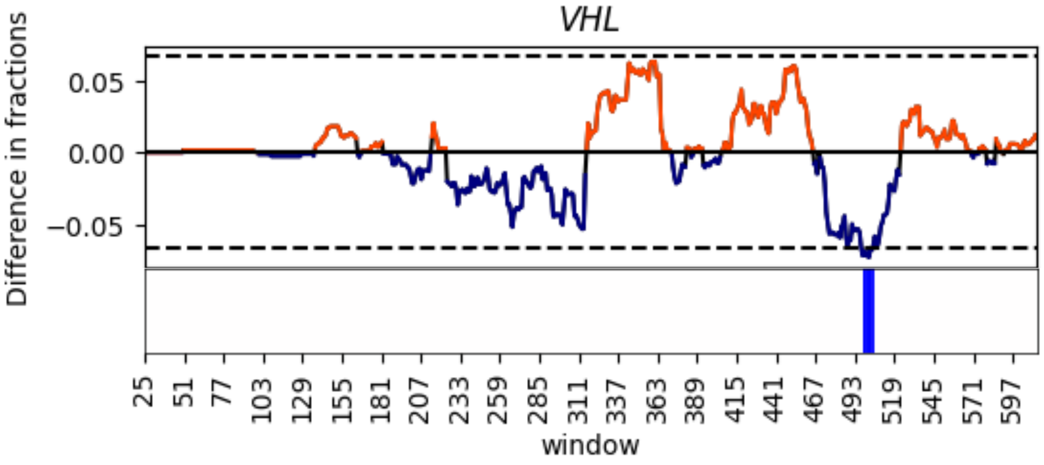


COSMIC

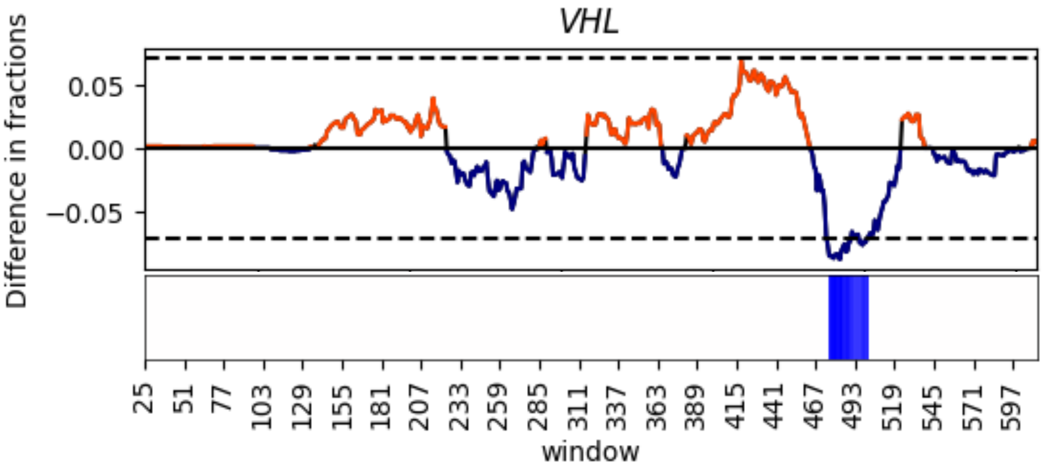

cBioPortal

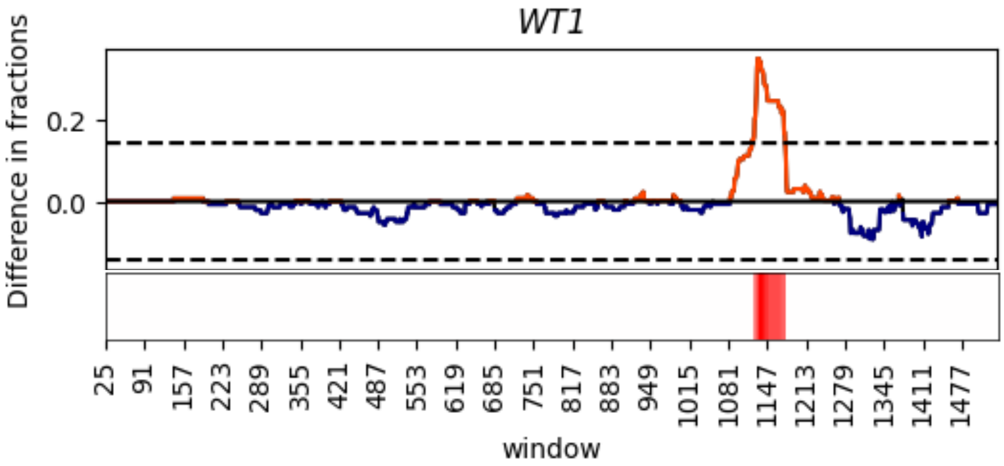


COSMIC

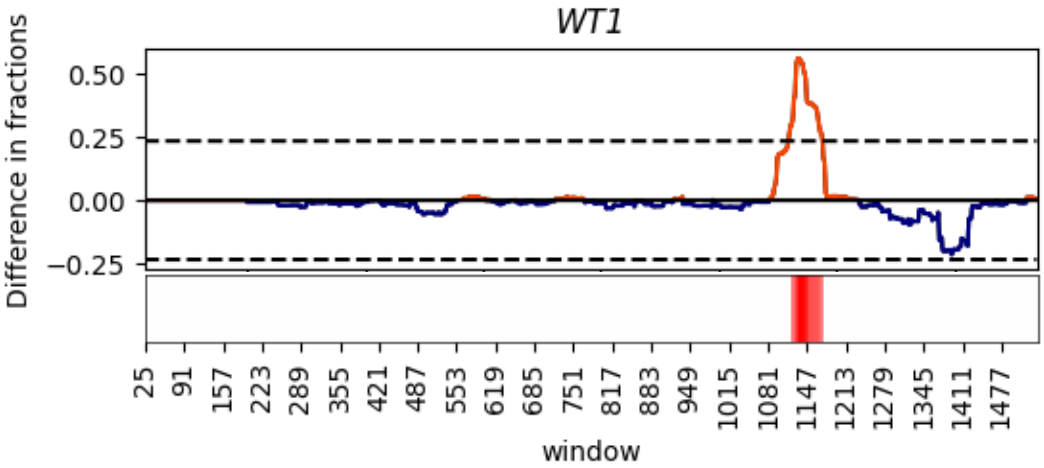


**Figure S7. Moving difference in fraction curves capturing preferential clustering of P/LP germline or O/LO somatic events in 40 TSGs.** For each gene, a sliding window, sized 50bp, moves along the cDNA 1bp at a time comparing O/LO somatic data from cBioPortal (top) or COSMIC (bottom) to P/LP germline data from ClinVar to capture preferentially clustered regions >±2.5SD from 0. A one-dimensional heatmap is included below the difference in fraction curve to demarcate germline (blue) or somatic (red) clusters along the cDNA (starting at the 25th coding position, the mid-point of the first 50bp window and ending at the mid-point of the last 50bp window). Significant clusters may not be depicted in the one-dimensional heatmap if they are smaller than the minimum length represented on the x-axis scale for that gene. These are included in a list of all significant clusters that can be found in Table S10 (cBioPortal) and Table S11 (COSMIC). P/LP, pathogenic/likely pathogenic; O/LO, oncogenic/likely oncogenic; TSGs, tumor suppressor genes; bp, base pair; COSMIC, Catalogue of Somatic Mutations in Cancer; SD, standard deviation.

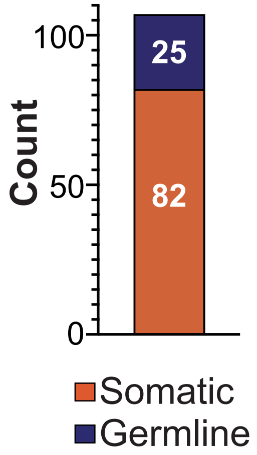


**Figure S8. Count of germline and somatic clusters when comparing O/LO somatic data from COSMIC to P/LP germline data from ClinVar by location.**
P/LP, pathogenic/likely pathogenic; O/LO, oncogenic/likely oncogenic; COSMIC, Catalogue of Somatic Mutations in Cancer.

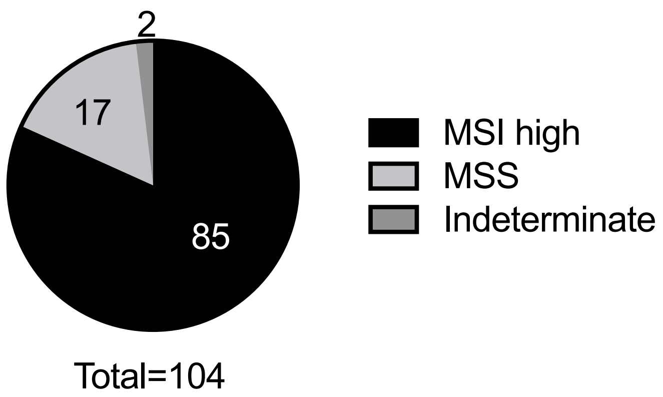


**Figure S9. Distribution of MSI status across tumor samples in cBioPortal.**
MSI, microsatellite instability; MSS, microsatellite stable.

A

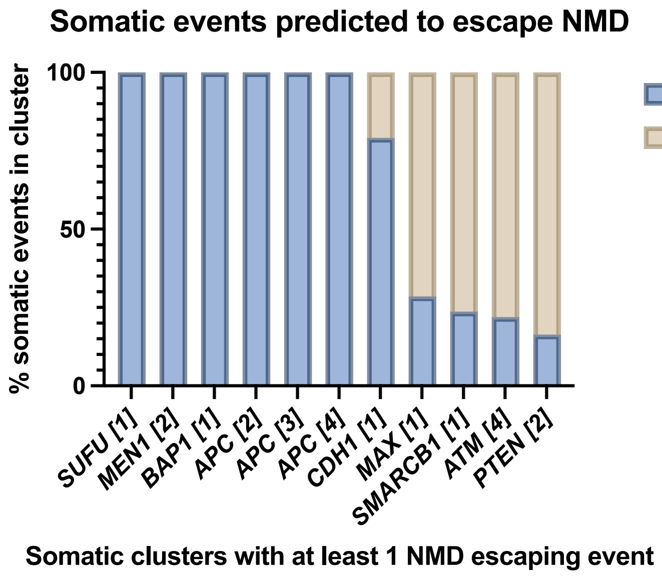


B

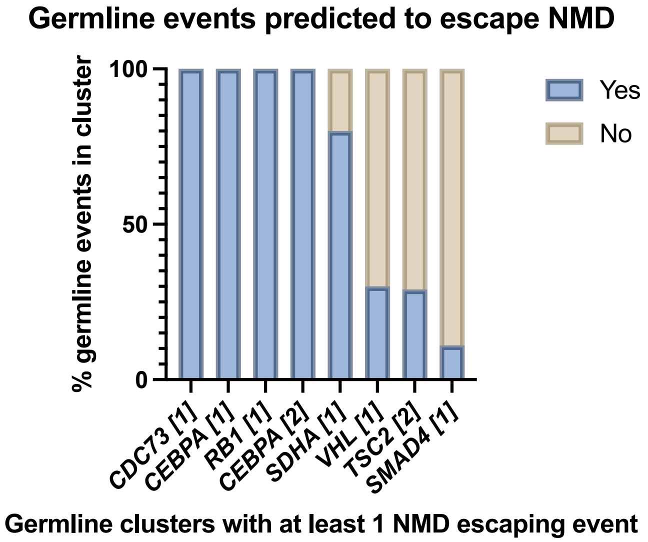


**Figure S10. Preferential clusters with NMD escaping variants.**
(A) Proportion of O/LO events from cBioPortal in each somatic cluster that are predicted to escape NMD. Only those somatic clusters with at least one O/LO event predicted to escape NMD are included.
(B) Proportion of P/LP events from ClinVar in each germline cluster that are predicted to escape NMD. Only those germline clusters with at least one P/LP event predicted to escape NMD are included.
P/LP, pathogenic/likely pathogenic; O/LO, oncogenic/likely oncogenic; NMD; nonsense mediated decay.
